# Carbohydrate sulfation as a mechanism for fine-tuning Siglec ligands

**DOI:** 10.1101/2021.06.27.450109

**Authors:** Jaesoo Jung, Jhon R. Enterina, Duong T. Bui, Fahima Mozaneh, Po-Han Lin, Nitin, Chu-Wei Kuo, Emily Rodrigues, Abhishek Bhattacherjee, Parisa Raeisimakiani, Gour C. Daskhan, Chris D. St. Laurent, Kay-Hooi Khoo, Lara K. Mahal, Wesley F. Zandberg, Xuefei Huang, John S. Klassen, Matthew S. Macauley

## Abstract

The immunomodulatory family of Siglecs recognize sialic acid-containing glycans as ‘*self*’, which is exploited in cancer for immune-evasion. The biochemical nature of Siglec ligands remains incompletely understood with emerging evidence suggesting the importance of carbohydrate sulfation. Here, we investigate how specific sulfate modifications affect Siglec ligands by overexpressing eight carbohydrate sulfotransferases (CHSTs) in five cell lines. Overexpression of three CHSTs (*CHST1*, *CHST2,* or *CHST4*) significantly enhances the binding of numerous Siglecs. Unexpectedly, two other CHSTs (*Gal3ST2* and *Gal3ST3*) diminish Siglec binding, suggesting a new mode to modulate Siglec ligands via sulfation. Results are cell type dependent, indicating that the context in which sulfated glycans are presented is important. Moreover, pharmacological blockade of *N*- and *O*-glycan maturation reveals a cell type-specific pattern of importance for either class of glycan. Production of a highly homogenous CD33 (Siglec-3) fragment enabled a mass spectrometry-based binding assay to determine 10-fold and 3-fold enhanced affinity for Neu5Acα2-3(6-*O*-sulfo)Galβ1-4GlcNAc and Neu5Acα2-3Galβ1-4(6-*O*- sulfo)GlcNAc, respectively, over Neu5Acα2-3Galβ1-4GlcNAc. CD33 showed significant additivity in affinity (36-fold) for the disulfated ligand, Neu5Acα2-3(6-*O*-sulfo)Galβ1-4(6-*O*-sulfo)GlcNAc. Moreover, overexpression of both *CHST1* and *CHST2* in cells greatly enhanced the binding of several Siglecs, including CD33. Finally, we reveal that *CHST1* is upregulated in numerous cancers, correlating with poorer survival rates and sodium chlorate sensitivity for the binding of Siglecs to cancer cell lines. These results provide new insights into carbohydrate sulfation as a modification that is a general mechanism for tuning Siglec ligands on cells, including in cancer.

## Introduction

To appropriately discriminate *non-self* (e.g. pathogens) from *self* (our own healthy cells and tissues), immune cells employ many activatory and inhibitory immunomodulatory receptors, respectively.^1^ In this context, Sialic acid-binding immunoglobulin-type lectins (Siglecs) are cell surface proteins that act predominantly as inhibitory receptors to dampen immune responses to self-associated molecular patterns (SAMPs).^1–3^ Ligands of Siglecs are sialic acid-containing glycans and the interactions of Siglecs with their glycan ligands influence the location and organization of Siglecs on the cell surface, either bringing them closer to or keeping them away from other immunomodulatory receptors. Physiological conditions that bring a Siglec and activatory receptor together can lead to phosphorylation of the cytoplasmic immunoreceptor tyrosine-based inhibitory motif (ITIM) on Siglecs, which recruits SH2-containing SHP-1 or -2 phosphatases to dampen immune cell signaling.^4, 5^ Consistent with an important role for Siglecs in recognition of *self* under healthy conditions, Siglec-ligand interactions help prevent autoimmune disease, which may be the basis of an immune checkpoint.^6, 7^ Conversely, cancer cells exploit Siglecs by upregulating Siglec ligands on their surface to dampen or skew immune responses within the tumor microenvironment.^8, 9^

As the immunomodulatory roles of Siglecs are controlled via interactions with their sialic acid-containing carbohydrates,^10^ a better understanding of these glycan ligands is needed. For example, despite the demonstration by many groups of increased Siglec ligands in cancer,^8, 9^ a biochemical description of these changes in many cases is poorly understood. A better understanding of increased Siglec ligands in cancer may open up new avenues to modulate Siglec-ligand interactions as a potential therapeutic strategy.^5, 11^ Sialic acid-containing glycans can be appended to glycoproteins or glycolipids in mammals, and more recently evidence has emerged on RNA.^12, 13^ Interactions between Siglecs and their sialic acid-containing glycans occur through the N-terminal V-set domain of Siglecs, strongly relying on a key ionic interaction between the carboxyl group of the sialic acid and a conserved arginine.^14, 15^ The shallow glycan-binding groove for each Siglec, composed of antiparallel β-sheets, dictates the specificity for certain glycan structures. One relatively well-understood structural determinant in the glycan that dictates whether it is recognized by a Siglec is the linkage between the sialic acid and monosaccharide to which it is appended.^16^ For example, Siglec-2 (referred to below as CD22) selectively recognizes α2-6 linked sialosides, Siglec-3 (referred to below as CD33) can bind both α2-3 or α2-6 linked sialosides, Siglec-1 has a preference for α2-3 linked sialosides, and Siglec-11 is specific for α2-8 linked sialosides in the context of polysialic acid.^17, 18^ Additional factors such as the type of glycan to which the sialic is appended (e.g. *N*-glycan, *O*-glycan, glycolipid),^19, 20^ the underlying structural modifications (e.g. fucosylation and branching),^21, 22^ as well as post-glycosylation modification (e.g. acetylation and sulfation),^23, 24^ can significantly impact the affinity of glycans for Siglecs.^25, 26^ Moreover, the context in which these glycans are presented has also shown to play a role, such as the specific protein to which the glycans are attached and the density of presented glycans on the polypeptide backbone as in the case of *O*-glycans appended to mucins.^19, 27^

A growing body of evidence points to sulfation of the underlying glycan as being an important modification to the glycan structure that can increase the affinity of glycans for Siglecs.^26^ For example, human B-cells contain high levels of 6-*O*-GlcNAc sulfation, which maintains CD22 in a *masked* state on naive B-cells.^28, 29^ By glycan microarray, sialyl LewisX (sLe^X^) containing 6- *O*-sulfated Gal (6-S-sLe^X^) was the top hit as a ligand for Siglec-8-Fc.^30, 31^ Elegant affinity measurements and structural studies on Siglec-8 with sulfated sLe^X^ demonstrated that Siglec-8 displays a 40-fold enhanced affinity with 6-S-sLe^X^ and 4-fold with the isomeric species containing the sulfate appended to the GlcNAc (6’-S-sLe^X^).^32^ Interestingly, the disulfated compound (6,6’- S,S-sLe^X^) displayed only modest additivity of these affinity gains, with a further 1.6-fold enhancement in affinity relative to 6-S-sLe^X^. The structural basis for the affinity gain provided by 6-S-sLe^X^ was revealed to be an ionic interaction between an arginine within a variable loop adjacent to the glycan binding site; it is currently unknown if this mode of recognition of the sulfate is shared with other Siglecs but the primary sequence of this loop is variable in both size and composition between Siglec family members. Sialylated keratan sulfate was discovered as a ligand for Siglec-8 in human airways, which is significant as keratan sulfate often contains sulfation on the underlying Gal and GlcNAc.^33^ By glycan microarrays also revealed that Siglec-7 has enhanced binding to 6-S-sLe^X^ and 6’-S-sLe^X^ over sLe^X^, whereas Siglec-9 displayed a preference for 6’-S-sLe^X^.^24^ Consistent with these earlier glycan microarray results, ECV304 cells transfected with *carbohydrate sulfotransferase 2* (*CHST2*), which installs a sulfate onto the 6- position of GlcNAc, showed enhanced binding by Siglec-7 and Siglec-9. ^25^ Very recently, probing Siglec ligands on HEK293 cells transfected with a series of carbohydrate sulfotransferases (CHSTs) genes demonstrated that *CHST1*, which catalyzes the installation of the sulfate onto the 6-hydroxyl of Gal,^34^ enhanced the binding of Siglec-3, -7, -8, and -15.^35^

Although it is increasingly clear that sulfation plays a role in Siglec biology, numerous unanswered questions remain about the ability of CHST1 to enhance Siglec ligands, including context dependence, which was not studied in earlier work. Here, we overexpressed eight CHSTs in a variety of cell lines and discovered both enhanced and diminished binding of human and mouse Siglec-Fc proteins. Several of these enhanced interactions were cell type dependent. Through determining the affinity of CD33 for several synthetic glycan ligands, a disulfated ligand, Neu5Acα2-3(6-sulfo)Galβ1-4(6-sulfo)GlcNAc (6,6’-S,S-3SLN), was found to bind 36-fold tighter than the corresponding non-sulfated counterpart. Genetic analyses reveal that many cancer types upregulate *CHST1,* and several show poorer survival rates in patients with higher levels of *CHST1*. Moreover, decreasing carbohydrate sulfation lowers Siglec binding in cancer cell lines that express higher levels of *CHST1 or CHST2*. Taken together, our study reveals new insights into carbohydrate sulfation as a mechanism for upregulating cellular Siglec ligands, which is prevalent in cancer.

## Results

### Enhanced glycan sulfation by CHST overexpression

CHSTs are Golgi-localized enzymes that install sulfate at defined locations in glycans.^36^ Of the large family of human CHSTs, at least eight (CHST1, CHST2, CHST4, CHST8, CHST9, Gal3ST2, Gal3ST3, and Gal3ST4) are known to install sulfate onto glycan subclasses that contain sialic acid,^37^ which could potentially serve as Siglec ligands (Table S1). Therefore, the genes encoding these eight CHSTs were overexpressed initially in U937 cells through lentiviral transduction (Figure 1a). U937 cells, a monocytic cell line, were chosen because they are known to express Siglec ligands and interactions between Siglecs with their glycan ligands on white blood cells is physiologically relevant.^17^ Three different methods were used to characterize the glycosylation on the CHST-overexpressing cells: mass spectrometry (MS) of released *N*-glycans, lectin microarrays, and lectin staining in conjunction with flow cytometry (Figure 1b-f).

**Figure 1.**
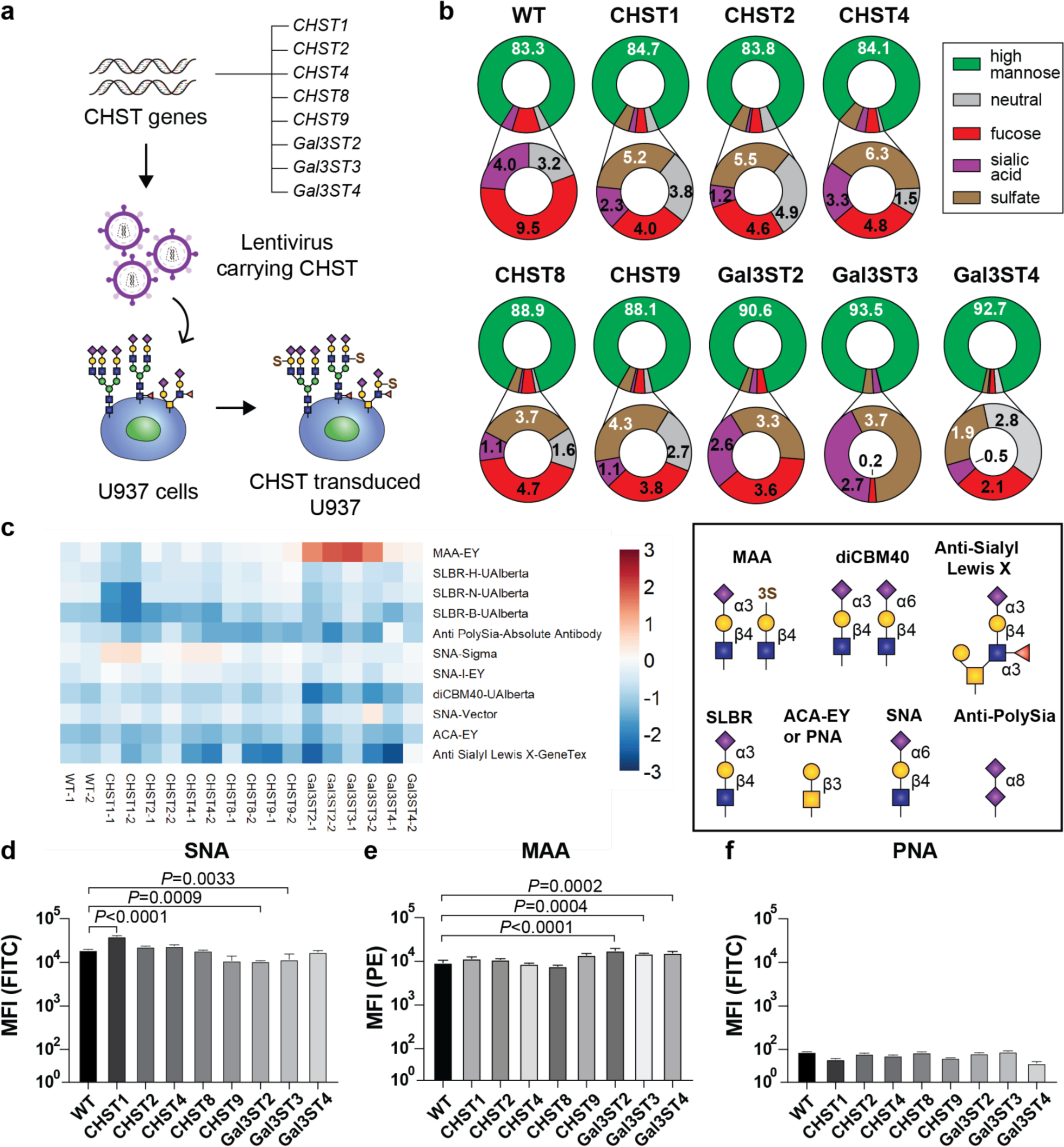
Characterization of CHST-overexpressing U937 cells. (a) Eight genes encoding CHSTs were transduced into U937 cells with lentivirus. (b) Mass spectrometry analysis of PNGase-F released *N*-glycans from the same number of CHST-transduced cells. The fraction (%) of each glycan class relative to the total is noted. (c) Select data from lectin microarray analysis on lysates from the *CHST*-transduced U937 cells. Lysates from each cell line were run in duplicate against a pooled reference. Median-normalized log2 ratio (Sample (S)/Reference (R) is shown. Full heatmap is shown in Figure S1. Glycan structures recognized by each lectin are shown to the right. (d-f) Flow cytometry of lectin binding, showing the mean fluorescence intensity of (d) SNA, (e) MAA, and (f) PNA. Data plots in d, e, and f are presented as mean ± SD. Statistical significance was calculated using a one-way ANOVA with Tukey’s multiple comparisons test.

By MS, enhanced sulfation was evident in all eight CHST-overexpressing lines (Figure 1b, Table S2 and S3). Indeed, 20-40% of unique, complex-type *N*-glycans contained a sulfate (accounting for an average of 4.2 ± 1.4% of the MS peak area) in the CHST-overexpressing lines, while no sulfated complex *N*-glycans were detected from the U937 cells transduced with control lentivirus. By lectin microarray, *Sambucus Nigra* lectin (SNA), which is specific for α2-6 linked sialosides independent of sulfation, showed enhanced binding to the *CHST1*-overexpressing cell lysate. Binding of *Maackia amurensis* lectin (MAA), which recognizes α2-3 linked sialosides or LacNAc-terminated glycans with 3-*O*-sulfated Gal, was highly enriched in the *Gal3ST2*- and *Gal3ST3*-overexpressing cell lysates (Figure 1c, Figure S1).^12, 38^ SLBR-H and -N, which are specific for α2-3 linked sialosides on *O*-linked glycans,^39^ and diCBM40, which recognizes both α2-3 and α2-6 linked sialosides,^40^ showed a decrease in signal towards the *Gal3ST2*- and *Gal3ST4*-overexpressing cell lysates. Likewise, a decrease in sLe^X^ structures were observed in the *CHST2-* and *Gal3ST4-*overexpressing cell lysates. The enhanced binding of MAA and decreased levels of α2-3 linked sialosides in *Gal3ST2-* and *Gal3ST3-*overexpressing cells strongly suggest a direct competition between sialyaltion and sulfation at the 3-position of terminal Gal. By flow cytometry, we find that SNA and MAA staining are consistent with the results from lectin microarray (Figure 1d,e). Binding of PNA, which recognizes desialylated Core1 *O*-glycans, was not altered (Figure 1f). While there are subtle changes in the levels of α2-3 and α2-6 linked sialic acid (Neu5Ac) structures in certain CHST-overexpressing cells, our results confirm that CHST overexpression increases carbohydrate sulfation on U937 cells.

### *CHST*-dependent binding of Siglecs to U937 cells

Recently, we produced a new panel of soluble human Siglec-Fc proteins for studying Siglec ligands on cells and tissues, with features that maximize recognition of their cellular ligands in a sialic acid-dependent manner (Figure 2a).^17^ Here, these Siglec-Fc probes were applied to study the effects of increased sulfation on human (Figure 2b-j) and mouse (Figure 2k) Siglec ligands. Significantly enhanced binding of CD33, Siglec-5, -7, -8, -14, and -15 were observed towards the *CHST1-*overexpressing U937 cells. Results for Siglec-8 were anticipated based on previous work, ^24, 32, 33^ while results with CD33, Siglec-7, and Siglec-15 are consistent with a recent study that overexpressed *CHST1* in HEK293 cells.^35^ However, the ability of *CHST1* overexpression to upregulate ligands for Siglec-5 and -14, which share an identical amino acid sequence in their first three domains^41^ (described from now on as Siglec-5/14), have not been described previously. A modest enhancement in binding of CD22 to the *CHST1*-overexpressing U937 cells was also not anticipated, but this may be due to enhanced expression of α2-6 linked sialosides, as evidenced by increased staining of SNA (Figure 1c,d). CD22 and Siglec-9 showed greatly enhanced binding to the *CHST2*- and *CHST4*-overexpressing U937 cells, which were anticipated from previous studies.^24, 26, 28^

**Figure 2.**
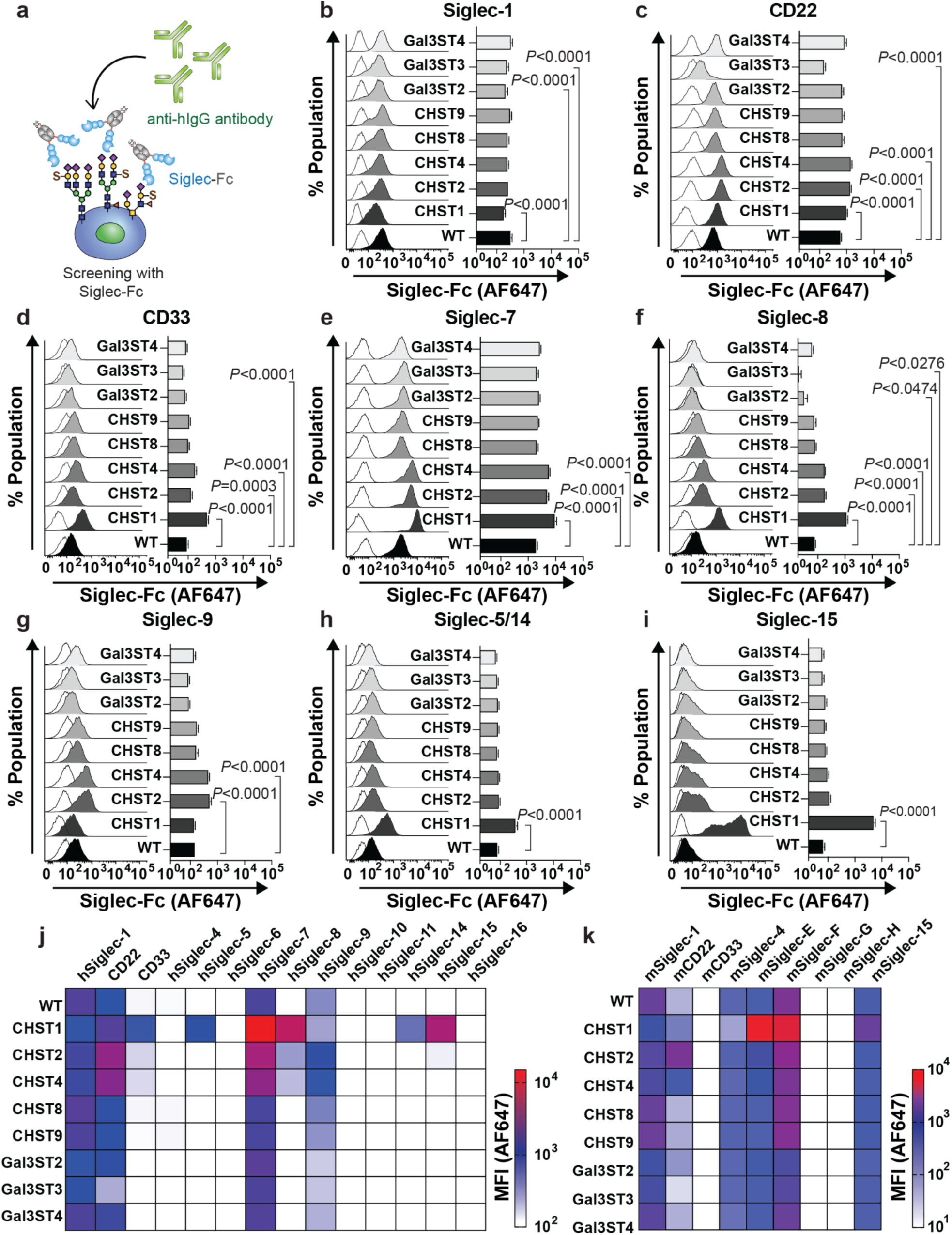
Probing the CHST-overexpressing U937 cells for Siglec ligands. (a) CHST-overexpressing cells were screened with Siglec-Fc proteins complexed with anti-human IgG-AF647. (b-i) Flow cytometry results (*left panels*) and summary of MFIs (*right panel*) for each Siglec-Fc screened against each CHST- overexpressing U937 cells. (j,k) Heatmap for both (j) human and (k) mouse Siglec-Fc binding to CHST- overexpressing U937 cell. Raw data for the mouse Siglec-Fc binding is show in Figure S2. Data plots in b-i are presented as mean ± SD. Statistical significance was calculated using a one-way ANOVA with Tukey’s multiple comparisons test.

Close inspection of the data indicates that binding of CD33, Siglec-5/14, Siglec-7, Siglec- 8, and Siglec-15 also showed modestly enhanced binding to the *CHST2*- and *CHST4*- overexpressing U937 cells. We also observed several interesting CHST-dependent decreases in binding. These include: Siglec-1 to *CHST1*-overexpressing U937 cells; Siglec-1, CD22, and CD33 to *Gal3ST3*-overexpressing U937 cells; and Siglec-8 to *Gal3ST2*- and *Gal3ST3*-overexpressing U937 cells. These latter results provide an intriguing possibility that sulfation can not only enhance Siglec binding, but also downregulate Siglec ligands.

Our previous study developing the new Siglec-Fc scaffold only focused on human Siglecs.^17^ Given the striking evolutionary differences between Siglecs in mouse and human,^42, 43^ and widespread use of mouse models to study concepts related to human health and disease,^44^ we developed a complementary set of mouse Siglec-Fc soluble proteins for this study and used them to examine binding to the CHST overexpressing U937 cells (Figure 2k, Figure S2). We find that mouse Siglecs conserved between mouse and human (Siglec-1, CD22, and Siglec-15)^45^ showed similar patterns as their human counterparts. Specifically, mouse CD22 (mCD22) showed greatly enhanced binding to *CHST2* and *CHST4*-overexpressing U937 cells, while mSiglec-15 showed enhanced binding to *CHST1*-overexpressing U937 cells. The significant binding enhancement for mCD22 was anticipated based on a 9-fold increase in affinity observed previously for Neu5Acɑ2-3Galβ1-4(6-*O*-sulfo)GlcNAc relative to its non-sulfated counterpart.^28^ For the more divergent CD33-related Siglecs, less predictable patterns were observed. Binding of mouse CD33, which has divergent properties compared to its human counterpart,^46^ was not enhanced in the *CHST1*-overexpressing U937 cells. These results have important implications for studying the role of CD33 on brain microglia and its relationship to Alzheimer’s disease susceptibility and the use of mouse models to study the relationship of CD33 to neurodegeneration.^46, 47^ Siglec-E, which is described as the mouse ortholog of Siglec-9,^26^ showed a similar binding pattern to Siglec-7, where binding to cells was predominantly enhanced by either *CHST1* or *CHST2/4* overexpression, which is consistent with findings from a previous study on Siglec-E.^48^ Lastly, Siglec-F binding was increased to *CHST1*-overexpressing U937 cells, which was anticipated based on previous work.^26^

### *Sialic acid-dependent binding and generality* of CHST-dependent Siglec binding

To examine whether the enhanced binding of Siglecs to CHST-overexpressing cells was sialic acid dependent, a complementary set of the CHST-overexpressing U937 cells were generated on a *CMAS*^-/-^ background, which we generated previously.^17^ In all cases, no binding was observed compared to WT U937 cells (Figure 3a). Moreover, *Arthrobacter Ureafaciens* neuraminidase was used to hydrolyze sialic acid residues, which likewise produced a near complete disappearance of ligands for CD33, Siglec-8, and Siglec-15 in *CHST1*-overexpressing U937 cells as well as CD22 and Siglec-9 in *CHST2*-overexpressing U937 cells (Figure 3b-c). These results establish that enhanced binding to sulfated cellular glycans still requires sialic acid.

To investigate to what extent our findings of enhanced Siglec binding to CHST- overexpressing U937 cells extend to other cells, *CHST1* and *CHST2* were transduced into four additional cell lines: K562, Jurkat, HEK293, and A549 cells. Similar trends were observed for many of the key CHST-upregulated Siglec binding interactions in U937 cells (Figure 3d-g). Interestingly, enhanced binding of Siglec-5/14 to the *CHST1*-overexpressing HEK293 cells was not observed previously,^35^ but was readily observed in our studies, which we speculate may be attributed to non-functional commercial Siglec-5/14-Fc. In contrast with the other cells, CD33, Siglec-5/14 and Siglec-15 showed either minimal or no increase of binding to the *CHST1-*overexpressing Jurkat cells. Moreover, the modest binding increase of CD33 in the *CHST2*-overexpressing U937 was not observed in the other cells. These results suggest that the correct ensemble and/or the unique repertoire of proteins expressed in each cell type impact the ability of CHST1 and CHST2 to create Siglec ligands, but overall point to carbohydrate sulfation as being an important mechanism for upregulating Siglec ligands.

**Figure 3.**
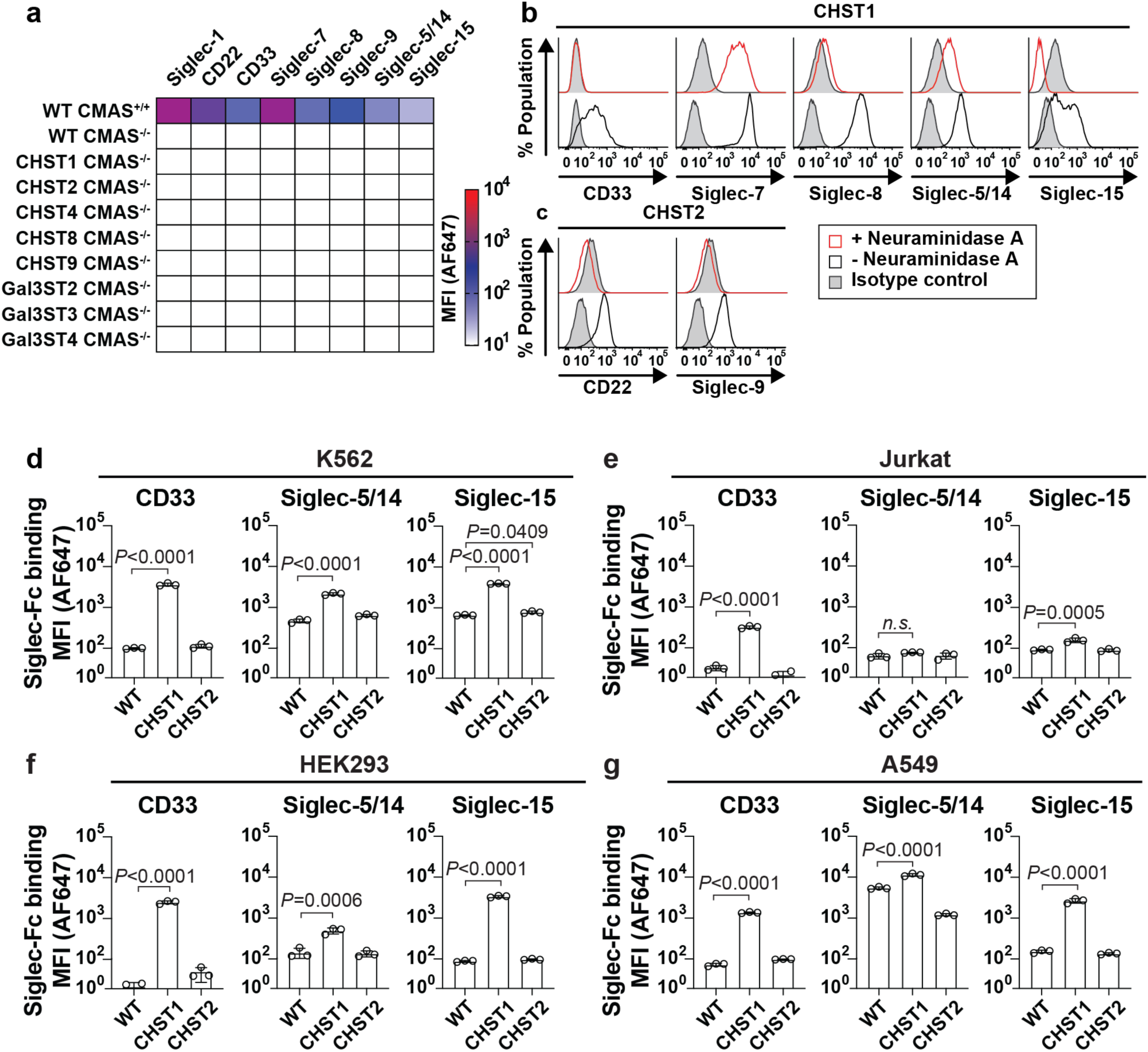
Sialic acid-dependent Siglec binding to CHST overexpressing cells. (a) Siglec-Fc binding to the CHST overexpressing *CMAS*^-/-^ U937 cells. (b,c) Probing the CHST1 and CHST2 overexpressing U937 cells with Siglec-Fc after neuraminidase digestion. (d-e) the four additional cell lines (K562, Jurkat, HEK293, and A549) transduced with either CHST1 or CHST2 were probed with CD33, Siglec-5/14, and Siglec-15. Data plots in d,e,f, and g are presented as mean ± SD. Statistical significance was calculated based on a one-way ANOVA with Tukey’s multiple comparisons test.

### Pharmacological perturbation of key cellular glycans

Kifunensine, benzyl-α-GalNAc, and Genz-123346 are inhibitors for the three major classes of glycans that contain sialic acid (Figure 4a). These three inhibitors were used in the CHST-transduced U937 cells to examine what sub-class of glycans are elaborated with sulfation and enhance binding of Siglecs (Figure 4b-h). As anticipated based on previous observations for CD22 preferring sialoside presented on *N*-glycans, ^49^ CD22-Fc binding was completely abolished in the kifunensine-treated *CHST1*- and *CHST2*-overexpressing U937 cells (Figure 4b). CD33, Siglec-5/14, Siglec-9, Siglec-15 were affected by inhibition of either *N*- or *O*-glycosylation (Figure 4c,f, g,h), suggesting that their sulfated ligands are presented on both classes of glycans. Benzyl- α-GalNAc significantly reduced Siglec-7 binding to the *CHST1*- and *CHST2*-overexpressing U937 cells (Figure 4d), which is in line with a recent study that found Siglec-7 ligands are sensitive to treatment with the mucin-selective protease, StcE.^50^ Interestingly, Siglec-8 binding was not perturbed by either kifunensine or benzyl-α-GalNAc treatment (Figure 4e). Pharmacological blockade of glycolipid biosynthesis did not perturb Siglec ligands in any instance.

**Figure 4.**
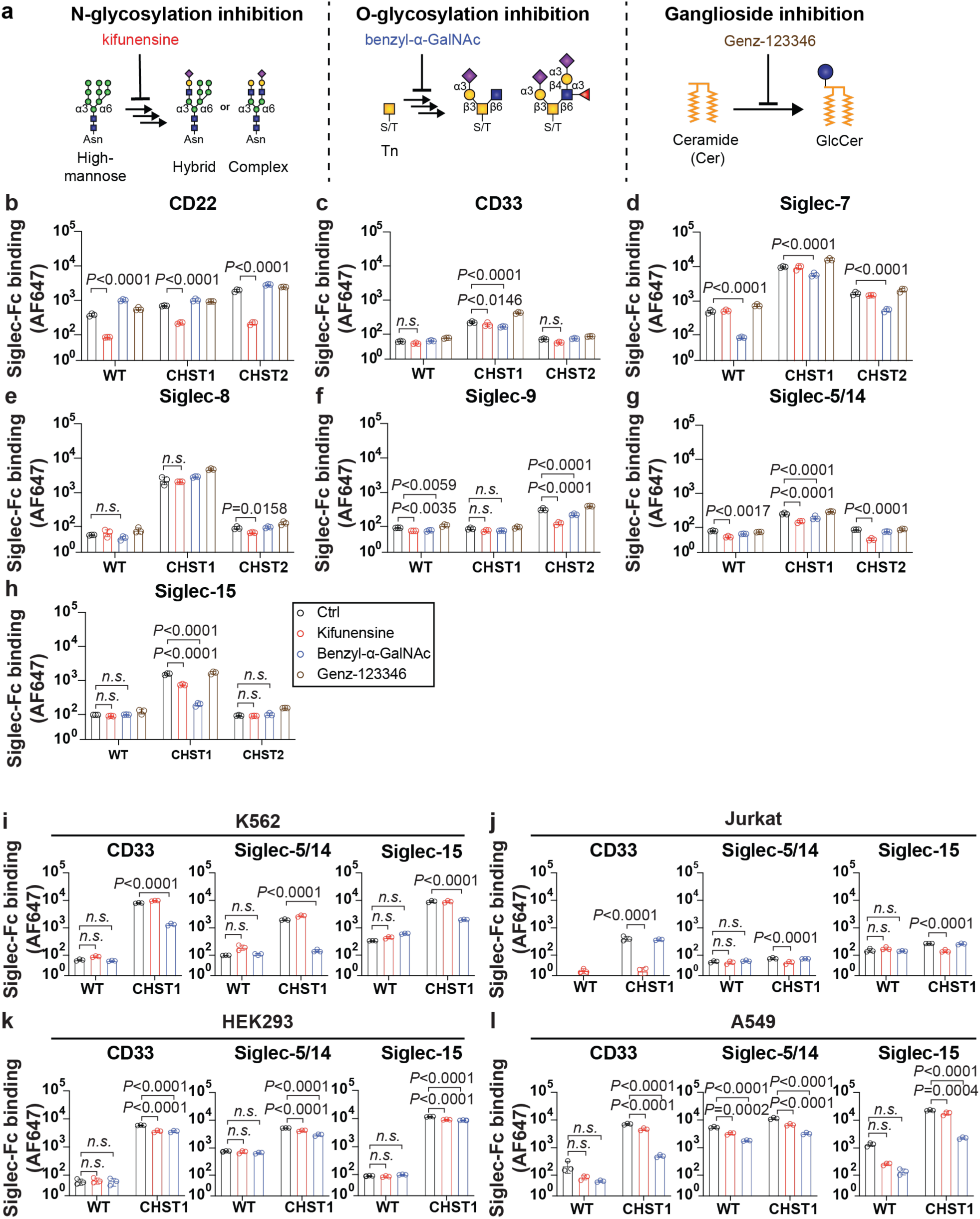
Inhibitors of cellular glycosylation reveal a cell type-specific pattern of Siglec ligands. (a) Inhibitors for *N*-glycosylation, mucin-type *O*-glycosylation, and ganglioside biosynthesis. (b-h) binding of CD22, CD33, Siglec-7, 8, 9, 14, and 15 to the WT, *CHST1*-, and *CHST2*-overexpressing U937 cells with inhibition of each glycosylation pathway. (i-l) K562, Jurkat, HEK293, and A549 overexpressing *CHST1* or EV were probed for ligands of CD33, Siglec-5/14 and Siglec-15 in cells treated with kifunensine or benzyl- α-GalNAc. Data are presented as mean ± SD. Statistical significance was calculated using a two-way ANOVA with Tukey’s multiple comparisons test.

To validate whether the glycosylation inhibition patterns are conserved in other cells, we further tested K562, Jurkat, HEK293, and A549 cells transduced with empty vector (EV), *CHST1*, or *CHST2* with CD33, Siglec-5/14, and 15 (Figure 4i-l). Cell-specific patterns emerged; benzyl-α- GalNAc was dominant in decreasing the binding of all three Siglecs in all types of K562 and A549 cells (Figure 4i,l), while kifunensine effectively decreased binding of all Siglecs in the *CHST1*- overexpressing Jurkat cells (Figure 4j). This later observation may be partially explained by the fact that mucin-type *O*-glycans in Jurkat cells are highly truncated, being composed primarily of the Tn antigen (Gal*α*1-3GalNAc).^51^ Kifunensine and benzyl-α-GalNAc had only minor effects in blocking Siglec binding in HEK293 cells (Figure 4k). Previous studies overexpressing CHSTs in HEK293 using genetic knockouts of subclasses of glycans made conclusions about where sulfated glycans are presented,^35^ which differ to some degree with the results presented here using the pharmacological approaches. Overall, our results strongly suggest that binding enhancement of Siglecs in *CHST1* or *CHST2*-overexpressing cells is cell type-specific and may be presented on different sub-classes of glycans on different cell types.

### Quantifying Siglec binding to sulfated glycans

Previously, we developed an electrospray ionization (ESI)-MS assay for quantifying the affinity of Siglecs for their glycan ligands.^12, 17^ Advantages of this approach are that it is label free, fast, has low sample consumption, can quantify very weak interactions, and the results are consistent with biochemical approaches such as isothermal titration calorimetry.^52^ The enhanced binding of human CD33 to several of the CHST-overexpressing cell lines and the potential relevance of these findings to glycan ligands of CD33 in Alzheimer’s disease susceptibility motivated us to apply this MS-based approach to quantify the affinity enhancement afforded by sulfation.^46, 47^ In our previous studies, we unexpectedly found that approximately half of the soluble CD33 was *O*-glycosylated.^17^ Such heterogeneity makes analyzing ligand binding more challenging because of spectral overlap of peaks corresponding to the protein without the *O*- glycan and those of the *O*-glycosylated glycoforms bound to ligands of similar molecular weights. Attempting to remove the *O*-glycans enzymatically, with *O*-glycosidase, was not successful even after first removing the sialic acid residues with a neuraminidase. Sites of *O*-glycosylation were mapped by MS and evidence of *O*-glycosylation was observed at T239 and T260 (Figure 5a,b). The latter site is located within the linker between CD33 and the Fc and is encoded by the AgeI restriction site used for the molecular cloning. Accordingly, a double mutant (T239A/T260A) of CD33-Fc was prepared in LEC1 CHO cells. Following removal of the Fc and high mannose *N*- glycans by Tobacco Etch Virus (TEV) protease and Endo-H endo-glycosidase, respectively, SDS- PAGE showed a single band for T239A/T260A CD33 in contrast to WT CD33, which runs as doublet by SDS-PAGE (Figure 5c,d). The mass spectrum of the T239A/T260A CD33 confirmed that it lacks *O*-glycosylation.

**Figure 5.**
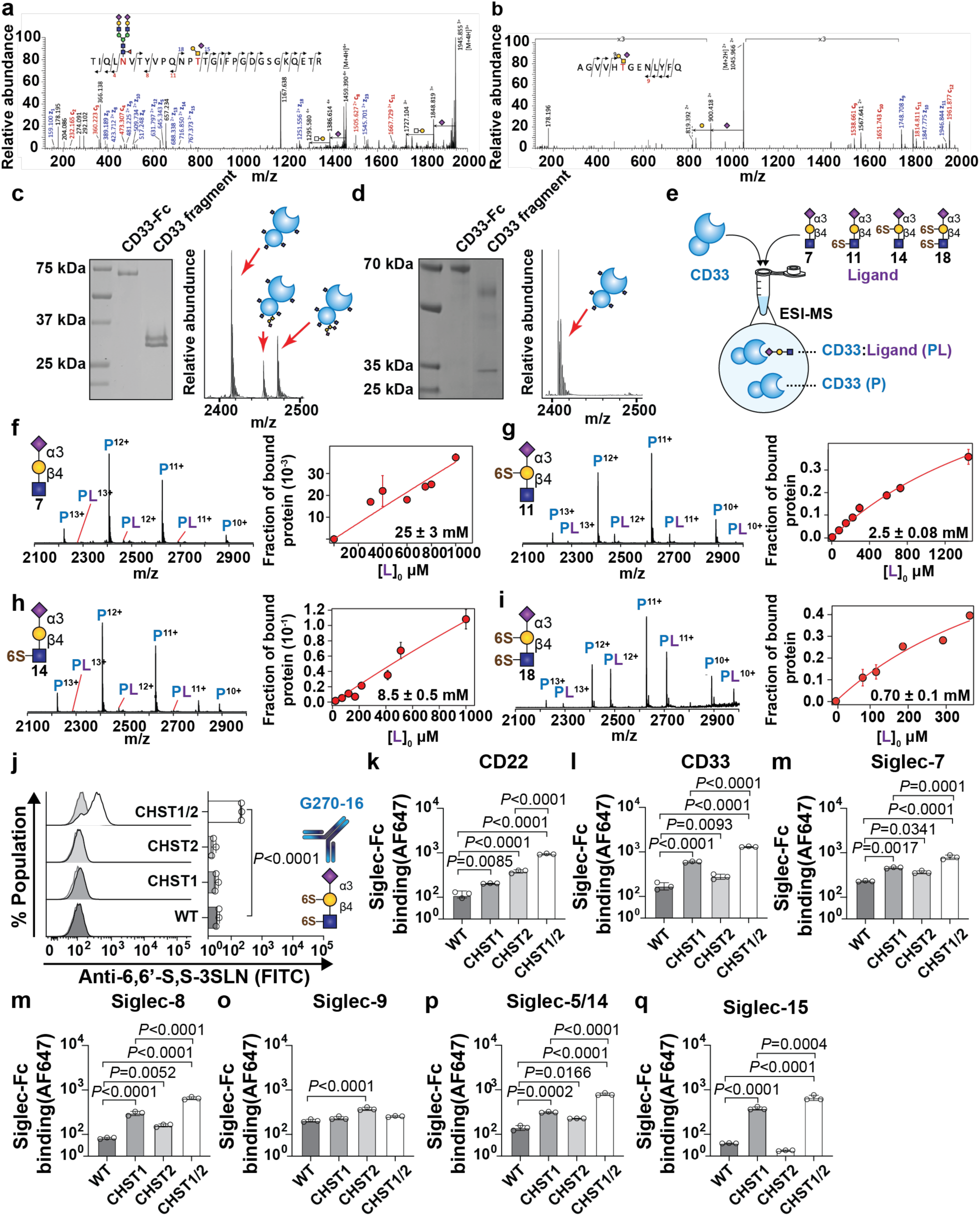
CD33 shows a strong preference for disulfated ligands. (a,b) EThcD MS^2^ analyses of glycopeptides derived from CD33 unambiguously identified *O*-glycosylation at (a) T239 and (b) T260. In addition to HexNAc_1_Hex_1_NeuAc_1_ shown here, two other O-glycans were detected for T239, corresponding to HexNAc_1_Hex_1_, and HexNAc_1_Hex1NeuAc_2_. (c-d) SDS-PAGE of the (c) WT CD33 fragment and (d) T239A/T260A CD33 fragment. (e) Scheme for direct binding measurement by ESI-MS to determine the affinity of the CD33 against the compound **7**, **11**, **14**, and **18**. (f-i) Titration curve of (f) 3SLN **7** (400 μM), (g) 6-S-3SLN **11** (280 μM), (h) 6’-S-3SLN **14** (400 μM), and (i) 6,6’-S,S-3SLN **18** (100 μM) to determine *K*_d_ of CD33 on each compound. (j) G270-16 staining on the U937 cells transduced with both *CHST1* and *CHST2*. (k-q) Probing *CHST1/2*-overexpressing cells with (k) CD22, (l) CD33, (m) Siglec-7, (n) Siglec-8. (m) Siglec- 8, (o) Siglec-9, (p) Siglec-5/14, and (q) Siglec-15. Data are presented as mean ± SD. Statistical significance was calculated using a one-way ANOVA with Tukey’s multiple comparisons test.

Four differentially sulfated compounds were chemo-enzymatically prepared based on a Neu5Acα2-3LacNAc-propylamine. Specifically, we prepared a non-sulfated compound **7** (3SLN), 6-*O*-sulfate at the Gal **11** (6-S-3SLN), 6-*O*-sulfate at the GlcNAc **14** (6’-S-3SLN), and disulfated at both positions **18** (6,6’-S,S-3SLN) (Figure S3). The T239A/T260A CD33 protein was titrated with these ligands to determine the dissociation constant (*K*_d_) by ESI-MS (Figure 5e). The affinity measured against **7** was surprisingly weak, with a *K*_d_ value of 25 ± 3 mM (Figure 5f). Such a weak affinity was not anticipated given that our previous measurements of CD33 against 3SLN (lacking the propylamine aglycone) yielded a *K*_d_ value of 2.7 ± 0.1 mM.^17^ To test whether a lack of the *O*- glycosylation on CD33 decreased affinity, we prepared the WT CD33 fragment and found a similarly weak affinity for 3SLN-propylamine (Figure S4a). Moreover, the T239A/T260A CD33 fragment yielded a similar *K*_d_ for 3SLN without the propylamine linker, in line with the WT CD33 fragment (Figure S4b). Thus, the propylamine aglycone has an unexpected deleterious effect on the binding affinity of CD33. The crystal structure of CD33 presents no obvious rationale for this loss of affinity.^53^ This surprising finding has important implications for the future design of high affinity glycans ligands for targeting CD33. Aware of this linker effect, we moved ahead and tested T239A/T260A CD33 against **11**, **14**, and **18** (Figure 5g-i). We find that the 6-*O*-sulfate modification at the Gal enhances the affinity by 10-fold (*K*_d_ = 2.5 ± 0.1 mM), (Figure 5g) while the 6-*O*-sulfate modification at the GlcNAc enhanced the affinity by 3-fold (*K*_d_ = 8.5 ± 0.5 mM) (Figure 5h). The disulfated compound **18** dispalyed the highest affinity (*K*_d_ = 0.70 ± 0.1 mM) for CD33, which is a 36-fold affinity enhancement relative to the non-sulfated **7.** (Figure 5f,i).

The only other study to perform quantitative binding measurements on a similar series of sulfated compounds was an investigation of Siglec-8 with sLe^X^ structures.^32^ This previous study found that a 6-*O*-sulfation at the Gal dominated the affinity gains, with a disulfated ligand showing only modestly increased affinity 1.6-fold relative to the best monosulfated ligand. Our binding measurements with CD33 demonstrate a 3.6-fold increase in affinity for the disulfated ligand relative to the best monosulfated ligand. Glycan microarray results, which represent semi-quantitative binding, suggest that additivity between the two sulfate modifications may also be at play for Siglec-7.^24^ Therefore, to our knowledge, our results with CD33 represents the largest enhancement in affinity provided by disulfation.

### Overexpression of both CHST1 and CHST2 in U937 cells greatly enhances Siglec ligands

The quantitative binding measurements above for the two monosulfated trisaccharides **11** and **14** are in line with results for CD33-Fc binding to the *CHST1* and *CHST2 or 4* overexpressing cells, respectively (Figure 2d). The additive effect in the disulfated species **18** suggests that dual expression of CHST1 and CHST2 will significantly enhance cellular ligands of CD33. To test this, *CHST1*-transduced U937 cells were further transduced with lentivirus carrying *CHST2* or an EV control. To accomplish this, our lentiviral vector was re-engineered to express an orthogonal fluorescent protein (EGFP) to mAmterine, enabling sorting of doubly transduced cells. The U937 cells overexpressing both *CHST1* and *CHST2* (described herein as *CHST1/2*) showed enhanced staining with an antibody that recognized a 6,6’-S,S-3SLN (clone G270-16), providing evidence that this strategy had been successful (Figure 5j). Comparing the binding of CD33 to the *CHST1/2*-overexpressing U937 cells to the WT, *CHST1*-, and *CHST2*-overexpressing cells, we find enhanced binding to the former (Figure 5i). Similar results were observed for CD22, CD33, Siglec-7, Siglec-8, Siglec-5/14, and Siglec-15 but, curiously, Siglec-9 did not show this effect (Figure 5k-q). In harmony with this, parts of these results are consistent with the previous study showing the additive effect of disulfation over 6-*O*-sulfation at Gal was valid in the binding of Siglec-7 and -8, but not in Siglec-9.

Staining of cells with CD33-Fc represents *trans* ligands. To examine *cis* ligands of CD33, we performed binding studies with fluorescent liposomes displaying a high affinity and selective ligand of CD33, described previously.^54^ The ability of CD33 on U937 cells to engage the liposome is a measure of its masked state,^17^ and therefore a measure of the engagement of CD33 by *cis* ligands. CD33 levels on U937 cells were first verified to be unchanged in the WT, *CHST1*-, *CHST2*-, and *CHST1/2*-overexpressing U937 cells (Figure S5a). Examining liposome binding by flow cytometry, we find that *CHST1*-overexpressing U937 cells showed significantly reduced engagement by CD33 ligand liposomes. *CHST2*- overexpressing U937 showed a small decrease in liposome binding relative to the WT U937 cells, but this was not statistically significant (Figure S5b). On the other hand, *CHST1/2*-overexpressing U937 showed the least amount of binding to the liposomes, demonstrating that CD33 is strongly masked on these cells. These results for *cis* ligands on U937 cells overexpressing both *CHST1* and *CHST2* provide further support that a disulfated ligand is a strong ligand for CD33.

### Upregulated expression of CHST1 in cancer

Elevated levels of sialic acid on cancer cells have emerged as a key player in controlling anti-tumour immune response, by engaging with Siglecs on immune cells.^5, 8^ Given the enhanced binding of Siglecs to their sulfated glycan ligands generated by *CHST1* overexpression, we hypothesized that cancer cells utilize glycan sulfation to further increase their affinity towards Siglecs and potentially enhance immune suppression. To evaluate expression levels of *CHST1* in cancer, we retrieved transcript data of 14 tumour types from The Cancer Genome Atlas (TCGA) with more than five matched non-malignant and primary tumour samples. Strikingly, 9 of 14 cancer types display significant upregulation in *CHST1* levels compared to their non-malignant tissue counterparts (Figure 6a). We next assessed whether expression of *CHST1* can predict patient outcome in 19 different cancer types. We used publicly available data from TCGA and kmplot.com and split cases into low and high expression levels based on median *CHST1* expression levels. Kaplan-Meier curves of 5-year overall survival revealed that *CHST1* expression is a predictor of poor patient outcome in 7 of 19 cancer types. These include lung adenocarcinoma (LUAD), stomach adenocarcinoma (STAD), breast invasive carcinoma (BRCA), liver hepatocellular carcinoma (LIHC), uveal melanoma (UVM), glioblastoma multiforme (GBM), and mesothelioma (MESO, Figure 6b). In two cancer types, ovarian serous cystadenocarcinoma (OV) and pancreatic adenocarcinoma (PAAD), *CHST1* expression correlated with favourable patient outcomes. Collectively, these results demonstrate that *CHST1* is upregulated in the majority of cancer types and can correlate with poorer patient outcomes in specific types of cancer. Given the numerous mechanisms of immune evasion used by tumors,^2, 55^ which can vary based on cancer type, it is unsurprising that these effects are not uniform across all cancers.

**Figure 6.**
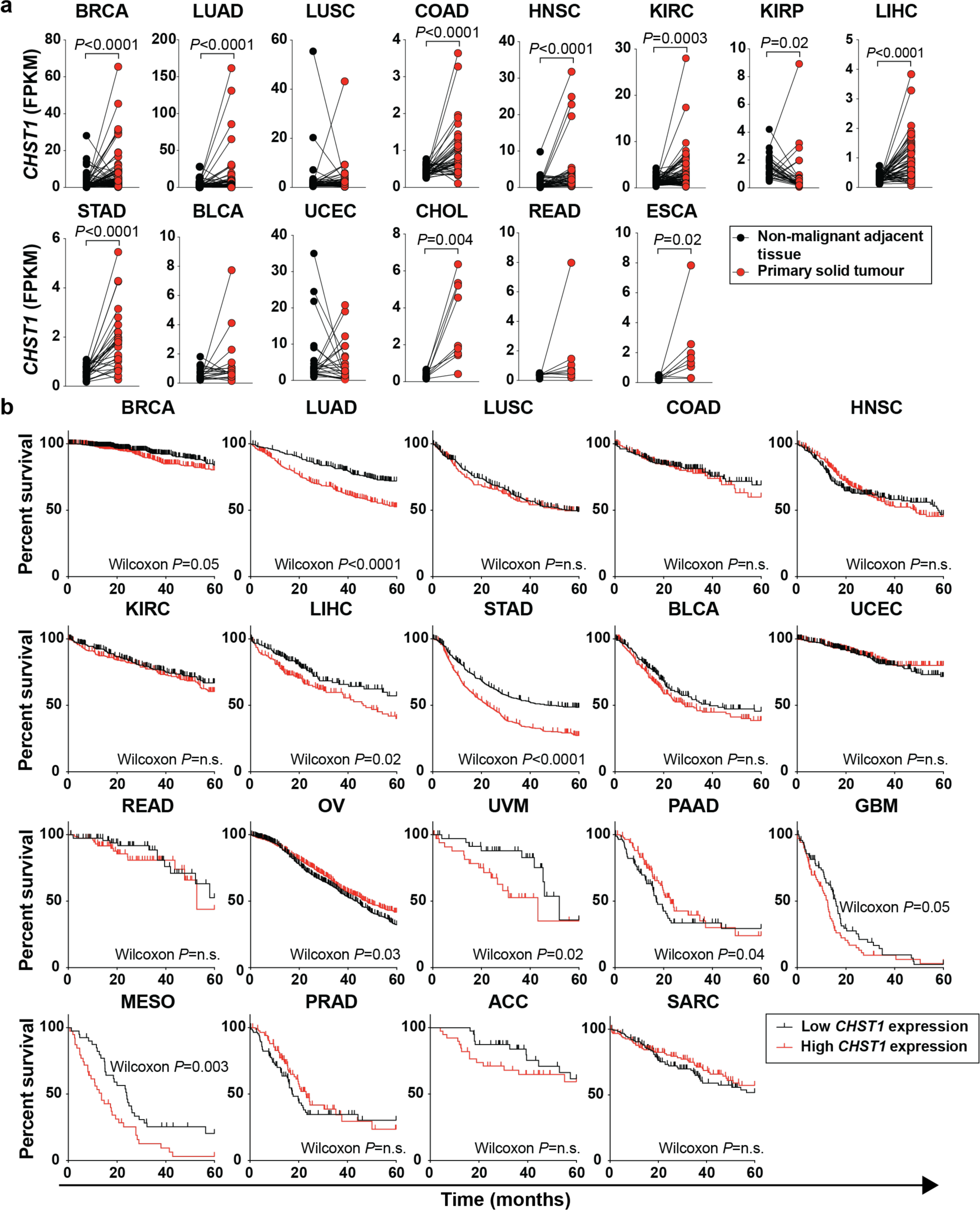
Upregulated *CHST1* expression in solid tumours correlates with poor patient outcome in several cancer types. (a) Transcript profiling of *CHST1* in patient-matched non-malignant adjacent and cancerous tissues in 14 cancer types from The Cancer Genome Atlas (TCGA) Consortium. Normalized *CHST1* expression (FPKM) data from cancer types with >5 matched samples were downloaded through the University of California Santa Cruz (UCSC) Xena tool (https://xenabrowser.net). TCGA cancer types included in the analysis are as follow: BRCA (breast cancer), LUAD (lung adenocarcinoma), LUSC (lung squamous cell carcinoma), COAD (colon adenocarcinoma), HNSC (head and neck squamous cell carcinoma), KIRC (kidney renal clear cell carcinoma, KIRP (kidney renal papillary cell carcinoma), LIHC (liver hepatocellular carcinoma), STAD (stomach adenocarcinoma), BLCA (bladder urothelial carcinoma), UCEC (uterine corpus endometrial carcinoma), CHOL (cholangiocarcinoma), READ (rectum adenocarcinoma), and ESCA (esophageal carcinoma). Statistical analysis was performed using Wilcoxon matched-pairs signed-rank test. (b) Kaplan-Meier 5-year overall survival curves of 19 cancer types based on *CHST1* expression. Patients were stratified based on median *CHST1* expression. Curated datasets for LUAD (lung adenocarcinoma), LUSC (lung squamous cell carcinoma), STAD (stomach carcinoma), and OV (ovarian serous cystadenocarcinoma) were retrieved from https://kmplot.com. TCGA datasets for BRCA(breast cancer), LIHC (liver hepatocellular carcinoma), KIRC (kidney renal clear cell carcinoma), BLCA (bladder urothelial carcinoma), HNSC (head and neck squamous cell carcinoma) PRAD (prostate adenocarcinoma), READ (rectum adenocarcinoma), COAD (colon adenocarcinoma), UVM (uveal melanoma), PAAD (pancreatic adenocarcinoma), GBM (glioblastoma multiforme), MESO (mesothelioma), UCEC (uterine corpus endometrial carcinoma), ACC (adrenocortical carcinoma), and SARC (sarcoma) were downloaded through the UCSC Xena tool (https://xenabrowser.net). Statistical analysis was performed using a Gehan-Breslow-Wilcoxon test.

### Disrupting glycan sulfation levels on cancer cells downregulates Siglec ligands

Given the results presented above demonstrating that *CHST1* is upregulated in numerous cancer types and correlates with poorer survival rates in a number of cancers, we wondered if upregulated *CHST1* expression gives rise to enhanced Siglec ligands on cancer cells. Sodium chlorate (NaClO_3_) is an *in vitro* inhibitor of 3’-phosphoadenosine 5’-phosphosulfate (PAPs) biosynthesis, and has been previously used to block the installation of sulfation on carbohydrates by CHSTs (Figure 7a).^56, 57^ To establish whether the increased binding of Siglecs to the CHST- transduced cells can be abolished by NaClO_3_, we treated the *CHST1*-overexpressing U937 cells with NaClO_3_ from 0 mM to 50 mM for two days to assess the binding of CD33-Fc and found a clear dose-dependent decrease in CD33-Fc binding (Figure 7b). Although 50 mM of NaClO_3_ showed the largest decrease, we chose to use 25 mM for subsequent experiments as there was minimal effect on cell viability at this concentration (Figure S6a). We additionally observed that the enhanced binding of CD33, Siglec-7, -8, -14, and -15 on the *CHST1*-overexpressing cells and CD22 and Siglec-9 on the *CHST2*-overexpressing U937 cells was abrogated by 25 mM NaClO_3_ treatment, providing further evidence for the critically importance role of sulfation in Siglec binding (Figure 7c-i). NaClO_3_ treatment did not perturb the binding of these Siglecs to WT U937 cells, suggesting that these cells do not express sulfated glycans, which is consistent with our MS data (Figure 1b).

**Figure 7.**
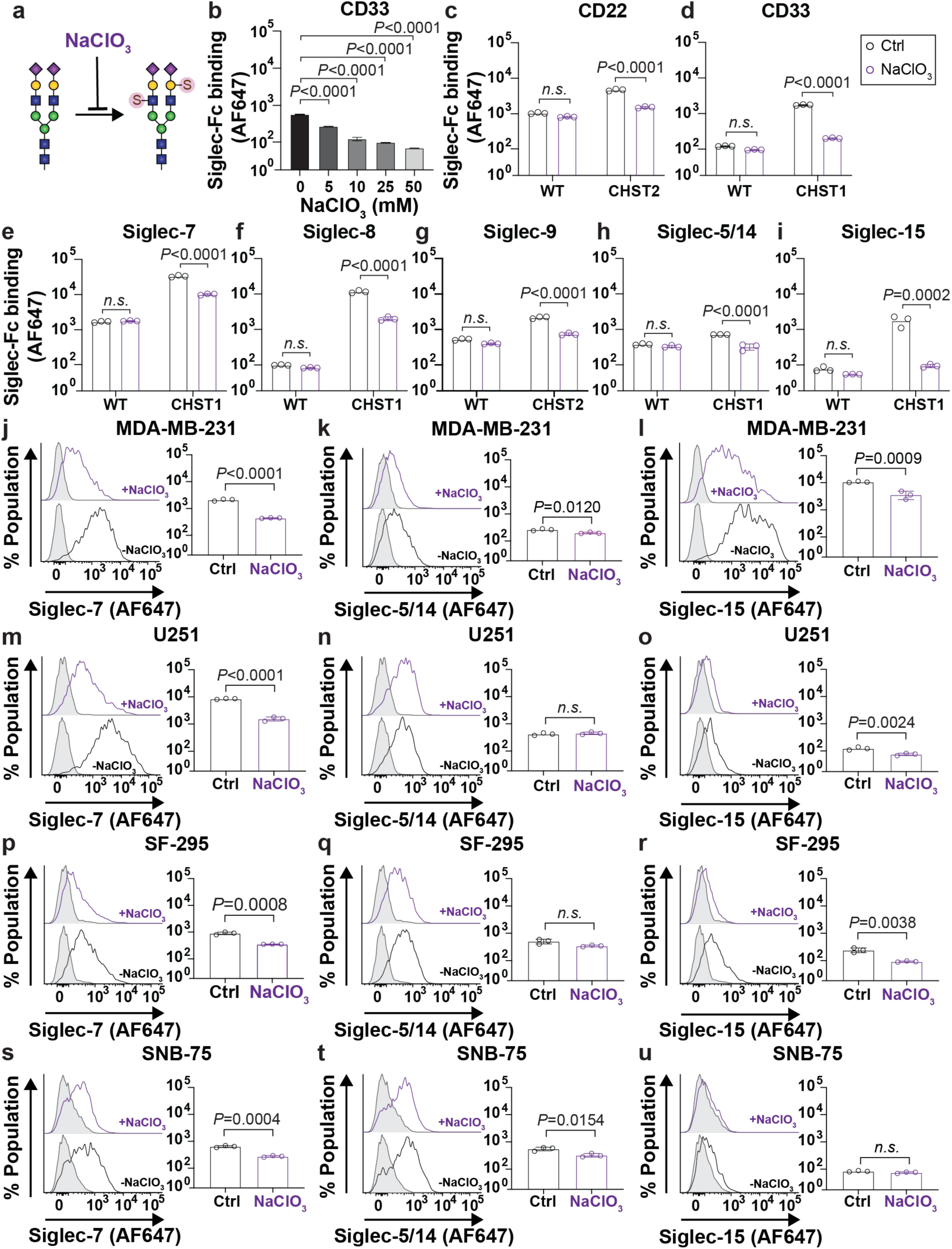
Inhibiting carbohydrate sulfation on cancer cells decreases Siglec ligands. (a) Illustration of NaClO_3_ for inhibiting cellular carbohydrate sulfation (b) Titration of NaClO_3_ in CD33 binding to the *CHST1*- overexpressing U937 cells. (c-i) The effect of NaClO_3_ on the binding (c) CD22, (d) CD33, (e) Siglec-7, (f) Siglec-8, (g) Siglec-9, (h) Siglec-5/14, and (i) Siglec-15 to the EV, *CHST1*-, and *CHST2*-overexpressing U937 cells. (j-u) The effects of inhibiting sulfation with NaClO_3_ on Siglec-Fc binding were assessed in (j-l) MDA-MB-231, (m-o) U251, (p-r) SF-295, and (s-u) SNB-75 with Siglec-7 (j,m,p,s), Siglec-5/14 (k,n,q,t) and Siglec-15 (l,o.r,u). Data plots in b-r are presented as mean ± SD. Statistical significance was calculated using a two-way ANOVA with Sidak’s multiple comparisons test (c-i) and unpaired t tests (j-u).

Next, we tested NaClO_3_ on a variety of cancer cell lines to determine if Siglec binding was abrogated. Based on transcript levels of each *CHST* gene in the NCI-60 cancer cell library,^58^ we selected MDA-MB-231, U251, SF-295, and SNB-75 as promising candidates since those cells express higher levels of *CHST1* or *CHST2* (Table S4). Conversely, CCRF-CEM, MCF-7, A549, and MDA-MB-468 show low levels of *CHST1* and *CHST2* expression. Therefore, we analyzed whether NaClO_3_ sensitivity to Siglec-Fc binding correlates with expression of these *CHST*s with these cancer cell lines, focusing on Siglec-7, -5/14, and -15 given that these Siglecs show robust binding to these cell lines. Viability of each cell was minimally affected by 25 mM NaClO_3_ treatment (Figure S6b). In the four high C*HST1-* or *CHST2*-expressing cancer cell lines, inhibiting endogenous sulfation led to decreases in Siglec binding in all cases except for Siglec-15 to MDA-MB-231 and Siglec-5/14 to U251 and SF-295 cells (Figure 7j-u). These results reinforce our earlier data showing that the context in which carbohydrate sulfation is presented is important, with the correct ensemble of underlying glycans required to support sulfation as a mechanism for increasing Siglec ligands. In the low expressing *CHST1* and *CHST2* cells (A549, MDA-MB-468, CCRF-CEM, and MCF-7), Siglec-7, Siglec-5/14 and Siglec-15 did not show any binding decrease after NaClO_3_ treatment (Figure S7). Treatment with NaClO_3_ did not result in a significant decrease to either SNA or MAA binding to CCRF-CEM, K562, MDA-MB-231, U251, SF-295, and SNB-75 cells (Figure S8). Taken together, expression levels of *CHST1* or *CHST2* in cancer cell lines correlate with sensitivity of Siglec ligands to inhibition of endogenous carbohydrate sulfation, strongly suggesting that enhanced carbohydrate sulfation is a mechanism by which cancer cells upregulate Siglec ligands.

## Conclusions

Our results demonstrate that overexpression of either *CHST1* or *CHST2* on several cancer cell lines by lentivirus transduction increases cellular sulfated sialosides, which enhances binding to CD22, CD33, Siglec-5/14, Siglec-7, Siglec-8, Siglec-9, and Siglec-15. To our knowledge, this is the first demonstration of *CHST1*-dependent up-regulation of Siglec-5/14 ligands. Although enhanced sulfation increases the levels of Siglec ligands in all cells tested, cell type-specific patterns emerged, pointing to the importance of the underlying glycome for supporting sulfated glycans as Siglec ligands. By overexpressing both *CHST1* and *CHST2* in cells to create disulfated glycans, we observed dramatically enhanced binding increases of several Siglecs (CD22, CD33, Siglec-7, Siglec-8, Siglec-5/14, and Siglec-15). This preference for disulfated glycans by human CD33 is supported by ESI-MSI affinity measurements, which showed that 6,6’-S,S-3SLN demonstrated a 36-fold increase in affinity relative to its non-sulfated counterpart. Interestingly, the binding enhancement by overexpression of *CHST1* or *CHST2* or both in CD33-Fc was not conserved with mCD33-Fc. Furthermore, we reveal that high expression of *CHST1* is found in many cancers and can lead to poor survival outcomes. The implications of upregulated *CHST* expression in cancer on Siglec ligands was demonstrated by showing that NaClO_3_ abrogates Siglec binding in cancer cells that express high levels of *CHST1* or *CHST2*. Therefore, we propose carbohydrate sulfation as a mechanism that cancer cells can exploit for enhancing the interactions with Siglecs.

## Methods

### Cell culture

All cell lines were obtained from ATCC, maintained under sterile conditions, cultured with either DMEM or RPMI (Gibco) containing 10% FBS (Gibco), 50 U/mL Penicillin/Streptomycin (Gibco), and 100 mM HEPES (Gibco) and grown at 37°C with 5% CO_2_.

### Cloning and transduction of CHSTs

The DNA constructs encoding *CHST1*, *CHST2*, *CHST4*, *CHST8*, *CHST9*, *Gal3ST2*, *Gal3ST3*, and *Gal3ST4* were synthesized from GeneArt Gene Synthesis (Thermo Fisher). Each gene was cloned to include 5’ NheI site and 3’ AgeI site, ligated to linearized RP172 vector^59^ with the same enzyme pair and transformed into NEB stable Competent *E. coli*. Colonies were grown in LB media containing 100 μg/mL of ampicillin (Fisher Scientific) and plasmids were isolated from these bacterial cells using GeneJET plasmid miniprep kit (Thermo Fisher). Sequences of *CHST*-encoding plasmids were verified through Sanger sequencing. Lentivirus was made for each *CHST* gene as described previously. Briefly, the RP172 plasmids containing the *CHST* genes were co-transfected with RP18 and RP19 helper plasmids into HEK293T cells after incubating with TransIT®-LT1 transfection reagent (Mirus Bio). After 3 days, viruses were harvested and concentrated using Lenti-x Concentrator (Takara Bio) and stored at -80°C in 10 uL aliquots. 200,000 of WT U937 cells were transduced with 10 uL of each virus and cultured for 3 days. Subsequently, positively transduced cells were selected using 3.3 μg/mL of Zeocin (Thermo Fisher). After zeocin selection, mAmetrine positive populations were gated and used in the binding assays.

### Construction of the RP173 vector

The mAmatrine fluorescent marker in RP172 was replaced with EGFP to create the RP173 vector. PCR was used to amplify the EGFP sequence, and it was cloned into the RP172 vector using 5’ BstBI and 3’ SalI restriction sites. The plasmid was transformed into NEB® Stable competent E. coli cells (NEB), and the isolated plasmids were verified through sanger sequencing.

### Preparation of Siglec-Fc

Construction of all human Siglec-Fc chimeric proteins was described elsewhere.^17^ For mouse Siglec-Fc, each mouse Siglec gene containing 2-3 domains with pairs of forward and reverse primers (Table S5) were cloned into pcDNA5/FRT/V5-His-TOPO vector (Invitrogen), which contained a C-terminal hIgG1-Fc, TEV cleavage site on the N-terminal side of the Fc, as well as a C-terminal His 6 tag and a Strep-Tag II, and co-transfected with pOG44 vectors into Chinese Ovary Hamster cell lines after incubating with Lipofectamine LTX (Thermo Fisher) at RT for 30 mins as previously described.^17^ Subsequently, transfected cells were selected under DMEM-F12 (Gibco) media containing 10% FBS, 50 U/mL Penicillin/Streptomycin, 100 mM HEPES, and 1 mg/mL of Hygromycin (Thermo Fisher). Selected cells were cultured in DMEM- F12 media containing 5% FBS, 50 U/mL Penicillin/Streptomycin, and 100 mM HEPES for one week after cells reached 90% confluency. Finally, the supernatants were filtered through a sterile filter unit (0.22 μm pore size, Thermo Fisher) and stored at 4°C.

### MS analysis of glycome of each sulfotransferase transduced U937 cells

Freeze-dried cell pellets were suspended in 18 MΩ·cm H_2_O via sonication, transferred to pre-weighted screw capped cryo-vials (Sarstedt) and lyophilized. Samples were weighed, dissolved in a denaturing buffer consisting of 40 mM dithiothreitol (DTT; Sigma) and 0.5% sodium dodecyl sulfate (SDS; Sigma), and boiled for 10 mins. After cooling, 10% NP-40 was added to a final concentration of 1% and samples were subsequently treated with PNGaseF for 16 h at 37°C. Proteins were precipitated by adding three volumes of -20 °C ethanol and incubation at -20°C for 30 min prior to centrifugation (16,000 rcf, 10 min, room temperature). The upper, *N*-glycan-containing layer was collected and dried using a SpeedVac centrifugal concentrator (Thermo Fisher). The dried samples were redissolved in H_2_O and loaded onto ENVICarb solid-phase extraction cartridges (250 mg bed volume; Supelco) that had been previously conditioned by with 3 mL 80% acetonitrile (I) containing 0.1% trifluoroacetic acid (TFA) followed by 6 mL H_2_O. Samples were passed through the cartridge dropwise, using a minimal amount of positive pressure, after which the SPE cartridge was washed with water (5 mL) and subsequently eluted with 50% ACN/0.1% TFA (2 x 2.2 mL). N-glycan-containing fractions were partially concentrated using a SpeedVac before snap freezing in liquid nitrogen and lyophilization in a screw-capped tube. Glycans were reduced by dissolving them in 50 mM NH_4_OH containing 1 M NaBH_4_ for 2 h at 65°C after which the reactions were quenched by carefully adding glacial acetic acid until noticeable fizzing stopped. Samples were again desalted by SPE (exactly as described above) and reconstituted in 200 μL H_2_O.

Liquid chromatography was done using an Agilent 1290 Infinity system, supplied with 1290 Infinity binary pump, 1290 Infinity autosampler and a 1290 Infinity temperature-controlled sample compartment (Agilent Technologies, Santa Clara, CA, USA). Mass spectrometry was performed using an Agilent 6530 Quadrupole Time-of-Flight (QToF) with jet stream electrospray ionization (ESI) source. A Hypercarb™ (Thermo-Fisher; 100 mm × 2.1 mm; particle size-3 µm) HPLC column was used for separation of glycans. Full-scan MS data were acquired in high resolution, negative ionization mode at a resolution of 6,600 at a *m*/*z* of 301.9981 and 12,151 at a *m*/*z* of 2233.9120; scanning was performed at a rate of 3 Hz from *m*/*z* 100 – 3200. Reference ions at *m/z* 112.9855, 966.0007 and 2533.8923 were used, with a detection window of ± 10 ppm and a minimum height of 1000 counts. Information on MS parameters and gradients are presented below in the tables.

MS data were analysed using Agilent Technologies’ MassHunter software (version B.07.00); specifically, the Find–by-Formula (FBF) algorithm was used to obtain the total area under a chromatographic peak. The FBF algorithm takes into consideration the mass accuracy of monoisotopic masses (*m/z*), in addition to the height and spacing of the different isotopic peaks associated with the given formula, and the retention times to give the area results along with a (confidence) score. A CSV-format database of potential *N*-glycan formulae was created in Microsoft Excel, compiling a list of glycans contain between 3 and 12 hexoses (Hex), 2 and 9 hexososamines (HexNAc), 1 to 4 Neu5Ac residues, 1 to 3 deoxyhexoses (assumed to be fucoses), and 1 to 2 sulfate moieties. All putative glycan formulae contained a reduced reducing end and the isobaric combinations of Hex/Neu5Ac *vs*. Fuc/Neu5Gc were assumed to be the latter. Extracted ion chromatograms were generated with a ± 10 ppm mass window and Gaussian smoothing.

### Lectin microarray

Cell lysates from the WT and CHST transduced U937 cells were centrifuged (18,000 rcf, 10 min). The supernatant was transferred to a clean tube and protease inhibitor cocktail (Thermo Fisher) was added (1% v/v). Samples were then diluted to 2 mg/mL in PBS (pH 7.4) and labeled with NHS-activated AF647 (Thermo Fisher) as described by Pilobello *et al*.^60^ Reference sample was prepared by mixing equal amounts of protein from each sample and labeling with NHS-activated AF555 (Thermo Fisher). Printing, hybridizing, and data analysis were performed as described by Pilobello *et al*.^60^

### Mutagenesis of the two O-glycosylation sites on CD33

The original CD33-Fc DNA template^17^ was amplified with a forward primer (5’-AGCAGCGCTAGCATGCCGCTGCTGCTACTGCTG-3’) and a reverse primer (5’-GGAAAGATACCAGTGGCTGGGTTCTGTGGAAC-3’) to create the T239A mutation. The PCR product (to be used as a megaprimer) was gel-purified, and concentrated into 50 μL of elution buffer, using the GeneJET Gel Extraction Kit (Thermo Fisher). The CD33-Fc template was amplified again using 3 μL of the megaprimer and a reverse primer (5’-AGCAGCACCGGTATGAACCACTCCTGCTCTGG-3’). The resulting PCR product was gel-purified, digested with NheI and AgeI, and ligated into a linearized pcDNA5/FRT/V5-His-TOPO vector containing C-terminal hIgG-His_6_-StreptagII. After transformation into DH5α competent cells (NEB), the colonies were selected on LB agar containing 100 μg/mL Ampicillin and plasmids from several colonies were isolated for Sanger sequencing. Next, the DNA of CD33-Fc with T239A was amplified with a forward primer (5’-AGCAGCGCTAGCATGCCGCTGCTGCTACTGCTG-3’) and a reverse primer (5’-AAGTACAGGTTCTCACCGGC-3’) to mutate T260 into Ala. The resulting megaprimer was gel-purified and concentrated into 50 μL of the elution buffer. The purified megaprimer and a reverse primer (5’-AGCAGCACCGGTTCACTTCTCGAACT-3’) were used to amplify CD33-Fc containing T239A. The final product was gel-purified, digested with NheI and AgeI, and ligated into a linearized pcDNA5/FRT/V5-His-TOPO vector. The ligated plasmids were transformed into DH5α competent cells and cultured on LB-agar containing 100 μg/mL Ampicillin. Several colonies were selected and cultured, and plasmids were isolated and sequenced by Sanger sequencing. The pcDNA5 plasmids containing CD33-Fc with T239A and T260A were transfected into CHO Lec-1 cells as previously described.^17^

### Synthesis of mono- and di-sulfated LacNAc with a propylamine linker

All commercial materials used were reagent grade as supplied except where noted. Anhydrous dichloromethane (DCM) was obtained from a solvent purification system. Anhydrous dimethylformamide (DMF) was obtained from a Sure/Seal™ container. Analytical thin-layer chromatography (TLC) was performed using silica gel 60 F254 glass plates. Compound spots were visualized by UV light (254 nm) and by staining with ninhydrin in ethanol or a yellow solution containing Ce(NH_4_)_2_(NO_3_)_6_(0.5 g) and (NH_4_)_6_Mo_7_O_24_·4H_2_O (24.0 g) in 6% H_2_SO_4_ (500 mL). Glycosylation reactions were performed in the presence of molecular sieves, which were flame-dried right before the reaction under high vacuum. Glycosylation solvents were dried using a solvent purification system and used directly without further drying. Flash column chromatography was performed on silica gel 60 (230-400 Mesh). NMR spectra were recorded on Agilent DDR2 500 MHz NMR spectrometers at 298 K and referenced using residual CHCl_3_ for ^1^H-NMR (δ 7.26 ppm), and CDCl_3_ for ^13^C-NMR (δ 77.0 ppm). High resolution mass spectra were recorded on a Waters Xevo G2-XS Q-tof quadrupole mass spectrometer. See Supplementary Material for full details on the synthesis and characterization of these compounds.

### Synthesis of sialylated mono- and di-sulfated LacNAc with a propylamine linker

CMP- Sialic acid and α2-3-Sialyltransferse, PmST1, were added to a solution of each LacNAc scaffold (Figure S3) in 100 mM Tris-HCl buffer containing 20 mM MgCl_2_ (pH 8.5). Each reaction mixture was incubated at 37 °C and monitored by thin layer chromatography (TLC) (iPrOH:NH_4_OH:H_2_O, 5:2:1, v:v:v) until the reaction was completed. The reactions were terminated by dilution with a 4- fold of 100% ethanol and put at −20 °C for an hour to precipitate the enzyme. The precipitated protein was centrifuged (3700 rcf, 15 min, 4 °C), the supernatant carefully decanted into a round bottom flask, and evaporated. The residual compound **7** was resuspended in water and purified on a P2 column equilibrated in 20% NH_4_OH to yield a white compound with 66% yield. The residual compounds **11**, **14**, and **18** were purified on a HILIC HPLC column and the fractions containing each compound were lyophilized giving the desired compounds with 35%, 64%, and 35% yield respectively. The final products were characterized by ^1^H on a 700 MHz NMR, which is available in Supplementary Materials.

### Preparation of soluble Siglec fragments for mass spectrometry

Siglec-Fc proteins were purified by Histrap™ Excel (Cytiva) according to manufacturer guidelines and dialyzed into a PBS buffer overnight. The dialyzed protein was subsequently purified by Hitrap™ Protein G (Cytiva) as manufacturer guidelines and 2 mL of the elution fractions were pooled together and adjusted at pH 7 for TEV and Endo-H digestion for 3 hr at 37°C and overnight at 4°C. The reaction mixture was diluted with 10 mL of PBS buffer, passed through a Histrap™ Excel column. The flow through was collected for further dialysis three times with 200 mM of ammonium acetate at 4°C. The proteins were concentrated with Amicon (cutoff 10 kDa, Sigma) to achieve around 0.200 mg/mL of protein solution for mass spectrometry.

### Staining method for Flow-cytometry assay

Cells on a 96-microwell plate were washed with PBS twice. Siglec-Fc were used at approximately 20 μg/mL and AF647 Goat anti-hIgG (Clone: HP6017, 0.4 μg/mL, Biolegend) were mixed together in a 1:1 volume ratio right before staining, and 100 μL of the mixture was used for staining each well. After incubation on ice for 30 min, the stained cells were washed twice with PBS, resuspended in PBS, and analyzed using Fortessa X- 20 Flow cytometer (BD Biosciences). Flow cytometric data were analyzed using FlowJo v10 software.

### Glycosylation or Sulfation inhibitors

500,000 of the WT, CHST1, and CHST2 overexpressing U937 cells were plated onto 6-well plates and incubated at 37 °C in 2 mL of RPMI media containing 10% FBS, 50 U/mL of Penicillin/Streptomycin, and 100 mM of HEPES with kifunensine (2.5 μg/mL, Toronto Research Chemicals) or benzyl-α-GalNAc (2.0 μg/mL, Sigma) or Genz123346 (5 μg/mL, Toronto Research Chemicals) for 72 hours. The same number of the WT, *CHST1*, and *CHST2* cells were treated with 25 mM of NaClO_3_ for 48 hours to inhibit cellular carbohydrate sulfation.

### N-glycan extraction

N-glycans were prepared from 10 million WT U937 cells and the *CHST* overexpressing U937 cells. Briefly, Cells were lysed, trypsinized and the following peptide mixtures were treated with PNGase-F as described previously.^12^

### Analysis of glycopeptides by LC-MS/MS

Proteins were reduced with 10 mM dithiothreitol at 37°C for 1 h and then alkylated with a final concentration of 50 mM iodoacetamide in 25 mM ammonium bicarbonate buffer for 1 h in the dark at room temperature. After that, dithiothreitol was added to a final concentration of 50 mM and proteins were digested overnight with sequencing grade trypsin (Promega) at an enzyme-to-substrate ratio of 1:50 at 37 °C. The digested products were then diluted with formic acid to a final concentration of 0.1% and cleaned up using ZipTip C18 (Merck Millipore). Data was obtained on an Orbitrap Fusion Lumos Tribrid mass spectrometer (Thermo Fisher Scientific) with an Easy-nLC 1200 (Thermo Fisher Scientific). Peptide mixtures were redissolved in 0.1% formic acid (Solvent A) and loaded onto an Acclaim PepMap RSLC 25cm x 75 µm i.d. column (Thermo Fisher Scientific) and separated at a flow rate of 300 nl/min using a gradient of 5% to 40% solvent B (80% acetonitrile with 0.1% formic acid) in 60 min. The mass spectrometer was operated in the data-dependent mode. The full scan MS spectra were acquired from 400 to 1800 m/z in the orbitrap with the resolution set to 120000. The highest charge state ions were sequentially isolated for MS^2^ analysis using the following settings: HCD MS^2^ with AGC target at 5×10^4^ and were detected in the orbitrap at a resolution of 30000, isolation window 2, step collision energy (%): 25, 28, 30. An HCD-pd-EThcD workflow was employed to additionally trigger EThcD events upon detecting at least one of the HCD product ions at m/z 204.0867, 138.0545, or 366.1396. The calibrated charge-dependent parameter was used, and supplemental activation was allowed with a 15% SA collision energy. The EThcD ions were detected in the orbitrap at a resolution of 60000. The HCD and EThcD MS2 data were processed by the Byonic software (Protein Metrics Inc., v 3.9.6) and tentatively identified O-glycopeptides were manually verified by examining the raw data.

### Statistical analyses

All statistical analyses in this study were performed using GraphPad Prism version 7 software. For experiments comparing two groups, an unpaired two-tailed Student’s *t*- test was used to evaluate statistical significance. For datasets with more than two sample groups, a one-way ANOVA with Tukey’s post hoc test was performed. For group analysis, a two-way ANOVA with either Tukey’s post hoc test or Sidak’s multiple comparisons test was performed. A non-parametric Wilcoxon matched-pairs signed-rank test was used to assess statistical significance for *CHST1* transcript expression levels in matched non-malignant adjacent tissues and their primary tumour counterparts. Statistical significance of the 5-year overall survival curves between assigned case groups were evaluated by Gehan-Breslow-Wilcoxon test. Data were considered statistically significant when *P*<0.05.

### Kd determination by ESI-MS

The affinities of the glycan ligands for the CD33 fragment were measured by ESI-MS, performed in positive ion mode using a Q Exactive Orbitrap mass spectrometer (Thermo Fisher Scientific, U.K) with a modified nanoESI source. Submicron nanoESI emitters were used to minimize nonspecific binding of glycans to the protein during the ESI process.^61^ The emitters were produced from borosilicate capillary glass (1.0 mm outside diameter (o.d), 0.69 mm inner diameter (i.d), 10cm length, Sutter Instruments, CA) using a P-100 micropipette puller. A voltage of 0.5 kV was applied to the platinum wire inserted into the back end of the emitters, in contact with the sample solution. Resolution power of 17500 at m/z of 200 was used. The automatic gain control target (AGC), the maximum injection time, capillary temperature and S-lens RF level were set to 1 × 106, 100 ms, 150 °C and 100, respectively. Data acquisition and processing were carried out using Xcalibur (Thermo Fisher, version 4.1). For all measurements, a reference protein (P_ref_) was added to the solution to confirm the absence of nonspecific glycan binding.^62^ To measure the dissociation constant (*K*_d_) for each protein-ligand complex, a titration approach was used. The initial protein ([P]_0_) was kept constant while the initial ligand concentraion ([L]_0_) varied. The *K*_d_ was then obtained by fitting Eq 1 to the fraction of bound protein (*R/(R* + 1)) versus [L]_0_:

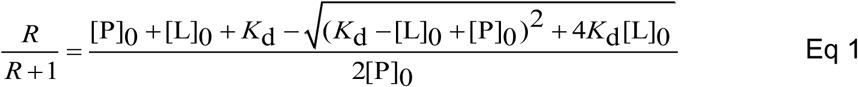

where *R* is the ratio of total ion abundance (Ab) of ligand-bound (PL) to free protein (P), Eq 2:

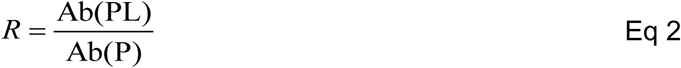

### Liposome binding assay

EV, CHST1, CHST2, and CHST1/2-overexpressing U937 cells were resuspended in media, and 100,000 cells were added to a 96-well U-bottom plate in 200 μL RPMI containing 10% FCS. Cells were centrifuged (300 rcf, 5 min) and the supernatant was discarded. The cell pellet was re-suspended in 50 μL of fresh media and 50 μL of media containing naked (without CD33 ligand) or CD33 ligand (CD33L) tagged liposomes was added to it. The final concentration of liposomes was 100 μM (total concentration of lipid). The CD33L concentration on the liposomes was 3.33 mol %, all liposomes contained 0.1 mol % AF647-conjugated lipid, and were prepared as described previously.^17^ These suspensions were incubated for 60 min at 37 °C . Following this incubation, 100 μL of media was added to each sample and they were centrifuged (300 rcf, 5 min). The supernatant was discarded, and the cell pellet was suspended in a flow buffer and further analyzed by flow cytometry.

### Measurement of cell viability

Adherent cell lines (MDA-MB-231, MDA-MB-468, MCF-7, SF-295, SNB-75 were trypsinized and resuspended into RPMI growth media (10% FBS, 50 U/mL penicillin/streptomycin, 100 mM HEPES). Non-adherent cell lines (U937, K562, and CCRF-CEM) were directly resuspended into the RPMI growth media. A half million of each cancer cell line was plated onto a 12-well plate with 2 mL the RPMI growth media containing 25 mM NaClO_3_ and cultured at 37°C with 5% CO_2_ for 48h. Subsequently, the adherent cell lines were incubated with PBS containing 1 mM EDTA for 5 min and carefully taken off from the surface by pipetting up and down. The non-adherent cell lines were moved into 15 mL tube. After washing the cells with PBS twice with centrifugation (300 rcf, 5 min), the cells were incubated with PBS containing 2 μg/mL propidium iodide (PI) on ice for 10 mins. The cell viability was measured by flow cytometry by assessing PI negative population.

## Acknowledgments

This study was supported by grants from the Natural Sciences and Engineering Research Council of Canada (RGPIN-2018-03815) and the Canadian Glycomics Network (CD-41 and CR-03), as well as a Tier II Canada Research Chair in Chemical Glycoimmunology to M.S.M. J.J. is supported by the Alberta Graduate Excellence Scholarship. J.R.E. is funded by fellowships from the Alberta Innovates Graduate Student Scholarships and Canadian Arthritis Society. E.R. is funded by a fellowship from the Alberta Innovates Graduate Student Scholarship. The work of L.K.M. and P.R. is funded by the Canada Excellence Research Chair in Glycomics (to L.K.M.). Dr. Peng Wu (Scripps) and Dr. Warren Wakarchuk (University of Alberta) are thanked for the PmST1 and PNGase-F clone, respectively. Mass spectrometry data for glycopeptides were acquired at the Academia Sinica Common Mass Spectrometry Facilities for Proteomics and Protein Modification Analysis (supported by grant AS-CFII-108-107).

