## Supplementary Information for "Carbohydrate sulfation as a mechanism for fine-tuning Siglec ligands"

**Table of Contents**

**Table S1. The known substrates for each *CHST*.** S3

**Table S2. Parameters for QToF-MS in negative electro-spray ionization (ESI) mode.** S4

**Table S3. MS data by Agilent Technologies' MassHunter software (version B.07.00)** S5

**Table S4. RNA expression of *CHST*s in the selected cancer cell lines.** S6

**Table S5. Forward and reverse primers for cloning mouse Siglec-Fc.** S7

**Figure S1. Full heatmap of lectin microarray analysis for U937 Cells.** S8

**Figure S2. mSiglec binding to the *CHST* transduced cells.** S9

**Figure S3. Chemical synthesis of sulfated LacNAc.** S10

**Figure S4. Comparison of the T239A/T260A CD33 fragment with the WT CD33 fragment.** S11

**Figure S5. CD33L-liposome binding assay on the *CHST1*, *CHST2*, and *CHST1/2* overexpressing U937 cells.** S12

**Figure S6. Cell viability assay with propidium iodide under $NaClO_3$ treatment.** S13

**Figure S7. $NaClO_3$ treatment on the cell lines expressing a low level of *CHST1* and *CHST2*.** S14

**Figure S8. The effect of $NaClO_3$ on SNA or MAA in several cancer cell lines.** S15

**General methods for chemical synthesis.** S16-S23

**NMR spectra.** S24-S61

**Reference.** S62

**Table S1. The known substrates for each *CHST*.**

| Gene | HGNC ID | Substrate | Reference |
|---|---|---|---|
| ***CHST1*** | **HGNC:1969** | 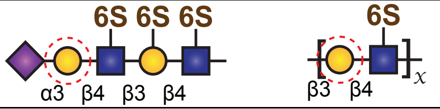  | 1 |
| ***CHST2*** | **HGNC:1970** | 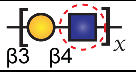  | 2 |
| ***CHST4*** | **HGNC:1972** | 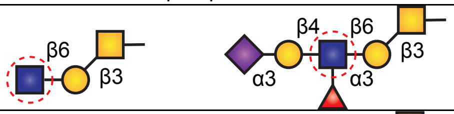  | 3 |
| ***CHST8*** | **HGNC:15993** | 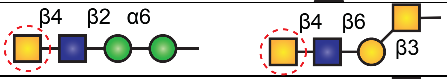  | 4 |
| ***CHST9*** | **HGNC:19898** | 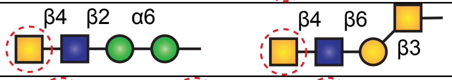  | 5 |
| ***Gal3ST2*** | **HGNC:24869** | 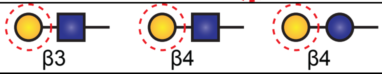  | 6 |
| ***Gal3ST3*** | **HGNC:24144** | 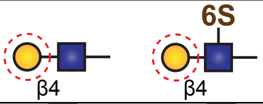  | 7 |
| ***Gal3ST4*** | **HGNC:24145** | 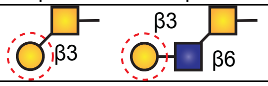  | 8 |

**Table S2. Parameters for QToF-MS in negative electro-spray ionization (ESI) mode.**

| Parameter (units) | Value |
|---|---|
| Sheath gas flow(L/min) | 10 |
| Sheath gas temperature (°C) | 350 |
| Drying gas flow (L/min) | 10 |
| Drying gas temperature (°C) | 300 |
| Nozzle voltage (V) | 1000 |
| Nebulizer (psi) | 35 |
| Fragmentor voltage (V) | 175 |

Gradient conditions used in uHPLC for separation of glycan.

| Time (min) | Mobile phase B (%) | Flowrate (μL/min) |
|---|---|---|
| 0.0 | 0 | 250 |
| 5.0 | 5 | 250 |
| 15.0 | 20 | 250 |
| 20.0 | 40 | 250 |
| 25.0 | 80 | 250 |
| 27.0 | 80 | 250 |
| 27.1 | 5 | 250 |

* Mobile phase A was ultrapure water+0.1% formic acid; mobile phase B was ACN+0.1% formic acid.

**Table S3. MS data by Agilent Technologies' MassHunter software (version B.07.00)**

| glycan code[(1)] | RT[(2)] (min) | relative peak area (%) [(3)] | | | | | | | | |
|---|---|---|---|---|---|---|---|---|---|---|
| | | WT[(4)] | CHST1 | CHST2 | CHST4 | CHST8 | CHST9 | Gal3ST2 | Gal3ST3 | Gal3ST4 |
| 25000 | 13.9 (0.08) | 5.7 | 7.0 | 5.5 | 6.4 | 5.9 | 6.0 | 6.9 | 6.0 | 8.2 |
| 26000 | 12.3(0.08) | 20.6 | 10.2 | 10.7 | 16.4 | 20.6 | 20.6 | 33.4 | 26.1 | 26.6 |
| 27000_1 | 12.1(0.1) | 7.8 | 8.6 | 7.1 | 7.0 | 7.2 | 6.6 | 9.2 | 6.6 | 6.2 |
| 27000_2 | 12.3(0.07) | 3.8 | 3.7 | 4.0 | 5.7 | 6.8 | 6.8 | 10.6 | 8.0 | 7.7 |
| 28000_1 | 12.1(0.09) | 21.1 | 22.1 | 21.2 | 20.3 | 19.1 | 18.5 | 25.7 | 18.6 | 17.9 |
| 28000_2 | 12.9(0.00) | 0 | 0 | 0 | 0 | 0 | 0 | 0.6 | 0.6 | 0.4 |
| 29000 | 12.1(0.09) | 21.9 | 28.5 | 31.0 | 24.6 | 24.7 | 25.9 | 0 | 25.4 | 23.6 |
| 2(10)000 | 12.5(0.13) | 1.8 | 3.7 | 4.3 | 3.0 | 2.6 | 2.5 | 3.1 | 2.0 | 1.3 |
| 34000_1 | 14.6 | 0 | 0 | 0.4 | 0 | 0 | 0 | 0 | 0 | 0 |
| 34000_2 | 12.6(0.00) | 0 | 0 | 0 | 0 | 0 | 0 | 0 | 0 | 0.3 |
| 35000 | 12.2(0.06) | 0.5 | 0.7 | 0 | 0.6 | 2.0 | 1.0 | 1.0 | 0 | 0.8 |
| 45000 | 13.4(0.07) | 3.2 | 3.8 | 4.0 | 0 | 0 | 2.7 | 0 | 0 | 2.5 |
| 63000 | 14.2(0.00) | 0 | 0 | 0.5 | 1.4 | 1.6 | 0 | 0 | 0 | 0 |
| 45100 | 14.7(0.07) | 9.0 | 4.0 | 4.4 | 1.3 | 4.1 | 2.5 | 1.6 | 0 | 1.8 |
| 45120 | 15.1(0.13) | 1.4 | 0.5 | 0 | 0.3 | 0 | 0.6 | 0 | 0.1 | 0 |
| 53100 | 12.2(0.13) | 0 | 0 | 0.1 | 2.1 | 0 | 0 | 0.9 | 0.2 | 0.03 |
| 56010 | 18.2(0.00) | 0.6 | 0 | 0 | 0 | 0 | 0.5 | 0.7 | 0 | 0.18 |
| 56030 | 12.4(0.1) | 0 | 0 | 0 | 1.4 | 1.1 | 0 | 0 | 0.4 | 0.3 |
| 56130_1 | 12.2(0.07) | 1.8 | 1.7 | 1.2 | 1.2 | 0 | 0 | 1.6 | 2.1 | 0 |
| 56130_2 | 11.2(0.00) | 0.2 | 0 | 0 | 0.4 | 0 | 0 | 0.3 | 0 | 0.03 |
| 73100_1 | 26.3(0.00) | 0.3 | 0 | 0 | 0.4 | 0.6 | 0.5 | 0 | 0 | 0.2 |
| 73100_2 | 12.5(0.00) | 0.2 | 0 | 0 | 0.9 | 0 | 0.8 | 1.0 | 0 | 0 |
| 34001 | 21.8(0.06) | 0 | 2.4 | 0 | 0 | 0 | 0 | 3.3 | 2.2 | 0 |
| 44001 | 12.8(0.8) | 0 | 0 | 2.0 | 0 | 0 | 0 | 0 | 0.9 | 0.6 |
| 53101_1 | 14.0(0.00) | 0 | 0.6 | 0.3 | 0 | 0 | 0.5 | 0 | 0 | 0 |
| 53101_2 | 12.7(0.8) | 0 | 0 | 0 | 1.6 | 1.3 | 1.2 | 0 | 0 | 0.4 |
| 53101_3 | 12.6(5.0) | 0 | 0 | 0.4 | 0 | 0 | 0 | 0 | 0 | 0.2 |
| 54101_1 | 12.1(0.05) | 0 | 1.1 | 0 | 0 | 0 | 1.1 | 0 | 0 | 0 |
| 54101_2 | 12.3(0.13) | 0 | 1.1 | 0 | 0 | 1.5 | 1.2 | 0 | 0.5 | 0.8 |
| 54101_3 | 14.0(3.25) | 0 | 0 | 0 | 1.5 | 0.9 | 0.3 | 0 | 0 | 0.3 |
| 63101 | 12.4(0.06) | 0 | 0 | 2.7 | 3.3 | 0 | 0 | 0 | 0 | 0 |

**Notes:** (1) Glycan codes refer to the number of hexosamine (HexNAc), hexose (Hex), deoxyhexose (*i.e.* L-fucose; Fuc), 5-*N*-acetyl-*D*-neuraminic acid (Neu5Ac), N-glycolylneuraminic acid (Neu5Gc) and sulfate (S) moieties present, as deduced by the known molecular formulae for these residues. For example 632001 = $HexNAc_6Hex_3Fuc_2S_1$. Where isobars are differentiated based on their unique retention times, these are denoted with a _1, _2, *etc*. suffix. (2) Mean retention times are provided with standard deviations in parentheses. (3) Peak areas as deduced by MassHunter's Find-By-Formula algorithm are reported as a percentage of the sum of the total peak areas identified in each sample. (4) The following abbreviations are used: WT = wild type; CHST = carbohydrate sulfotransferase; and Gal3ST = galactose-3-*O*-sulfotransferase.

**Table S4. RNA expression of *CHST*s in the selected cancer cell lines.**

| Cell lines | CHST1 | CHST2 | CHST3 | CHST4 | CHST5 | CHST6 | CHST7 | CHST8 |
|---|---|---|---|---|---|---|---|---|
| **MCF7** | 0.205 | 0 | 0.309 | 0.274 | 0.384 | 0.068 | 0.491 | 1.432 |
| **MDA-MB-231** | 0.067 | 1.832 | 0.354 | 0.531 | 0.046 | 0.016 | 2.326 | 0.06 |
| **SF-295** | 2.662 | 0.682 | 2.571 | 0 | 0.181 | 0.2 | 1.493 | 0.439 |
| **SNB-75** | 1.847 | 2.929 | 3.161 | 0.081 | 0.228 | 0.924 | 1.025 | 0.653 |
| **U251** | 2.451 | 1.828 | 3.629 | 0 | 0.105 | 0.257 | 2.135 | 0.106 |
| **K-562** | 0.061 | 2.617 | 0.064 | 0 | 0 | 0 | 0.143 | 0.094 |
| **A549** | 0.11 | 0.054 | 1.367 | 0.147 | 0.158 | 0.088 | 1.124 | 0.366 |

**Table S5. Forward and reverse primers for cloning mouse Siglec-Fc.**

| Name | Forward primer | Reverse primer |
|---|---|---|
| **mSiglec-1** | agcagcgctagcatgcacctgggcactgggatg | agcagcaccggtaaggggtggatatctgac |
| **mCD22** | agcagcgctagcatgcgcgtccattacctgtgg | agcagcaccggtcctggagggttctggagc |
| **mCD33** | agcagcgctagcatgctgtggccactgccg | agcagcaccggtgcctgatttccgggtaac |
| **mSiglec-4** | agcagcgctagcatgatattcctcgccacc | agcagcaccggtcgtcccattcactgtggg |
| **Siglec-E** | agcagcgctagcatgctgctgttgctgctg | agcagcaccggtagacaggctcaaggaaatg |
| **Siglec-F** | agcagcgctagcatgcggtgggcatggctg | agcagcaccggttcccagcagctgtagaag |
| **Siglec-G** | agcagcgctagcatgtcactgctgctgttc | agcagcaccggtagtcactctcaggtcctgtg |
| **Siglec-H** | agcagcgctagcatgctgctgtcccgactgctg | agcagcaccggtctgtgaattataggtgac |
| **mSiglec-15** | agcagcgctagcatggaggggtccctccaactc | agcagcaccggtggcgccgtggaagcggaac |

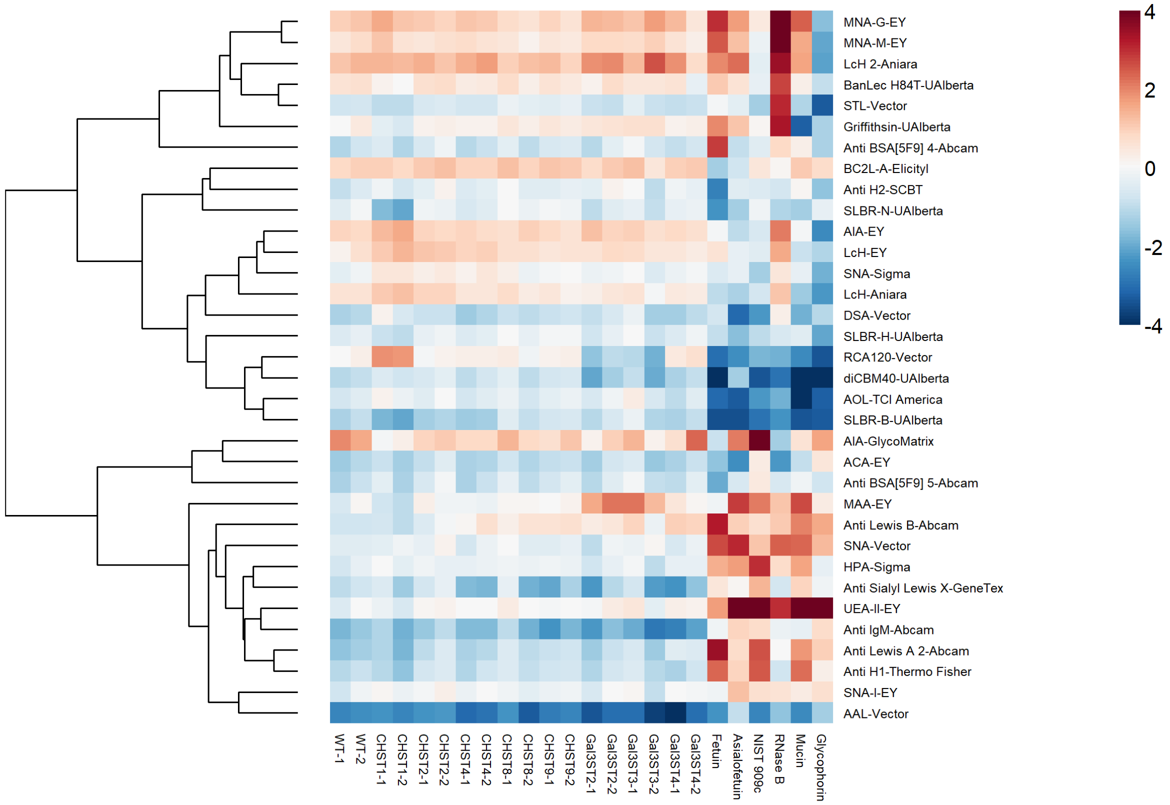


**Figure S1. Full heatmap of lectin microarray analysis for U937 Cells.** Lysates from each cell line were labeled with AlexaFluor 647 and run against an equal amount (by protein) of an orthogonally labeled pooled reference (labeled with AlexaFluor 555). Each cell line was run in biological duplicate. Median normalized $\log_2$ ratios (Sample (S)/Reference(R)) of all lectins passing quality control are shown.

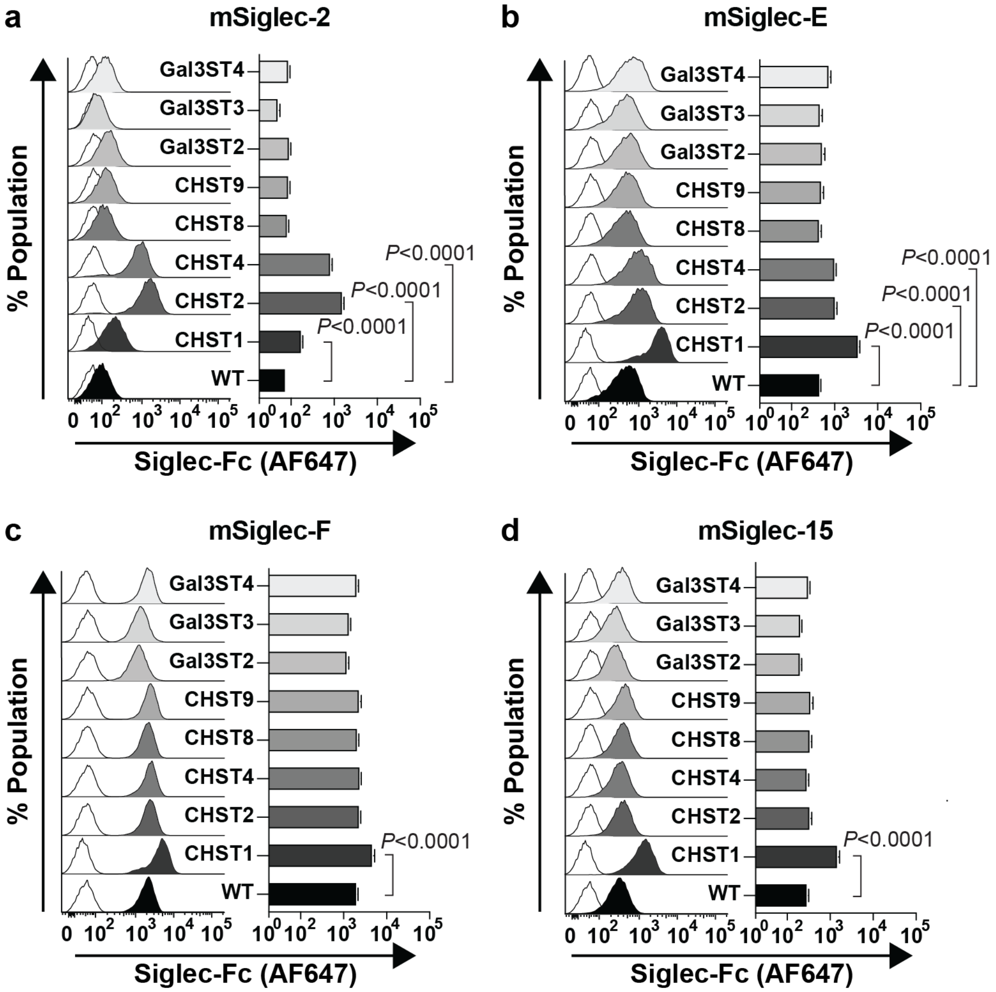


**Figure S2. mSiglec binding to the *CHST* transduced cells.** (a) mCD22, (b) mSiglec-E, (c) mSiglec-F, and (d) mSiglec-15. Data plots are presented as mean ± SD. Statistical significance was calculated using a one-way ANOVA with Tukey's multiple comparisons test.

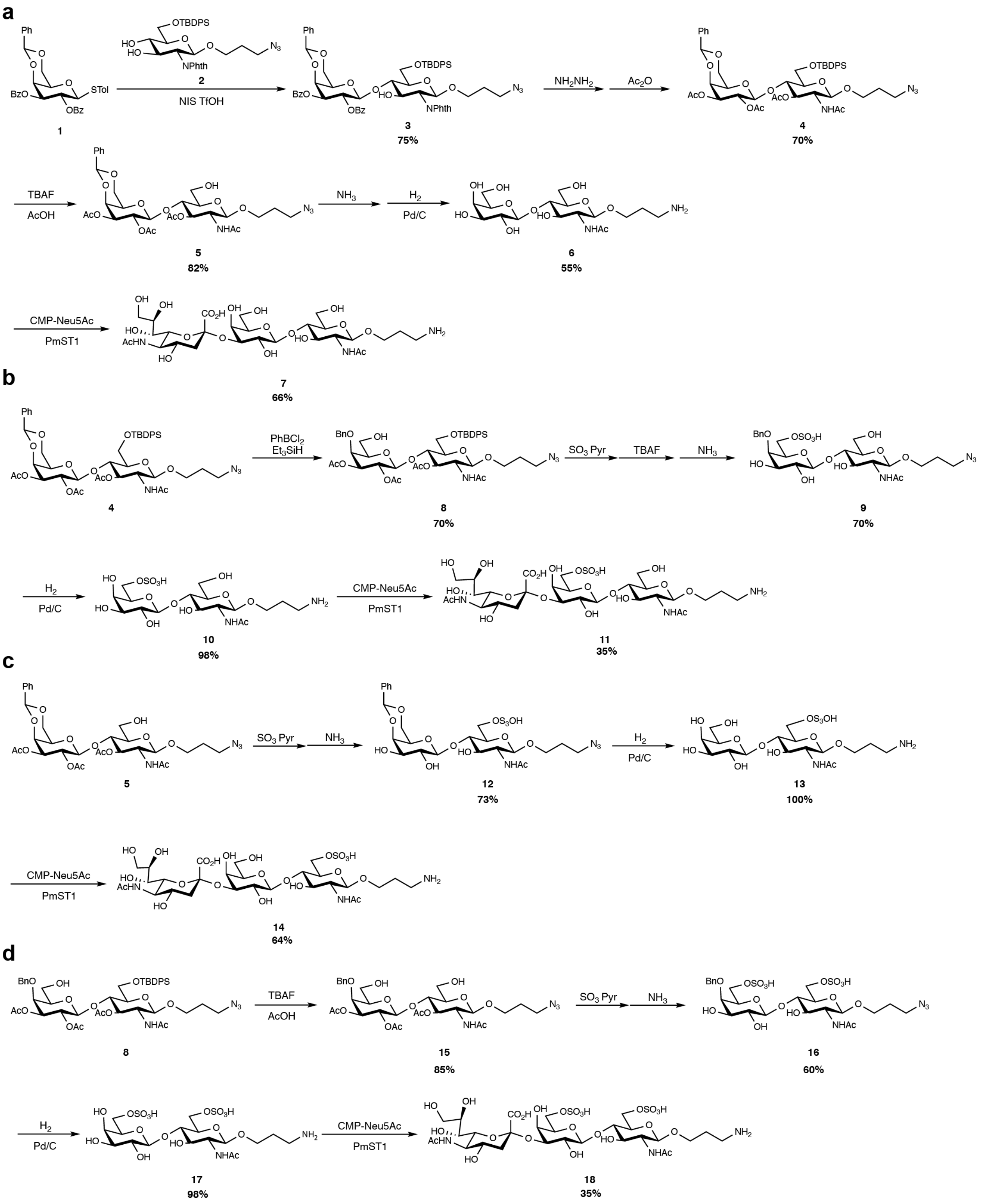


**Figure S3. Chemical synthesis of sulfated LacNAc.** (a) 3SLN **7**, (b) 6-S-3SLN **11**, (c) 6'-S-3SNL **14**, and (d) 6,6'-S,S-3SLN **18**. Compound 1 and 2 were synthesized as described in the literature.[9,10]

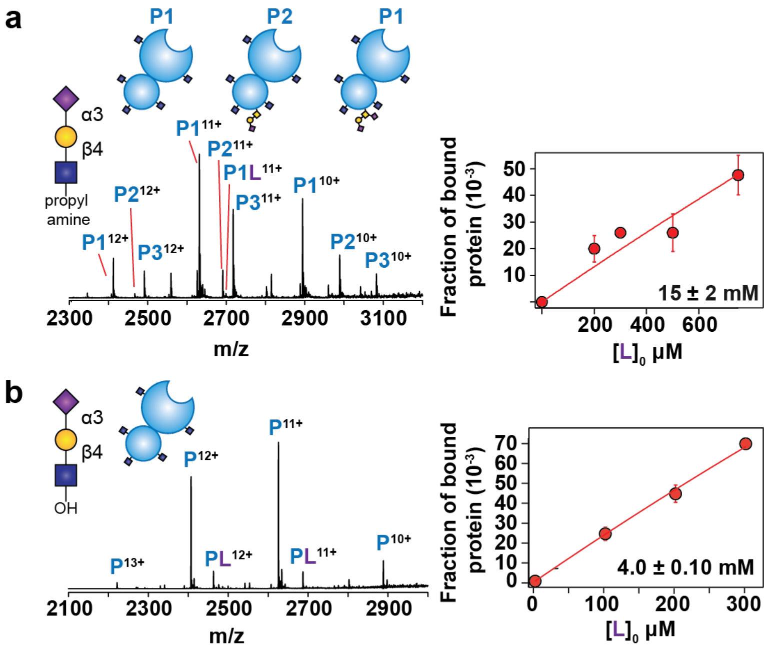


**Figure S4. Comparison of the T239A/T260A CD33 fragment with the WT CD33 fragment.** (a) $K_d$ determination of the WT fragment with a disialyl T antigen on 3SLN with a propyl amine linker. (b) $K_d$ determination of the T239A/T260A CD33 fragment on 3SLN without any linker.

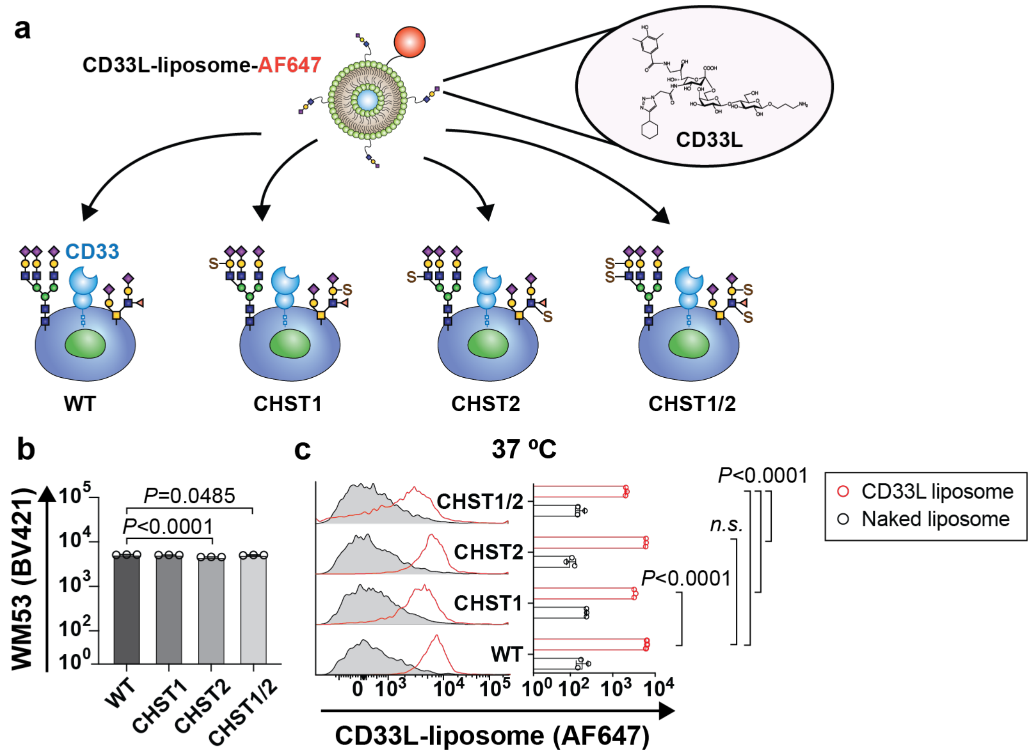


**Figure S5. CD33L-liposome binding assay on the *CHST1*, *CHST2*, and *CHST1/2* overexpressing U937 cells.** (a) Scheme for the masking assay to assess *cis* ligands of CD33. Liposome binding is measured using flow cytometry. The structure of the CD33 high affinity ligand is shown in the inset.[11] (b) Anti-CD33 (clone WM53) staining on CHST-transduced U937 cells to evaluate CD33 expression. (c) Results of the liposome assay performed at 37 ºC. Data plots are presented as mean ± SD. Statistical significance was calculated using one-way ANOVA with Tukey's multiple comparisons test.

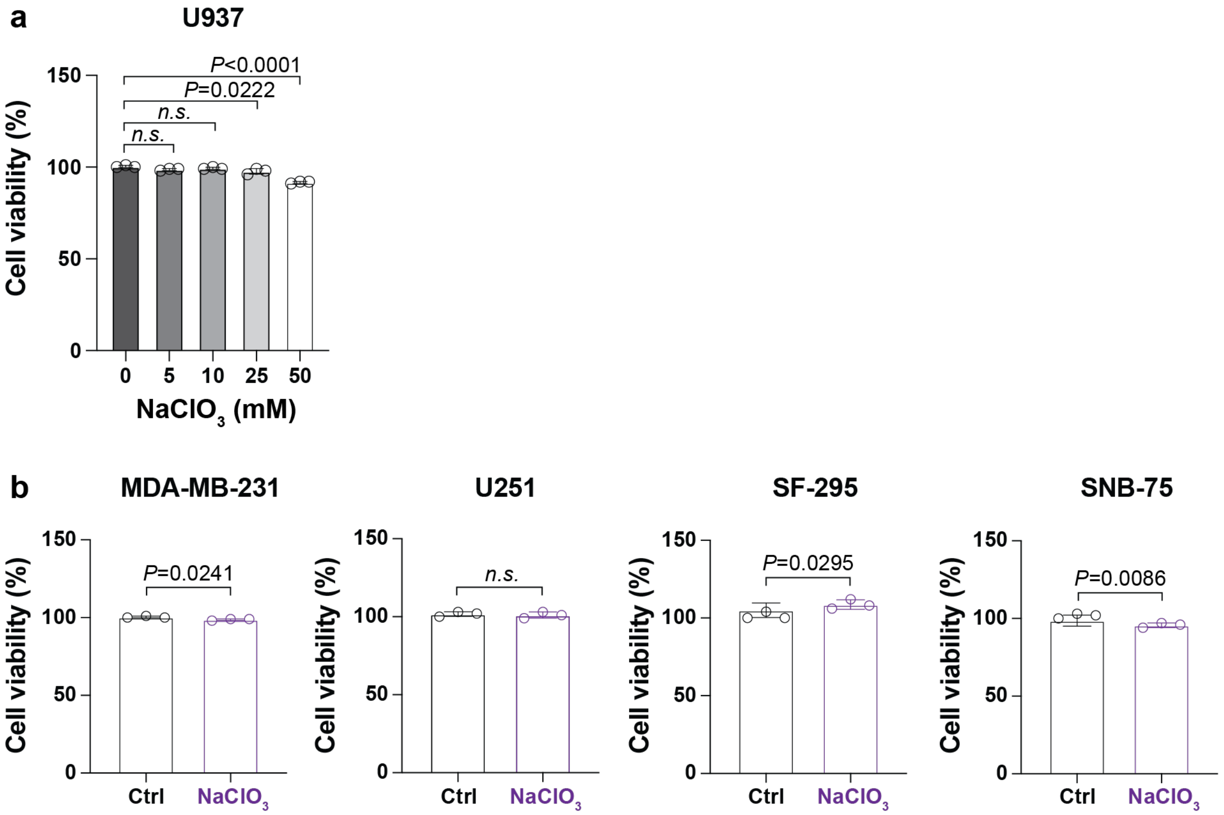


**Figure S6. Assessing cell viability following $NaClO_3$ treatment.** (a) Titration of $NaClO_3$ to assess the effect of $NaClO_3$ on cell viability in U937. (b) Cell viability assay under 25 mM of $NaClO_3$ in MDA-MB-231, U251, SF-295, and SNB-75. Data plots are presented as mean ± SD. Statistical significance was calculated using either a one-way ANOVA with Tukey's multiple comparisons test or an unpaired student's t-test.

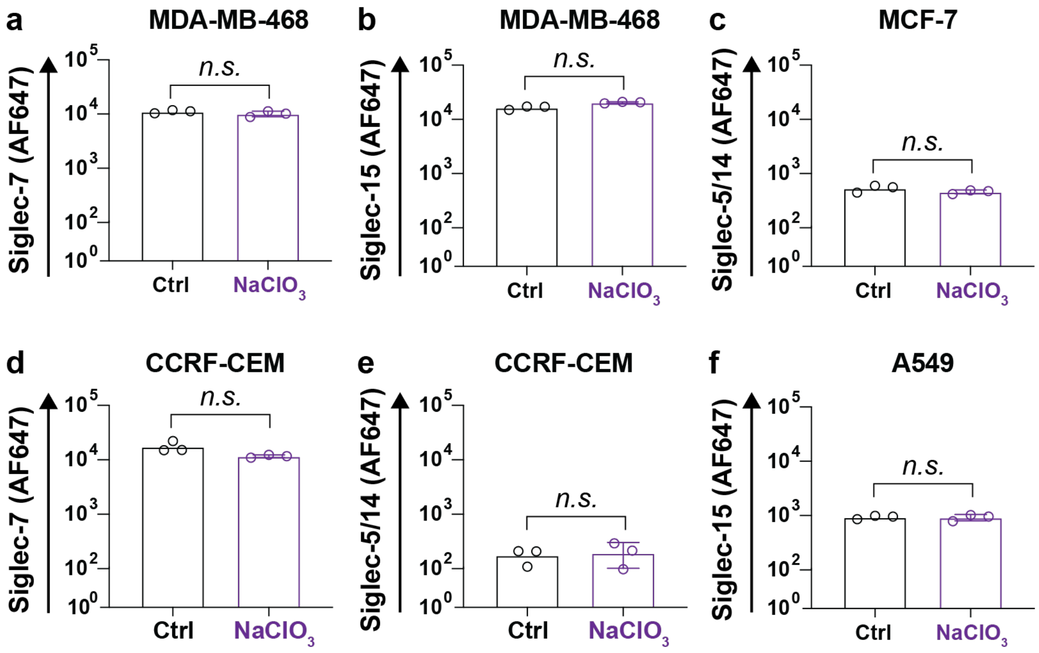


**Figure S7. Assessing Siglec-Fc binding to cell lines expressing low levels of *CHST1* and *CHST2*.** (a,b) MDA-MB-468, (c) MCF-7, (d,e) CCRF-CEM, and (f) A549 cells were probed by Siglec-Fc with and without 25 mM $NaClO_3$. Data plots are presented as mean ± SD. Statistical significance was calculated using an unpaired two-tailed Student's *t*-test.

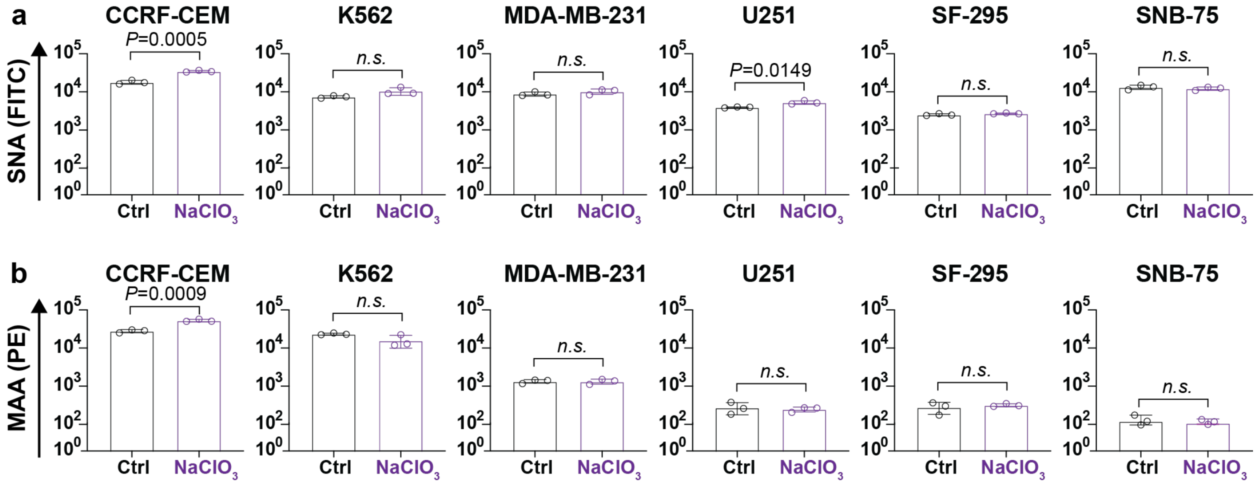


**Figure S8. Treatment of cancer cell lines with $NaClO_3$ does not alter staining with SNA or MAA.** CCRF-CEM, K562, MDA-MB-231, U251, SF-295, and SNB-75 were treated with $NaClO_3$ and probed by (a) SNA and (b) MAA, and measured by flow cytometry. Statistical significance was calculated using an unpaired t-test.

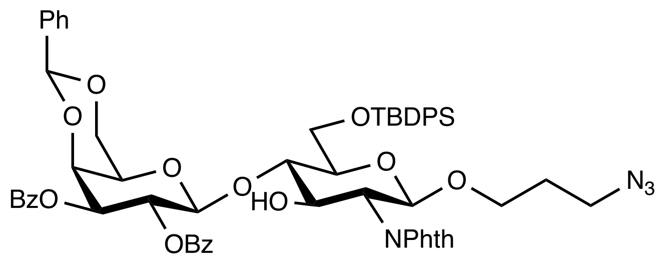


**3**

3-Azidopropyl 2,3-di-*O*-benzoyl-4,6-*O*-benzylidene-β-D-galactopyranosyl-(1→4)-6-*O*-$^t$butyldiphenylsilyl-2-deoxy-2-phthalimido-β-D-glucopyranoside **3**.

A mixture of glycosyl donor **1** (87.3 mg, 0.15 mmol), glycosyl acceptor **2** (63.0 mg, 0.1 mmol), and freshly activated molecular sieves (4 Å, 100 mg) in DCM (10 mL) was stirred under nitrogen for 30 min. The solution was cooled to −78 °C. *N*-Iodosuccinimide (NIS, 120 mg, 0.16 mmol) and TfOH (15 µL ,0.02 mmol) were added. Lumps of dry ice were removed from the acetone bath, and the reaction was allowed to slowly reach 0 °C over 2 h. Upon completion, the solids were filtered off, and the filtrate was washed with an aqueous 10% $Na_2S_2O_3$ solution. The organic layer was separated, dried over $MgSO_4$, and concentrated *in vacuo*. The residue was purified by flash column chromatography giving compound **3** (81.6 mg, 0.075 mmol) in 75% yield. $[\alpha]_D^{20}$ = +14 (c 0.1, DCM). $^1$H NMR (500 MHz, Chloroform-*d*) δ 8.01 – 7.96 (m, 2H), 7.91 – 7.69 (m, 10H), 7.53 – 7.30 (m, 15H), 7.23 – 7.15 (m, 2H), 5.90 (dd, *J* = 10.5, 8.3 Hz, 1H), 5.51 (s, 1H), 5.35 (dd, *J* = 10.5, 3.5 Hz, 1H), 5.18 (d, *J* = 8.3 Hz, 1H), 5.05 (d, *J* = 8.3 Hz, 1H), 4.59 (dd, *J* = 3.6, 1.1 Hz, 1H), 4.49 (dd, *J* = 10.8, 8.5 Hz, 1H), 4.43 (d, *J* = 12.2 Hz, 1H), 4.21 (dd, *J* = 10.8, 8.5 Hz, 1H), 3.87 – 3.69 (m, 4H), 3.66 (br, 1H), 3.48 – 3.34 (m, 3H), 3.13 (t, *J* = 6.7 Hz, 2H), 1.70 – 1.66 (m, 2H), 1.12 (s, 9H). $^{13}$C NMR (126 MHz, Chloroform-*d*) δ 168.39, 168.10, 166.08, 164.96, 137.31, 135.97, 135.55, 134.04, 133.46, 133.17, 132.72, 129.98, 129.96, 129.64, 128.97, 128.95, 128.41, 127.65, 126.20, 100.87, 100.82, 97.80, 79.25, 74.59, 73.35, 72.41, 69.38, 69.07, 68.50, 66.89, 65.59, 61.70, 56.20, 48.11, 26.93, 26.79, 19.56. ESI-MS: $[M+NH_4^+]$ $C_{60}H_{64}N_5O_{14}Si$ calculated: 1106.4213, observed: 1106.4214.

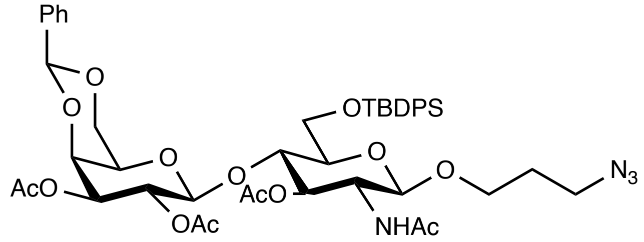


**4**

3-Azidopropyl 2,3-di-*O*-acetyl-4,6-*O*-benzylidene-β-D-galactopyranosyl-(1→4)-6-*O*-$^t$butyldiphenylsilyl-2-*N*-acetyl-3-*O*-acetyl-2-deoxy-β-D-glucopyranoside **4**.

Compound 3 (1.08 g, 1 mmol) was dissolved in a mixture of DCM: MeOH (1:1, 40 mL) and hydrazine monohydrate (64-65%, 10 mL, 0.2 mmol). The reaction was stirred overnight and monitored by mass spec till completion. The reaction mixture was then concentrated, dried *in vacuo*. The residue was dissolved in DCM (30 mL) then 1 M AcCl in DCM (20 mL, 20 mmol) was added. The reaction was stirred 2 h and monitored by TLC. Upon completion, the mixture was cooled with ice and quenched with MeOH, then concentrated *in vacuo*. The residue was purified by flash column chromatography giving compound 4 (642 mg, 0.07 mmol) in 70% yield. $[\alpha]_D^{20}$ = +8 (c 0.1, DCM). $^1$H NMR (500 MHz, Chloroform-*d*) δ 7.78 - 7.72 (m, 4H), 7.50 – 7.34 (m, 9H), 7.20 –7.14 (m, 2H), 5.70 (d, *J* = 9.5 Hz, 1H), 5.47 (s, 1H), 5.23 (dd, *J* = 10.3, 8.0 Hz, 1H), 5.03 – 4.97 (m, 2H), 4.87 (dd, *J* = 10.3, 3.6 Hz, 1H), 4.81 (d, *J* = 8.0 Hz, 1H, Gal -1H), 4.37 – 4.33 (m, 2H), 4.29 (d, *J* = 3.6 Hz, 1H), 4.23 – 4.21 (m, 2H), 4.02 (dd, *J* = 12.5, 1.7 Hz, 1H), 4.00 – 3.89 (m, 3H),

3.54 – 3.49 (m, 1H), 3.43 – 3.30 (m, 4H), 2.08 (s, 3H), 2.07 (s, 3H), 1.97 (s, 3H), 1.93 – 1.80 (m, 2H), 1.77 (s, 3H), 1.08 (s, 9H). $^{13}$C NMR (126 MHz, Chloroform-*d*) 171.96, 170.58, 170.30, 168.70, 135.96, 135.43, 133.44, 132.20, 129.93, 129.91, 128.24, 128.19, 127.90, 127.67, 126.40, 123.29, 101.63, 101.26, 100.22, 75.36,73.29, 69.12, 68.62, 66.19, 65.53, 61.27, 48.14, 26.83, 23.37, 21.07, 20.86, 20.85, 20.49. ESI-MS: [M+H$^+$] $C_{46}H_{59}N_4O_{14}Si$ calculated: 919.3791, observed: 919.3726.

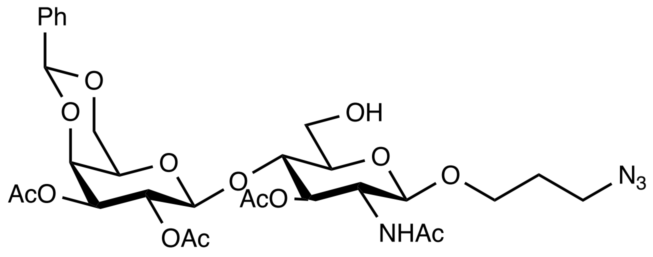


**5**

3-Azidopropyl 2,3-di-*O*-acetyl-4,6-*O*-benzylidene-β-D-galactopyranosyl-(1→4)-2-*N*-acetyl-2-deoxy -3-*O*-acetyl-β-D-glucopyranoside **5**.

Compound **4** (918 mg, 1 mmol) was dissolved in THF (10 mL). A solution composed of 1 M TBAF in THF (1.1 mL, 1.1 mmol) and AcOH (66 μL, 1.1 mmol) was added dropwise into compound **4** and the reaction was stirred overnight. Upon completion, the solution was first concentrated and the residue was dissolved in ethyl acetate. The organic layer was washed with sat. $NH_4Cl$, brine and concentrated again. The crude product was purified by silica gel column giving compound **5** (557 mg, 0.82 mmol) in 82% yield.$[\alpha]_D^{20}$ = -6 (c 0.1, DCM). $^1$H NMR (500 MHz, Chloroform-*d*) δ 7.49 – 7.33 (m, 5H), 6.32 (d, *J* = 9.5 Hz, 1H), 5.47 (s, 1H), 5.22 (dd, *J* = 10.6, 7.9 Hz, 1H), 5.08 (dd, *J* = 10.6, 9.1 Hz, 1H), 4.92 (dd, *J* = 10.6, 3.6 Hz, 1H), 4.58 (d, *J* = 7.9 Hz, 1H, Gal-1H), 4.43 (d, *J* = 8.4 Hz, 1H, GlcN-1H), 4.33 (d, *J* = 3.6 Hz, 1H), 4.27 (dd, *J* = 12.5, 1.7 Hz, 1H), 4.15 – 4.00 (q, *J* = 9.5 Hz, 1H), 4.03 (dd, *J* = 12.5, 1.7 Hz, 1H), 3.97 – 3.82 (m, 2H), 3.84 (t, *J* = 9.4 Hz, 1H), 3.70-3.76 (m, 1H), 3.57 – 3.49 (m, 2H), 3.31-3.35 (m, 1H), 3.28 – 3.16 (m, 2H), 2.09 (s, 3H), 2.04 (s, 3H), 2.03 (s, 3H), 1.97 (s, 3H), 1.72 (t, *J* = 6.3 Hz, 2H). $^{13}$C NMR (126 MHz, Chloroform-*d*) δ 171.93, 170.61, 170.33, 169.23, 137.54, 128.27, 126.40, 101.60, 101.35, 100.89, 79.19, 75.27, 75.19, 73.19, 73.11, 71.86, 69.18, 68.53, 66.16, 60.65, 53.35, 47.87, 28.84, 23.30, 20.85, 20.81, 20.66. ESI-MS: [M+H$^+$] $C_{30}H_{41}N_4O_{14}$ calculated: 681.2613, observed: 681.2645.

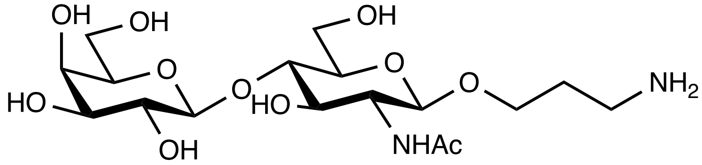


**6**

3-Aminopropyl β-D-galactopyranosyl-(1→4)-2-*N*-acetyl-2-deoxy-β-D-glucopyranoside **6**.

Compound **5** (6.8 mg, 0.01 mmol) was dissolved in 7N ammonia solution in methanol (20 mL) and stirred overnight. Upon completion, the mixture was concentrated, and the residue was redissolved in DCM (0.5 mL) and MeOH (0.5 mL) along with palladium on carbon (20 mg). The mixture was stirred overnight under $H_2$ at room temperature and then filtered *via* a PTFE membrane (pore size 0.22 μm). The filtrate was concentrated and purified by flash column chromatography, giving compound 13 (4.2 mg, 0.1 mmol) in 55% yield. $[\alpha]_D^{20}$ = -15 (c 0.1, $H_2O$). $^1$H NMR (500 MHz, Deuterium Oxide) δ 4.34 (d, *J* = 8.0 Hz, 1H, Gal-1H), 4.29 (d, *J* = 7.8 Hz, 1H, GlcN-1H), 3.87 – 3.80 (m, 2H), 3.75 (d, *J* = 3.4 Hz, 1H), 3.66 (dd, *J* = 12.3, 5.2 Hz, 1H), 3.61 – 3.46 (m, 8H), 3.45 – 3.40 (m, 1H), 3.36 (dd, *J* = 10.0, 7.8 Hz, 1H), 2.90 (t, *J* = 7.1 Hz, 2H), 1.87 (s, 3H), 1.82– 1.73 (m, 2H). $^{13}$C NMR (126 MHz, Deuterium Oxide) δ 174.60, 102.77, 101.07, 78.31, 75.26,

74.60, 72.37, 72.09, 70.84, 68.41, 67.82, 60.92, 54.92, 48.73, 37.49, 26.53, 22.00. [M-H$^+$] $C_{17}H_{31}N_2O_{11}$ calculated: 439.1933, observed: 439.1935.

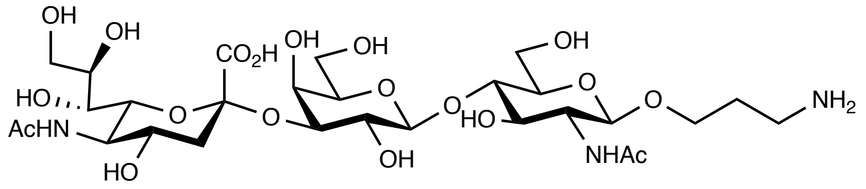

**7**

3'-Aminopropyl 5-acetamido-3,5-dideoxy-D-glycero-α-D-galacto-2- nonulopyranosylonate-(2-3)-(6-*O*-sulfo)-β-D-galactopyranosyl-(1→4)-(6-*O*-sulfo)-2-acetamido-2-deoxy-β-D-glucopyranoside **7**.

Compound **6** (2 mg, 0.004) and CMP-Neu5Ac (3.5 mg, 0.005 mmol) were dissolved in Tris-HCl buffer containing 20 mM $MgCl_2$ (450 µL, pH 8.5). Recombinant shrimp alkaline phosphatase (rSAP) (NEB) (3 µL) and PmST1 (0.4 mg/ml) were added and the reaction was placed in a shaking incubator (37 °C, 3 h) and was monitored by TLC in iPrOH:$NH_4OH$:$H_2O$ (5:2:1 by volume). The reaction was stopped by dilution with a 4-fold of 100% ethanol and put at −20 °C for 1 h to precipitate the enzyme. Precipitated protein was centrifuged (3700 rcf, 15 min, 4 °C), the supernatant carefully decanted into a round bottom flask, and evaporated. The residue was resuspended in water and purified on a P2 column equilibrated in 20% $NH_4OH$ giving compound **7** in 66% yield**.** $^1$H-NMR (700 MHz, Deuterium Oxide) $\delta = 4.54$ (d, $J = 7.7$ Hz, 1H), 4.50 (d, $J = 8.4$, 1H), 4.10 (dd, $J = 9.8$, 17.6 Hz, 1H), 3.97-4.10 (m, 4H), 3.82–3.90 (m, 4H), 3.61–3.66 (m, 6H), 3.54–3.60 (m, 5H), 3.12 (t, $J = 7.0$ Hz, 2H), 2.75 (dd, $J = 7.7$, 12.6 Hz, 1H), 2.04 (s, 3H), 2.02 (s, 3H), 1.94-1.98 (m, 2H), 1.8 (t, $J = 11.9$ Hz, 1H).

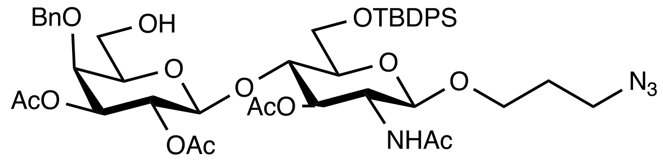

**8**

3-Azidopropyl 2,3-di-*O*-acetyl-4-*O*-benzyl-β-D-galactopyranosyl-(1→4)-6-*O*-$^t$butyldiphenylsilyl-2-*N*-acetyl-3-*O*-acetyl-2-deoxy-β-D-glucopyranoside **8**.

Molecular sieves 4Å (1.5 g) were placed in a dried 50 mL flask. Compound **4** (390 mg, 0.42 mmol) and DCM (5 mL) were added to the flask. After stirring for 1 h at room temperature, the mixture was cooled to -78°C, and then to the stirred solution, $Et_3SiH$ (220 µl, 1.38 mmol) and TfOH (94 µl, 1.22 mmol) were added successively. After being stirred for 1 h, $Et_3N$ (1 mL) and MeOH (1 mL) were added to the mixture, and the mixture was diluted with DCM and washed with aqueous $NaHCO_3$, dried over $MgSO_4$, filtered and concentrated. The crude product was purified by silica gel column giving compound **5** (238 mg, 0.61 mmol) in 61% yield. $[\alpha]_D^{20} = -8$ (c 0.1, DCM). $^1$H NMR (500 MHz, Chloroform-*d*) δ 7.74-7.67 (m, 4H), 7.48 – 7.28 (m, 11H), 5.81 (d, $J$ = 9.5 Hz, 1H), 5.28 – 5.19 (m, 1H), 5.03 – 4.94 (m, 2H), 4.90 (dd, $J$ = 10.5, 3.2 Hz, 1H), 4.74 (d, $J$ = 11.6 Hz, 1H), 4.69 (d, $J$ = 7.95 Hz, 1H, Gal-1H), 4.49 (d, $J$ = 11.6 Hz, 1H), 4.35 (d, $J$ = 7.3 Hz, 1H, GlcN-1H), 4.16 – 4.09 (m, 2H), 3.96 – 3.85 (m, 3H), 3.79 (dd, $J$ = 10.5, 7.0 Hz, 1H), 3.56 – 3.27 (m, 6H), 2.04 (s, 3H), 2.00 (s, 3H), 1.97 (s, 3H), 1.92- 1.85 (m, 2H) 1.83 (s, 3H), 1.06 (s, 9H). $^{13}$C NMR (126 MHz, Chloroform-*d*) δ 171.29, 170.34, 170.26, 169.36, 135.84, 135.45, 129.94, 129.59, 128.56, 128.26, 128.23, 128.17, 127.92, 127.71, 115.34, 101.20, 100.28, 79.18, 75.60, 74.98, 74.87, 73.84, 73.64, 73.19,

72.48, 70.09, 65.40, 61.74, 52.69, 48.14, 28.92, 26.81, 23.30, 20.85, 20.83, 20.63, 19.33. ESI-MS: [M+H+] $C_{46}H_{61}N_4O_{14}Si$ calculated: 921.3947, observed: 921.3926.

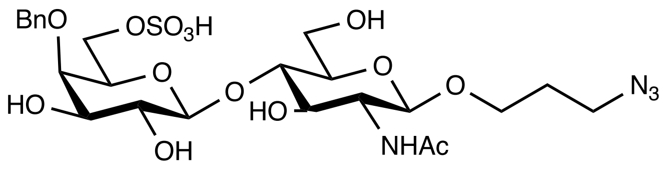


**9**

3-Azidopropyl 4-*O*-benzyl-6-*O*-sulfo-β-D-galactopyranosyl-(1→4)-2-*N*-acetyl-2-deoxy-β-D glucopyranoside **9**.

Compound **8** (92 mg, 0.1 mmol) was dissolved in pyridine (5 mL) along with $SO_3$·Pyr (31.8 mg, 0.2 mmol). The mixture was stirred for 3 h and monitored by TLC. Upon completion, the solution was nitrogen dried and the residue was redissolved in THF (5 mL) with 1M tetra-*n*-butylammonium fluoride (TBAF) in THF solution (2 mL, 2 mmol). When all starting material was consumed, the reaction was dried under vacuum and then redissolved in 7N ammonia in methanol (10 mL). The reaction was monitored by mass spec. Upon completion, the mixture was first dried and then redissolved in water (2 mL), loaded to a Sep-Pak® C18 Cartridge, then washed with an excess of water to elute TBAF. Finally, the crude was eluted out with pure acetonitrile, and the mixture was concentrated *in vacuo*. The residue was purified by flash column chromatography giving compound **9** (38 mg, 0.06 mmol) in 60% yield. $[\alpha]_D^{20}$ = -6 (c 0.1, $H_2O$). $^1$H NMR (500 MHz, Methanol-$d_4$) δ 7.47 –7.22 (m, 5H), 5.12 (br, 1H), 4.94 (d, *J* = 10.7 Hz, 1H), 4.67 (d, *J* = 10.7 Hz, 1H), 4.43 (d, *J* = 7.65 Hz, 1H, Gal-1H), 4.39 (d, *J* = 8.3 Hz, 1H, GlcN-1H) , 4.16 – 4.08 (m, 2H), 3.96 – 3.80 (m, 5H), 3.74 – 3.50 (m, 6H), 3.39 – 3.33 (m, 3H), 1.97 (s, 3H), 1.86 – 1.74 (m, 2H). $^{13}$C NMR (126 MHz, Methanol-$d_4$) δ 138.49, 128.20, 127.85, 127.31, 103.44, 101.29, 79.57, 76.49, 75.29, 75.07, 74.06, 72.83, 72.70, 71.28, 65.79, 65.58, 60.58, 58.06, 55.25, 28.65, 21.57. ESI-MS: [M+2Na-H+] $C_{24}H_{35}N_4O_{14}SNa_2$ calculated: 686.1669, observed: 686.1660.

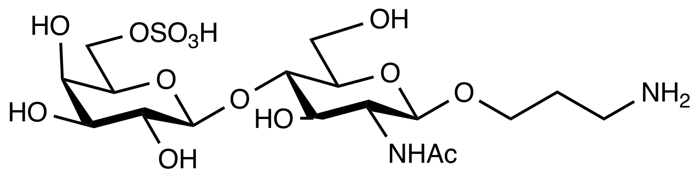


**10**

3-Aminopropyl 6-*O*-sulfo-β-D-galactopyranosyl-(1→4)-2-*N*-acetyl-2-deoxy-β-D-glucopyranoside **10**.

Compound **9** (64.1 mg, 0.1 mmol) was dissolved in DCM (5 mL) and MeOH (5 mL) with palladium on carbon (200 mg). The mixture was stirred overnight under $H_2$ at room temperature and then filtered *via* a PTFE membrane (pore size 0.22 μm). The filtrate was concentrated with no further purification, giving compound **7** (51 mg, 0.1 mmol) in 98% yield. $[\alpha]_D^{20}$ = -18 (c 0.1, $H_2O$). $^1$H NMR (500 MHz, Deuterium Oxide) δ 4.36 (d, *J* = 8.0 Hz, 1H, Gal-1H), 4.35 (d, *J* = 8.0 Hz, 1H, GlcN-1H), 4.05 – 3.99 (m, 2H), 3.89-3.78 (m, 4H), 3.66 (m, 2H), 3.61 – 3.43 (m, 6H), 3.35 (t, *J* = 8.0 Hz, 1H), 2.91 (t, *J* = 7.0 Hz, 2H), 1.87 (s, 3H), 1.81 – 1.74 (m, 2H). $^{13}$C NMR (126 MHz, Deuterium Oxide) δ 174.59, 102.53, 101.02, 78.91, 74.53, 72.71, 72.17, 71.79, 71.74, 70.63, 68.11, 67.82, 67.18, 54.98, 37.52, 26.49, 22.00. ESI-MS: [M-H+] $C_{17}H_{31}N_2O_{14}S$ calculated: 519.1501, observed: 519.1524.

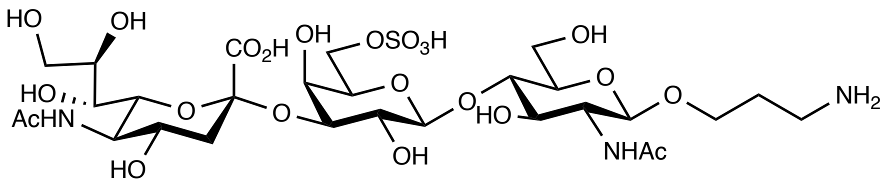

**11**

3'-Aminopropyl 5-acetamido-3,5-dideoxy-D-glycero-α-D-galacto-2- nonulopyranosylonate-(2-3)-(6-*O*-sulfo)-β-D-galactopyranosyl-(1→4)-(6-*O*-sulfo)-2-acetamido-2-deoxy-β-D-glucopyranoside **11**.

The same reaction conditions were used as mentioned above for the synthesis of compound **7**. The residue was purified using HILIC HPLC column, starting with 100% buffer A (25 mM NH4OAc, 20% water, 80% acetonitrile) and then gradually switched to 100% buffer B (25 mM NH4OAc, 90% water, 10% acetonitrile) over 1 hour. The product (NAc-amide) was detected at 214 nm giving compound **11** in 35% yield. $^{1}$H-NMR (700 MHz, Deuterium Oxide) $\delta$ = 4.59 (d, $J$ = 8.4 Hz, 1H), 4.45 (d, $J$ = 7.7, 1H), 4.18 (m, 2H), 4.12 (dd, $J$ = 9.1, 13.3 Hz, 2H), 3.94-4.02 (m, 4H), 3.82-3.89 (m, 4H), 3.74-3.76 (m, 2H), 3.66–3.73 (m, 4H), 3.62-3.66 (m, 4H), 3.45-3.60 (m, 3H), 3.07 (t, $J$ = 7 Hz, 2H), 2.75 (dd, $J$ = 4.9, 12.6 Hz, 1H), 2.03 (s, 3H), 2.01 (s, 3H), 1.91-1.96 (m, 2H), 1.80 (t, $J$ = 11.9 Hz, 1H).

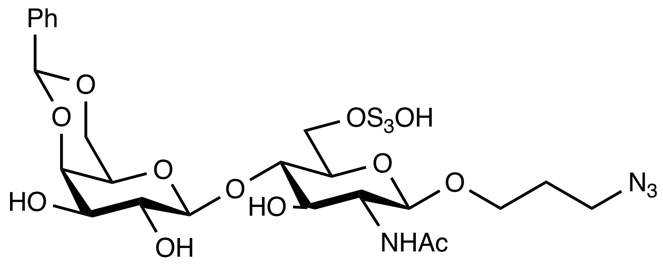

**12**

3-Azidopropyl 4,6-*O*-benzylidene-β-D-galactopyranosyl-(1→4)-2-*N*-acetyl-2-deoxy-6-*O*-sulfo-β-D-glucopyranoside **12**.

Compound **5** (68 mg, 0.1 mmol) was dissolved in pyridine (5 mL) along with $SO_3$·Pyr (31.8 mg, 0.2 mmol). The mixture was stirred for 3 h and the progress of the reaction was monitored by TLC. Upon completion, solution was nitrogen dried and the residue was redissolved in 7N ammonia in methanol (10 mL) then stirred overnight. The reaction was monitored by mass spec. Upon reaction completion, the reaction mixture was concentrated *in vacuo*. The residue was purified by flash column chromatography giving compound **12** (46 mg, 0.073 mmol) in 73% yield. $[\alpha]_D^{20}$ = -77 (c 0.1, MeOH). $^{1}$H NMR (500 MHz, Methanol-$d_4$) δ 7.61 – 7.24 (m, 5H), 5.64 (d, $J$ = 1.8 Hz, 1H), 4.65 (dd, $J$ = 7.8 Hz, 1H, Gal-1H), 4.42 (m, 2H, GlcN-1H), 4.32 (d, $J$ = 10.9 Hz, 1H), 4.28 – 4.15 (m, 3H), 3.95 – 3.89 (m, 1H), 3.78 – 3.54 (m, 8H), 3.39 – 3.32 (m, 3H), 1.98 (s, 3H), 1.89 – 1.74 (m, 2H). $^{13}$C NMR (126 MHz, Methanol-$d_4$) δ 138.08, 128.49, 127.64, 126.02, 102.92, 101.45, 100.73, 78.64, 76.04, 73.02, 72.52, 71.99, 70.86, 68.77, 66.88, 66.07, 65.68, 55.13, 48.45, 28.62, 21.61. ESI-MS: [M-H$^{+}$] $C_{24}H_{33}N_4O_{14}S$ calculated: 633.1719, observed: 633.1721.

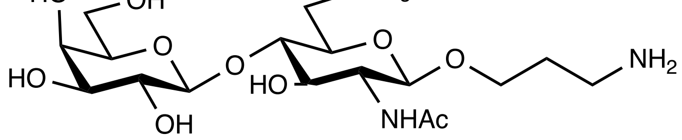

**13**

3-Aminopropyl β-D-galactopyranosyl-(1→4)-2-*N*-acetyl-2-deoxy-6-*O*-sulfo-β-D-glucopyranoside

**13**.

Compound **12** (63.4 mg, 0.1 mmol) was dissolved in DCM (5 mL) and MeOH (5 mL) with palladium on carbon (200 mg). The mixture was stirred overnight under $H_2$ at room temperature and then filtered *via* a PTFE membrane (pore size 0.22 μm). The filtrate was concentrated with no further purification, giving compound **7** (52 mg, 0.1 mmol) in 100% yield. $[\alpha]_D^{20}$ = -13 (c 0.1, $H_2O$). $^1$H NMR (500 MHz, Deuterium Oxide) δ 4.36 (d, *J* = 8.3 Hz, 1H, Gal-1H), 4.34 (d, *J* = 7.9 Hz, 1H, GlcN-1H), 4.23 (d, *J* = 11.2 Hz, 1H), 4.14 (dd, *J* = 11.2, 3.8 Hz, 1H), 3.84 – 3.78 (m, 1H), 3.73 (d, *J* = 3.3 Hz, 1H), 3.65 – 3.44 (m, 9H), 3.34 (t, *J* = 8.1 Hz, 1H), 2.91 (t, *J* = 6.8 Hz, 2H), 1.85 (s, 3H), 1.82 – 1.72 (m, 2H). $^{13}$C NMR (126 MHz, Deuterium Oxide) δ 174.60, 102.43, 101.18, 77.24, 75.24, 72.35, 71.99, 70.84, 68.46, 68.14, 66.13, 60.94, 54.88, 48.71, 37.61, 26.53, 21.99. ESI-MS: [M-H$^+$] $C_{17}H_{31}N_2O_{14}S$ calculated: 519.1501, observed: 519.1575.

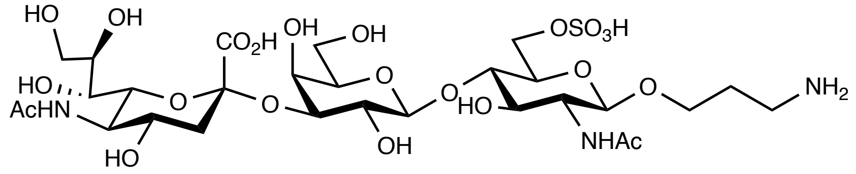


**14**

3'-Aminopropyl 5-acetamido-3,5-dideoxy-D-glycero-α-D-galacto-2-nonulopyranosylonate-(2-3)-β-D-galactopyranosyl-(1→4)-(6-*O*-sulfo)-2-acetamido-2-deoxy-β-D-glucopyranoside **14**.

The same reaction conditions were used as mentioned above for synthesis of compound **7**. The residue was purified using HILIC HPLC column, starting with 100% buffer A (25 mM NH4OAc, 20% water, 80% acetonitrile) and then gradually switched to 100% buffer B (25 mM NH4OAc, 90% water, 10% acetonitrile) over 1 hour. The product (NAc-amide) was detected at 214 nm giving compound **14** in 64% yield .$^1$H-NMR (700 MHz, Deuterium Oxide) *δ* = 4.59 (d, *J* = 7.7 Hz, 1H), 4.53 (d, *J* = 8.4, 1H), 4.41 (d, *J* = 10.9, 1H), 4.30 (dd, *J* = 11.2, 4.2 Hz, 1H), 4.09 (dd, *J* = 9.8, 7, 1H), 3.92-3.98 (m, 3H), 3.81–3.89 (m, 4H), 3.62–3.69 (m, 4H), 3.52–3.62 (m, 4H), 3.12 (t, *J* = 7.0 Hz, 2H), 2.74 (dd, *J* = 4.2, 12.6 Hz, 1H), 2.03 (s, 3H), 2.01 (s, 3H), 1.89-1.94 (m, 2H), 1.78 (t, *J* = 11.9 Hz, 1H).

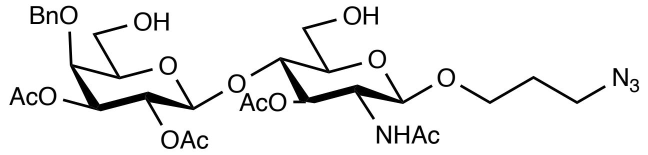


**15**

3-Azidopropyl 2,3-di-*O*-acetyl-4-*O*-benzyl-β-D-galactopyranosyl-(1→4)-2-*N*-acetyl-2-deoxy-3-*O*-acetyl-β-D-glucopyranoside **15**.

Compound **8** (920 mg, 1 mmol) was dissolved in THF (10 mL). A solution composed of 1 M TBAF in THF (1.1 mL, 1.1 mmol) and acetic acid (66 μL, 1.1 mmol) was added dropwise into compound **8** and the mixture was stirred overnight. Upon completion, the solution was first concentrated and the residue was dissolved in ethyl acetate. The organic layer was washed with sat. $NH_4Cl$, brine and concentrated again. The crude product was purified by silica gel column giving compound **15** (579 mg, 0.85 mmol) in 85% yield. $[\alpha]_D^{20}$ = -11 (c 0.1, DCM). $^1$H NMR (500 MHz, Chloroform-*d*) δ 7.35 – 7.20 (m, 5H), 6.48 (d, *J* = 9.5 Hz, 1H), 5.15 (dd, *J* = 10.4, 7.9 Hz, 1H), 4.99 (t, *J* = 6.5 Hz, 1H), 4.90 (dd, *J* = 10.4, 3.1 Hz, 1H), 4.64 (d, *J* = 11.7 Hz, 1H), 4.54 (d, *J* = 7.9 Hz, 1H, Gal-1H), 4.47 (d, *J* = 11.7 Hz, 1H), 4.39 (d, *J* = 8.2 Hz, 1H, GlcN-1H), 3.97 (q, 9.6 Hz, 1H), 3.92-3.80 (m, 4H), 3.73 (dd, *J* = 10.9, 6.6 Hz, 1H), 3.65 (dd, *J* = 12.2, 3.7 Hz, 1H), 3.59 – 3.47 (m, 3H), 3.36 – 3.21 (m, 2H), 2.98 – 2.88 (m, 1H), 2.00 (s, 3H), 1.99 (s, 3H), 1.93 (s, 3H), 1.86 (s, 3H), 1.75 –

1.67 (m, 2H). $^{13}C$ NMR (126 MHz, Chloroform-*d*) δ 171.66, 170.61, 170.40, 169.64, 137.66, 128.42, 128.08, 127.97, 101.16, 100.89, 75.21, 74.84, 74.70, 73.89, 73.80, 73.76, 70.32, 66.10, 61.23, 60.61, 53.54, 52.10, 47.95, 28.78, 23.12, 21.05, 20.78, 20.74. ESI-MS: [M+H$^+$] $C_{30}H_{43}N_4O_{14}$ calculated: 683.2770, observed: 683.2757.

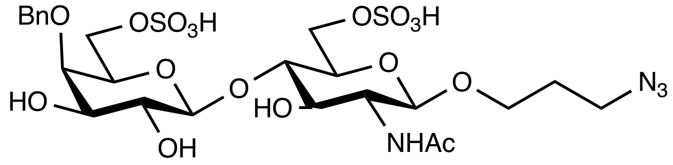


**16**

3-Azidopropyl 4-*O*-benzyl-6-*O*-sulfo-β-D-galactopyranosyl-(1→4)-2-*N*-acetyl-2-deoxy-6-*O*-sulfo-β-D-glucopyranoside **16**.

Compound **15** (68 mg, 0.1 mmol) was dissolved in pyridine (5 mL) along with $SO_3$·Pyr (63.6 mg, 0.4 mmol). The mixture was stirred for 3 h and the progress of the reaction was monitored by TLC. Upon completion, the solution was nitrogen dried and the residue was redissolved in 7N ammonia in methanol (10 mL) then stirred for overnight. The reaction was monitored by mass spec and then concentrated *in vacuo*. The residue was purified by flash column chromatography giving compound **16** (42.9 mg, 0.06 mmol) in 60% yield. $[\alpha]_D^{20}$ = -87 (c 0.1, MeOH). $^1H$ NMR (500 MHz, Methanol-$d_4$) δ 7.53 – 7.13 (m, 5H), 4.93 (d, *J* = 10.7 Hz, 1H), 4.65 (d, *J* = 10.7 Hz, 1H), 4.51 (d, *J* = 7.8 Hz, 1H, Gal-1H), 4.39 (d, *J* = 8.3 Hz, 1H, GlcN-1H), 4.35 – 4.28 (m, 2H), 4.16 – 4.06 (m, 2H), 3.94 – 3.84 (m, 3H), 3.75 – 3.48 (m, 7H), 3.36-3.38 (m, 2H), 1.98 (s, 3H), 1.86 – 1.68 (m, 2H). $^{13}C$ NMR (126 MHz, Methanol-$d_4$) δ 172.44, 138.43, 128.21, 127.88, 127.35, 103.45, 101.21, 79.60, 76.57, 75.23, 74.07, 72.99, 72.96, 72.92, 71.51, 65.99, 65.97, 54.98, 48.45, 28.62, 21.60. [M-H$^+$] $C_{24}H_{35}N_4O_{17}S_2$ calculated: 715.1444, observed: 715.1423.

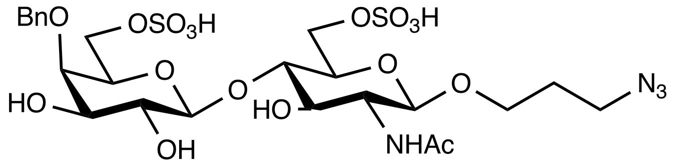


**17**

3-Aminopropyl 6-*O*-sulfo-β-D-galactopyranosyl-(1→4)-2-*N*-acetyl-2-deoxy-6-*O*-sulfo-β-D-glucopyranoside **17**.

Compound **16** (7.1 mg, 0.01 mmol) was dissolved in DCM (0.5 mL) and MeOH (0.5 mL) with palladium on carbon (20 mg). The mixture was stirred overnight under $H_2$ at room temperature and then filtered *via* a PTFE membrane (pore size 0.22 μm). The filtrate was concentrated with no further purification, giving compound **17** (6 mg, 0.1 mmol) in 98% yield. $[\alpha]_D^{20}$ = -21 (c 0.1, $H_2O$). $^1H$ NMR (500 MHz, Deuterium Oxide) δ 4.36 (d, *J* = 8.0 Hz, 1H, Gal-1H), 4.36 (d, *J* = 8.0 Hz, 1H, GlcN-1H), 4.25 (dd, *J* = 11.2, 2.0 Hz, 1H), 4.14 (dd, *J* = 11.2, 4.9 Hz, 1H), 4.06 – 3.98 (m, 2H), 3.84 – 3.76 (m, 3H), 3.68 – 3.63 (m, 1H), 3.71 – 3.48 (m, 4H), 3.51 (dd, *J* = 10.0, 3.2 Hz, 1H), 3.40 – 3.28 (m, 1H), 2.92 (t, *J* = 6.8 Hz, 2H), 1.86 (s, 3H), 1.81 – 1.73 (m, 2H). $^{13}C$ NMR (126 MHz, Deuterium Oxide) δ 174.65, 102.64, 101.19 78.46, 72.68, 72.35, 72.18, 71.82, 70.70, 68.19, 67.09, 66.41, 54.94, 48.77, 37.69, 26.55, 22.06. [M-H$^+$] $C_{17}H_{31}N_2O_{17}S_2$ calculated: 599.1069, observed: 599.1085.

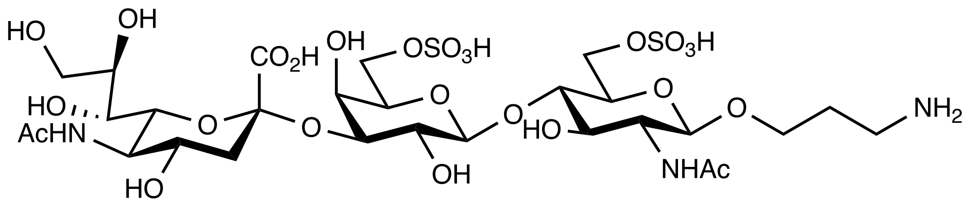

**18**

3'-Aminopropyl 5-acetamido-3,5-dideoxy-D-glycero-α-D-galacto-2- nonulopyranosylonate-(2-3)-(6-*O*-sulfo)-β-D-galactopyranosyl-(1→4)-(6-*O*-sulfo)-2-acetamido-2-deoxy-β-D-glucopyranoside **18**.

The same reaction conditions were used as mentioned above for synthesis of compound **7**. The residue was purified using HILIC HPLC column, starting with 100% buffer A (25 mM $NH_4OAc$, 20% water, 80% acetonitrile) and then gradually switched to 100% buffer B (25 mM NH4OAc, 90% water, 10% acetonitrile) over 1 hour. The product (NAc-amide) was detected at 214 nm giving compound **18** in 35% yield**.** $^1$H-NMR (700 MHz, Deuterium Oxide) $\delta$ = 4.61 (d, $J$ = 7.7 Hz, 1H), 4.54 (d, $J$ = 7.7 Hz, 1H), 4.45 (d, $J$ = 10.5 Hz, 1H), 4.30 (dd, $J$ = 11.9, 13.3 Hz, 1H), 4.09-4.19 (m, 5H), 3.94-4.02 (m,4H), 3.83-3.91 (m, 5H), 3.72-3.80 (m, 6H), 3.66-3.72 (m, 2H), 3.62-3.66 (m, 3H), 3.53-3.59 (m, 3H), 3.12 (t, $J$ = 7 Hz, 2H), 2.74 (dd, $J$ = 4.2, 8.4 Hz, 1H), 2.03 (s, 3H), 2.02 (s, 3H), 1.91-1.99 (m, 2H), 1.8 (t, $J$ = 11.9 Hz, 1H).

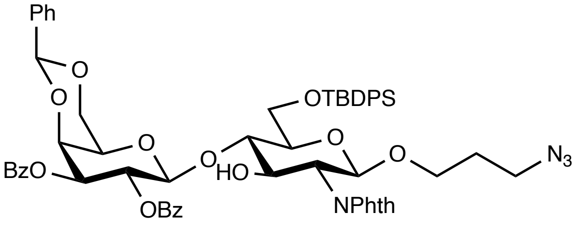


**3**

$^{1}$H-NMR ($CDCl_3$, 500 MHz) of **3**

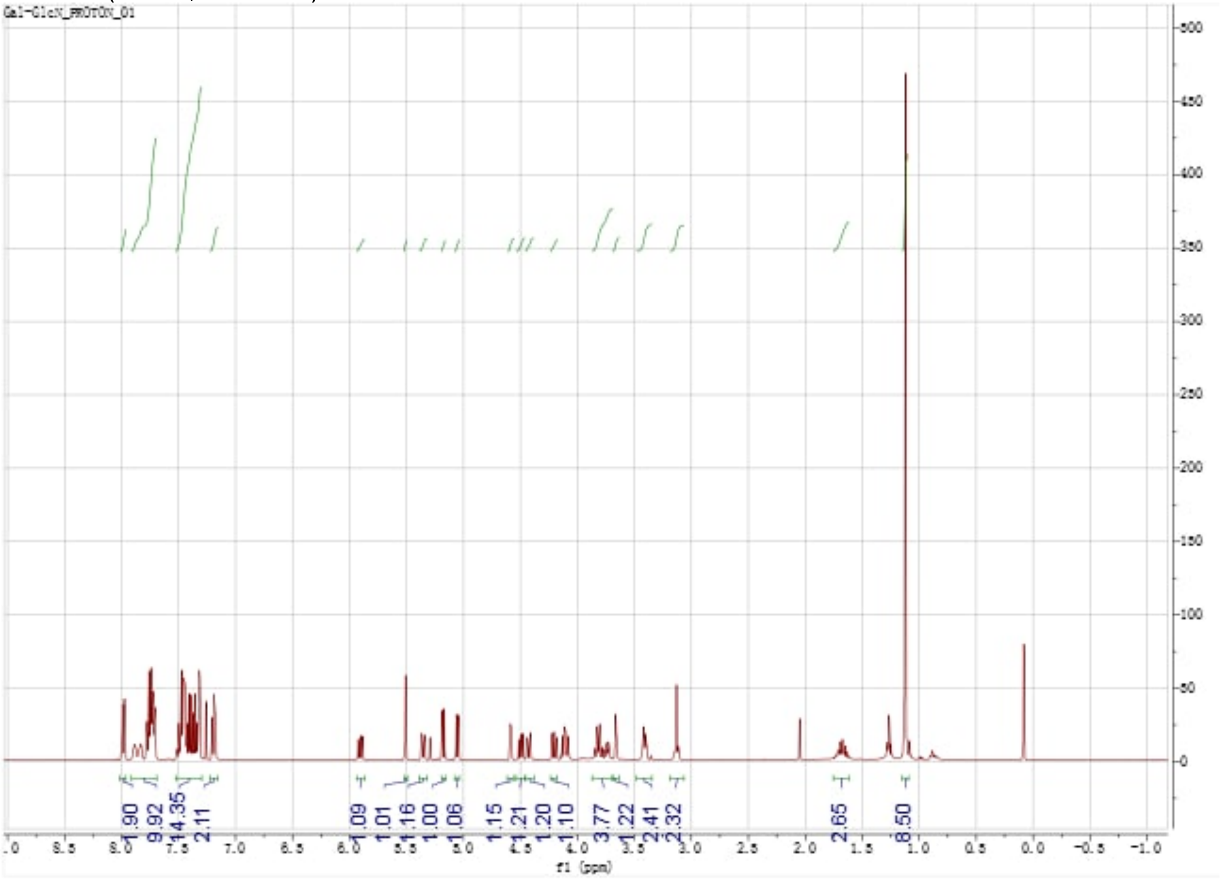

$^{13}$C-NMR ($CDCl_3$, 126 MHz) of **3**

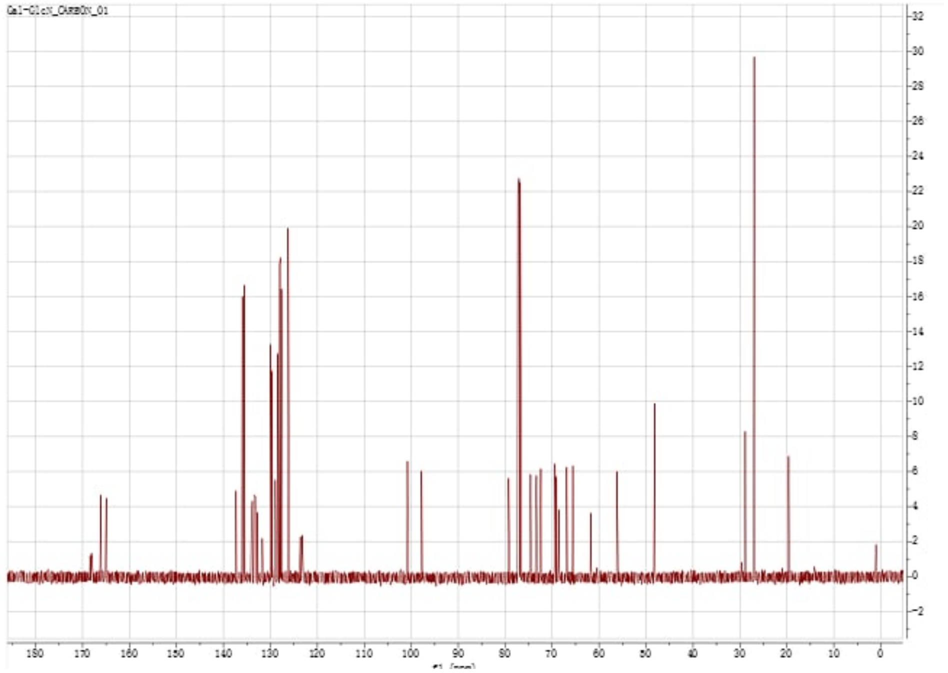


gHSQC ($CDCl_3$, 500 MHz) of **3**

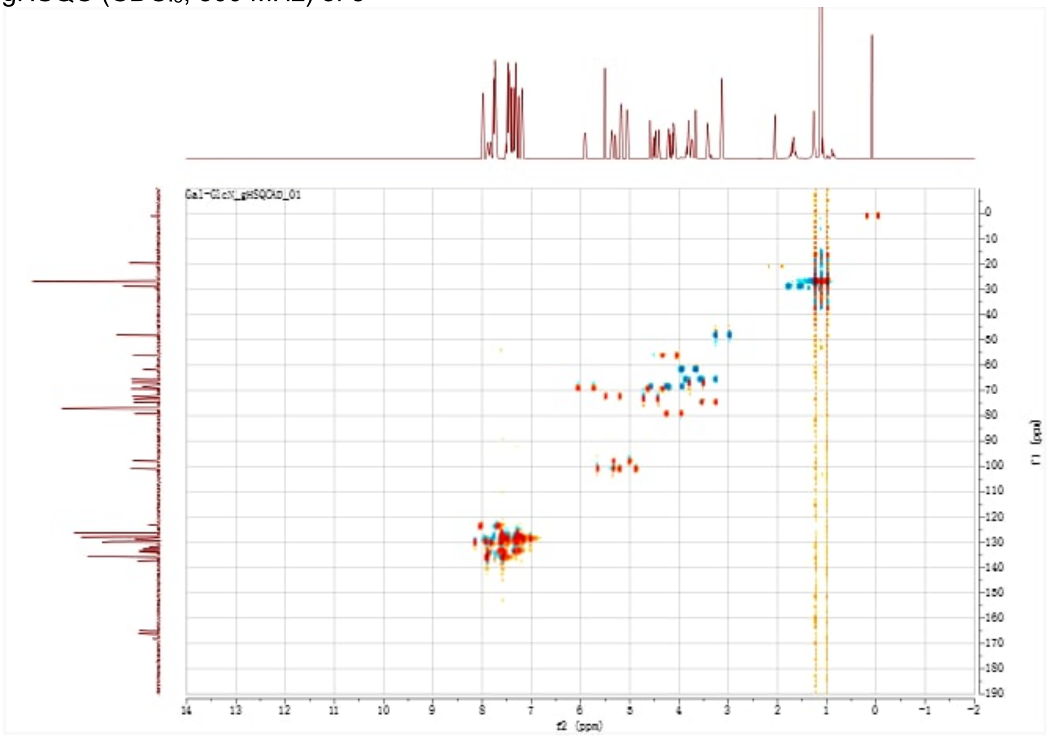

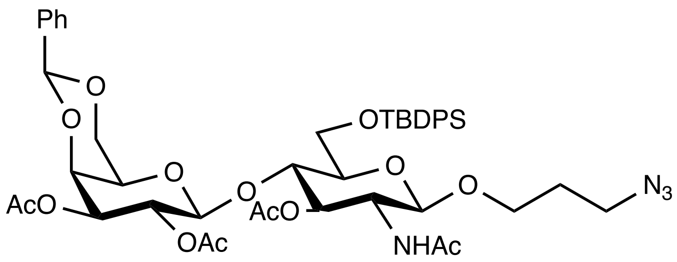

**4**

$^{1}$H-NMR ($CDCl_3$, 500 MHz) of **4**

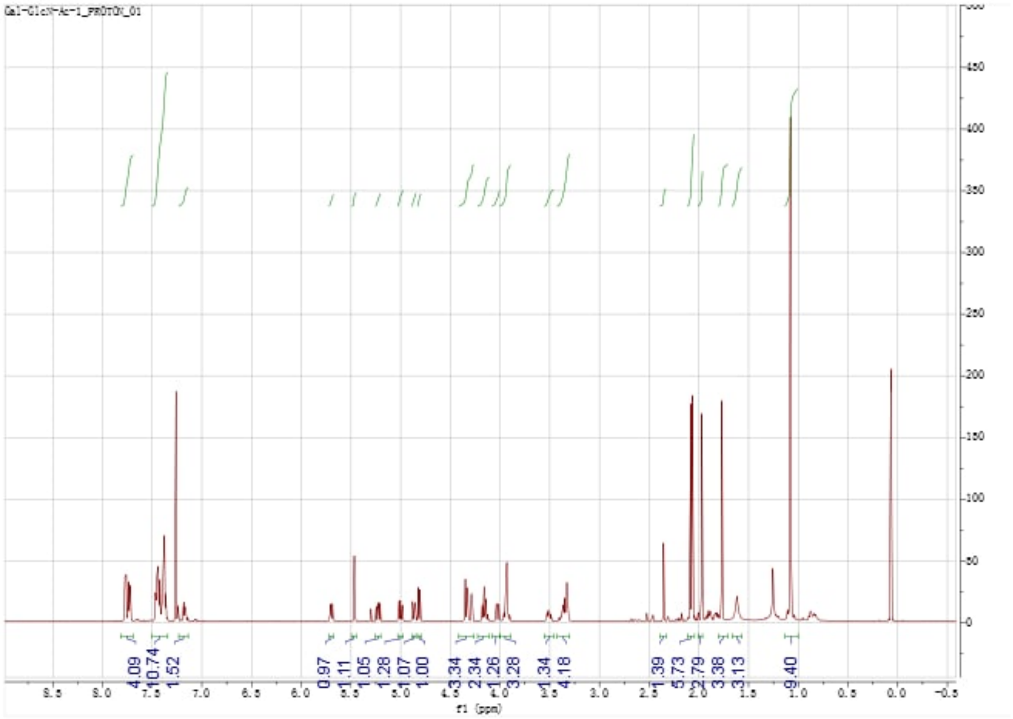

$^{13}$C-NMR ($CDCl_3$, 126 MHz) of **4**

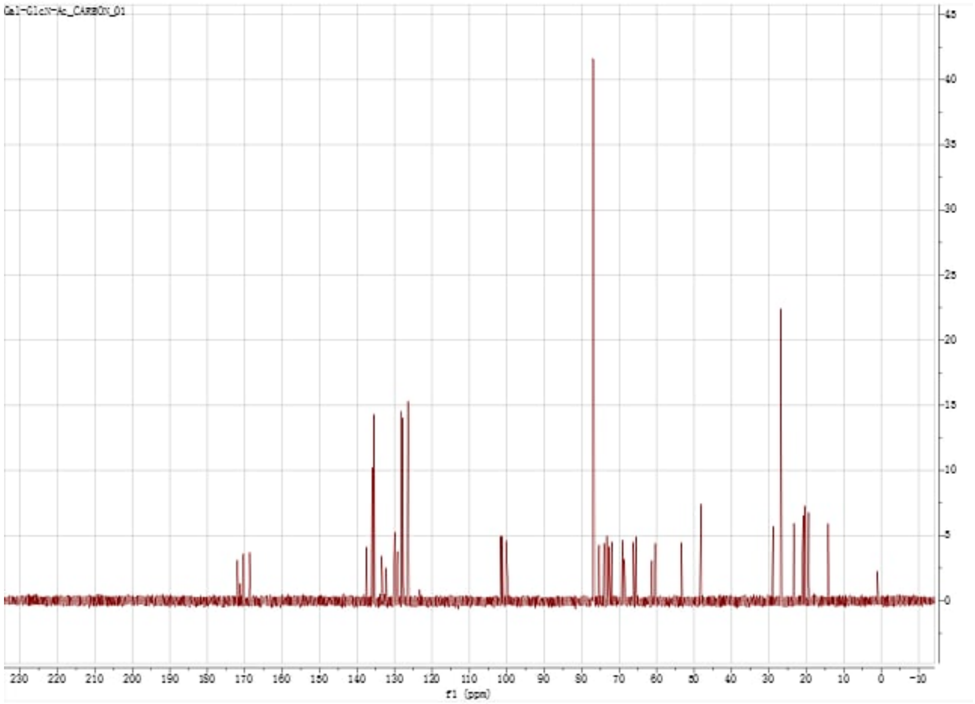


gCOSY(CDCl₃, 500 MHz) of **4**

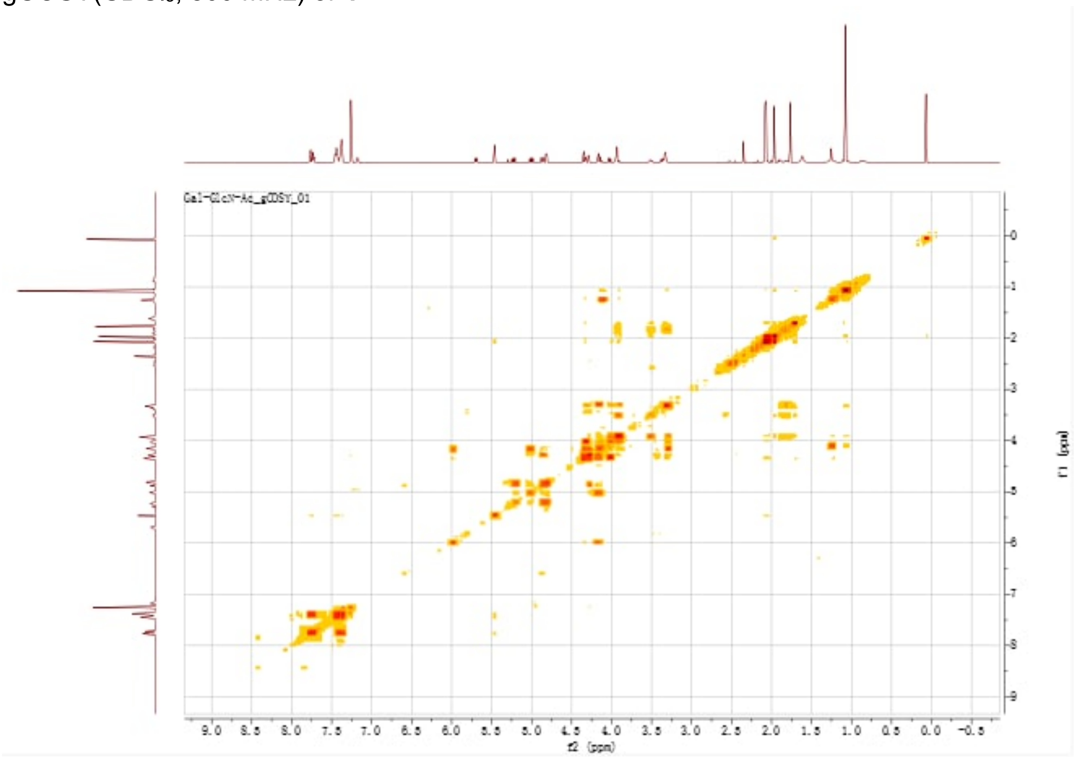


gHSQC(CDCl₃, 500 MHz) of **4**

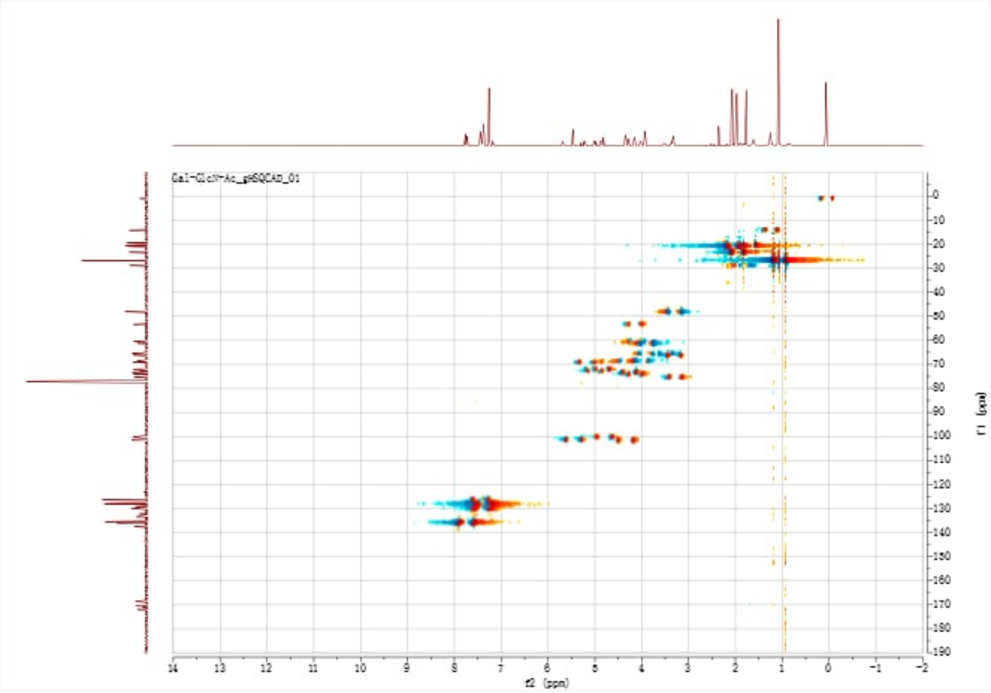


gHMBC($CDCl_3$, 500 MHz) of **4**

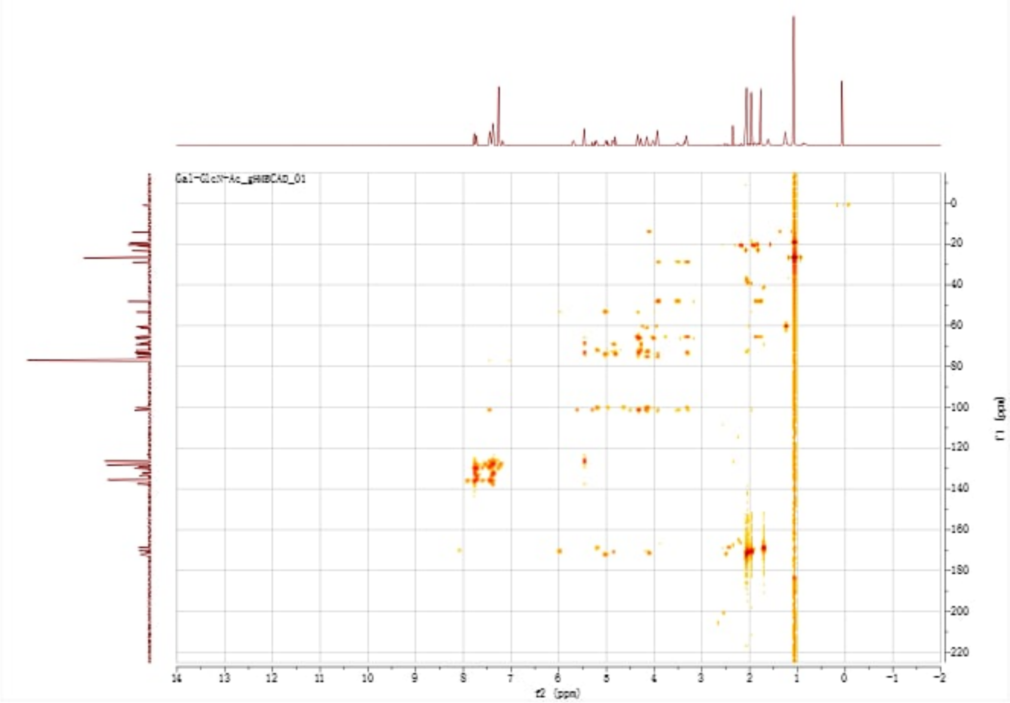

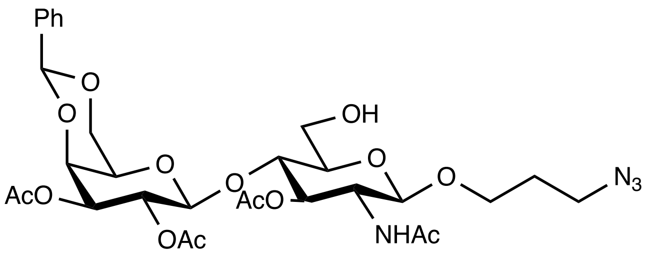


**5**

[1]H-NMR ($CDCl_3$, 500 MHz) of **5**

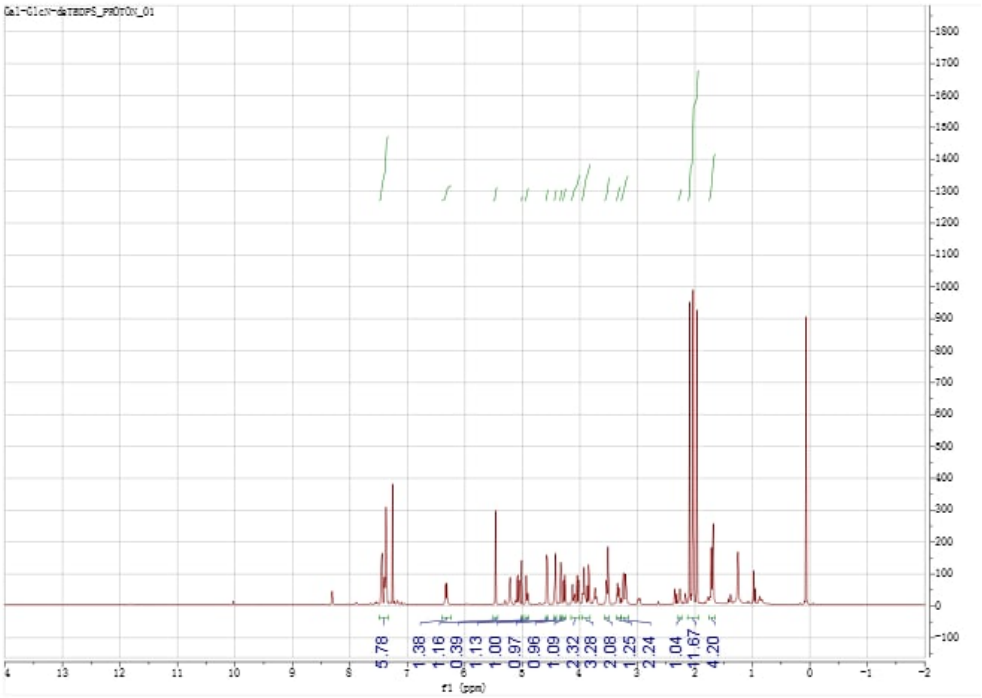


[13]C-NMR ($CDCl_3$, 126 MHz) of **5**

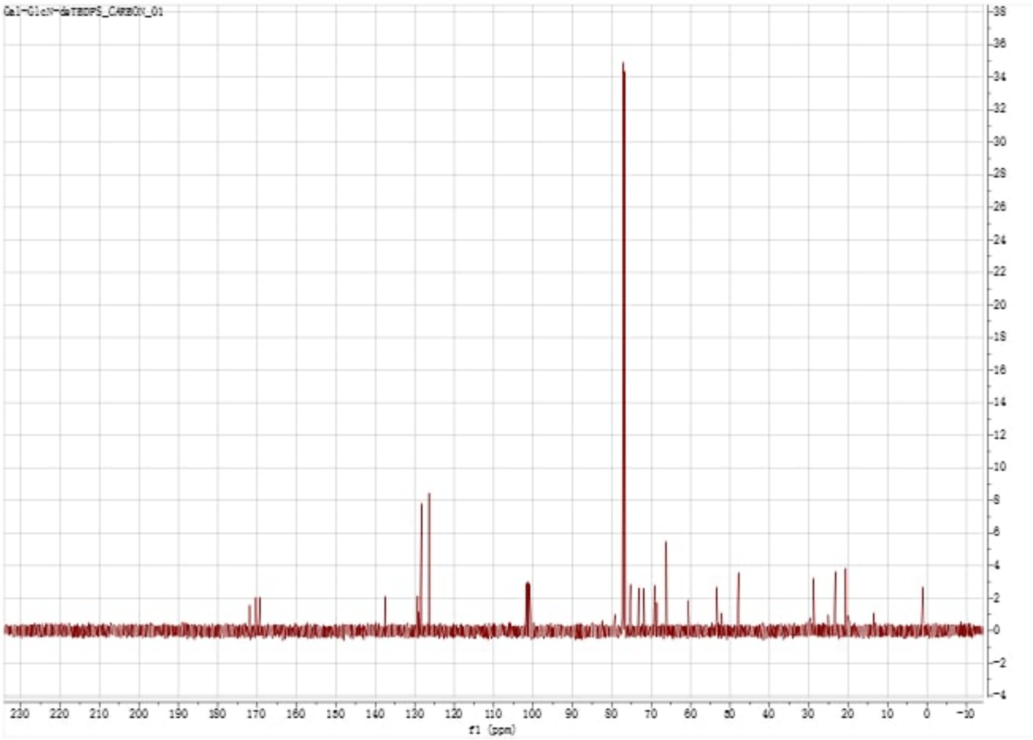


gCOSY($CDCl_3$, 500 MHz) of **5**

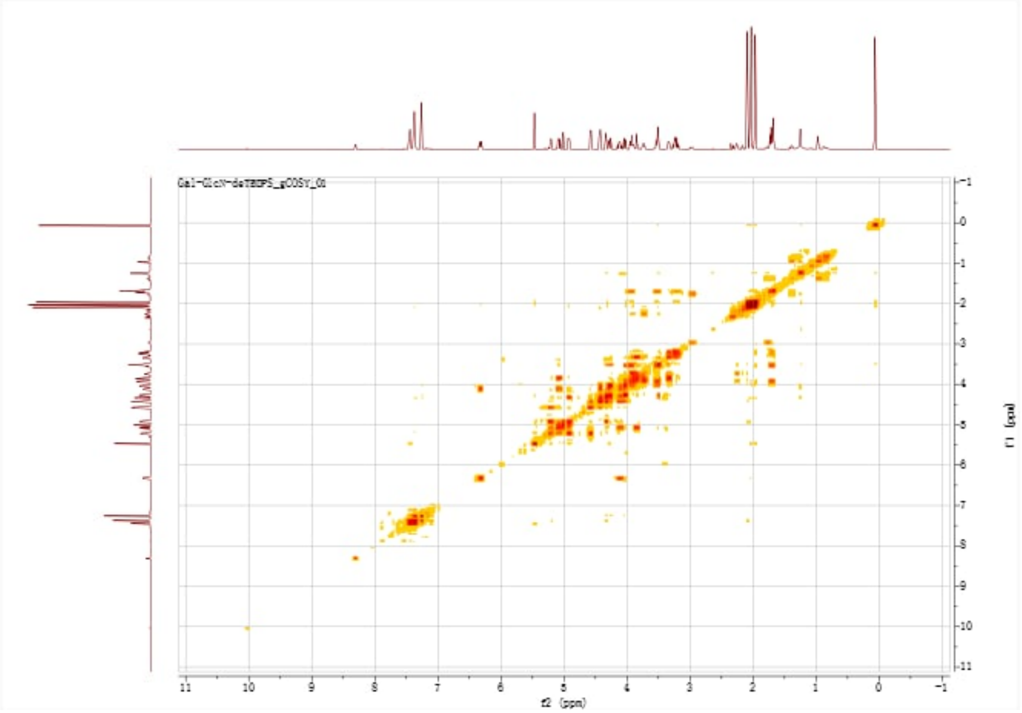


gHSQC($CDCl_3$, 500 MHz) of **5**

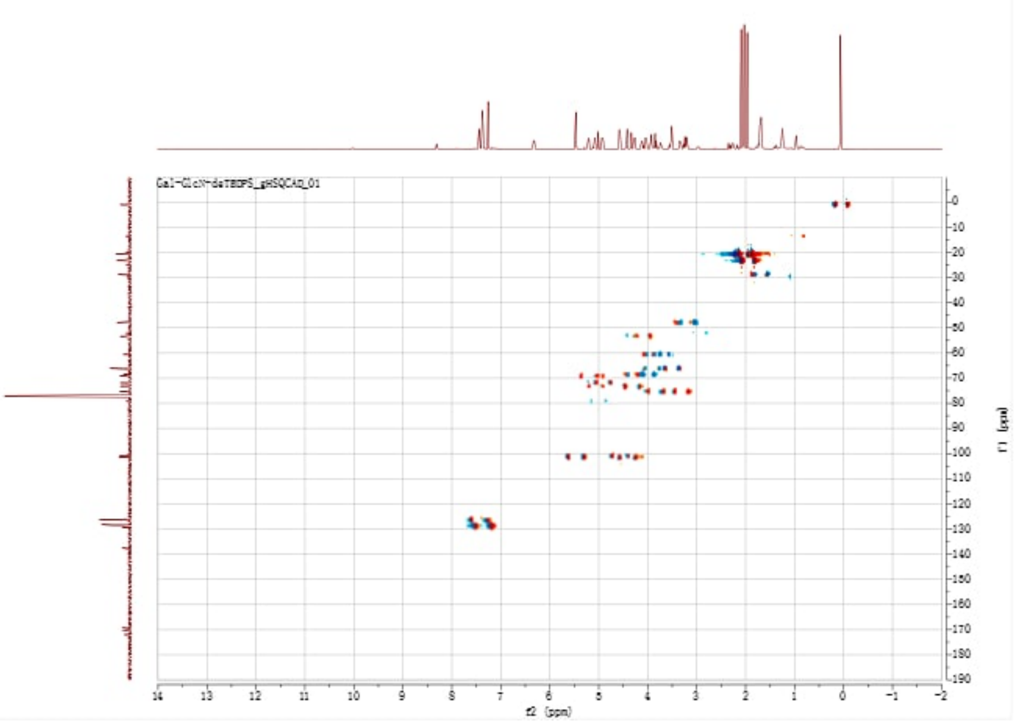


gHMBC($CDCl_3$, 500 MHz) of **5**

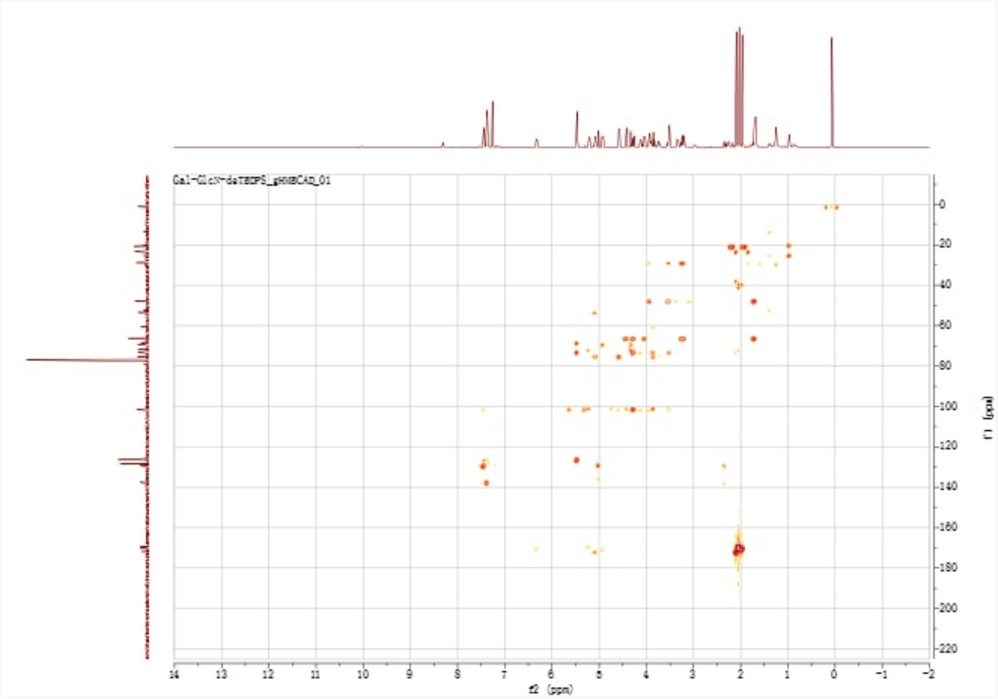

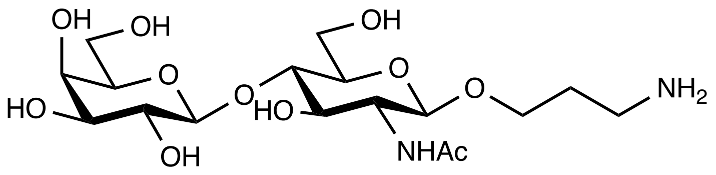


**6**

[1]H-NMR ($CDCl_3$, 500 MHz) of **6**

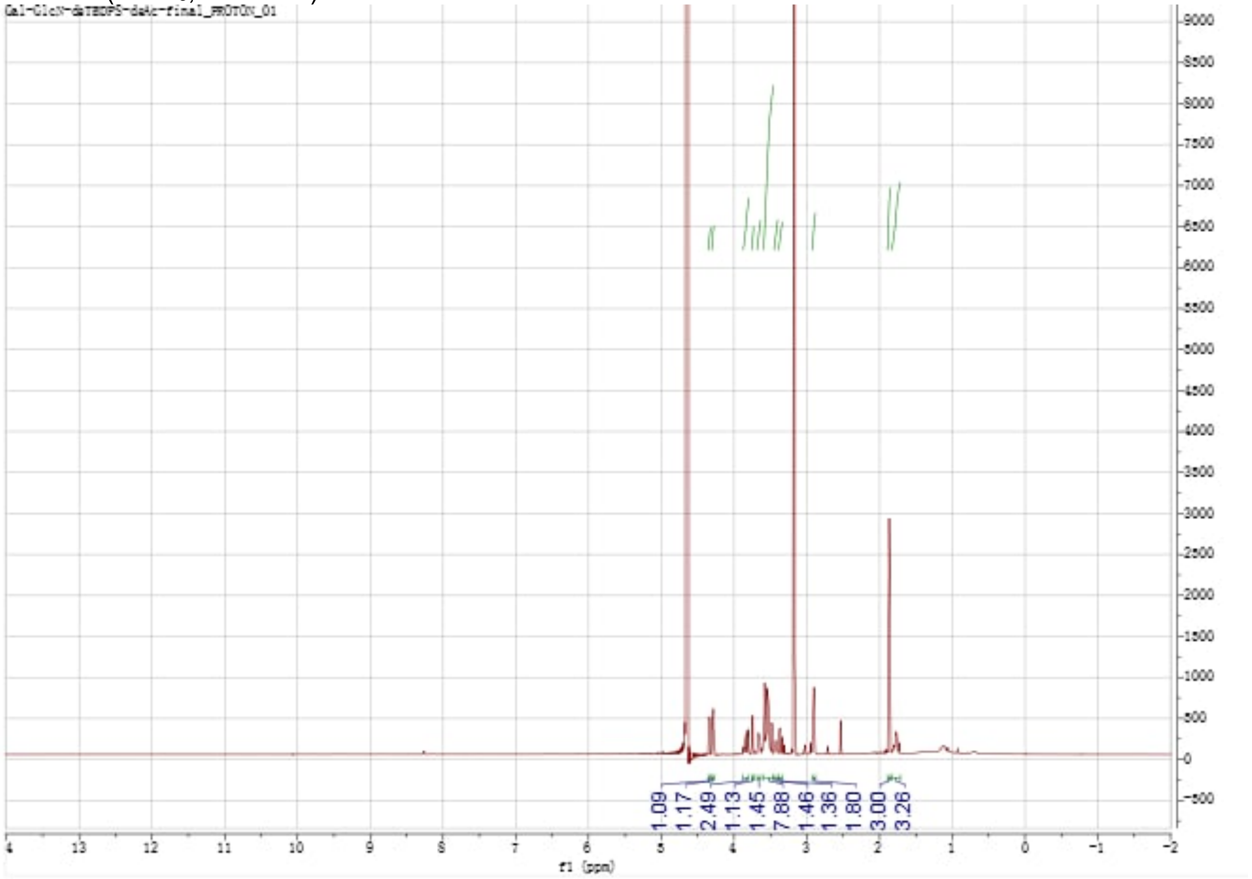

$^{13}C$-NMR ($CDCl_3$, 126 MHz) of **6**

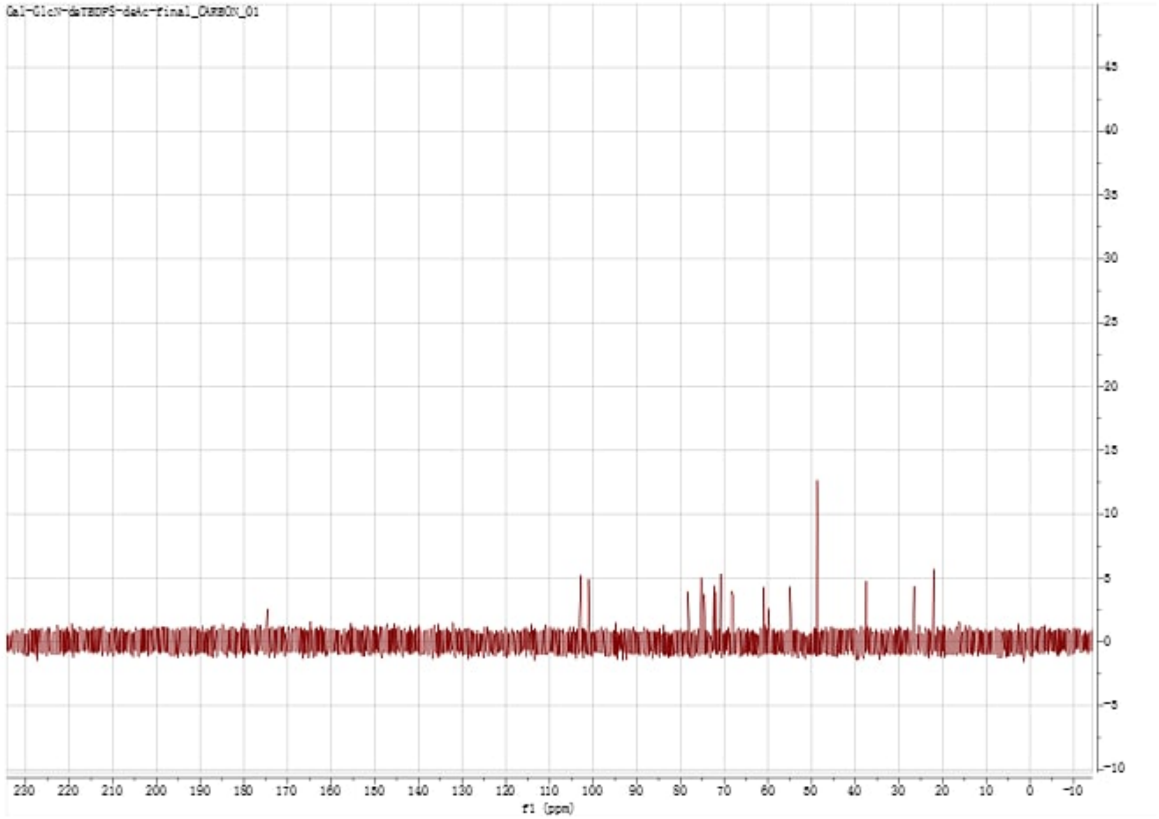


gCOSY($CDCl_3$, 500 MHz) of **6**

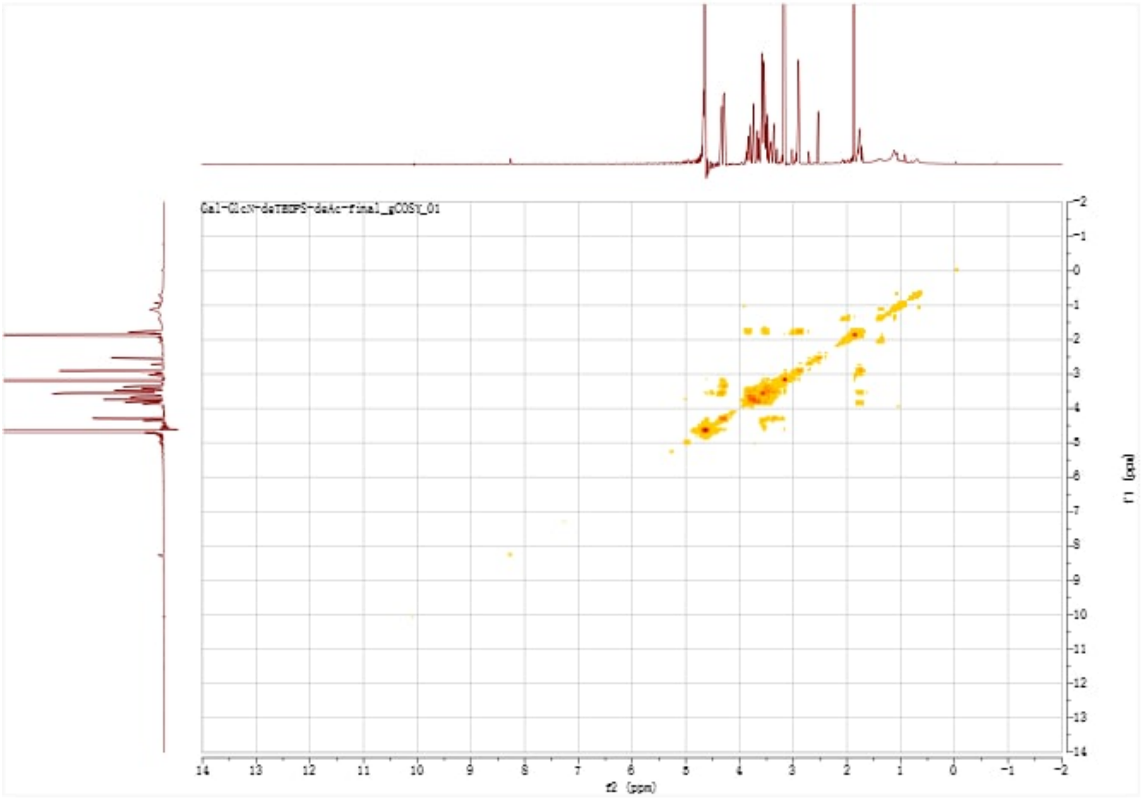

gHSQC($CDCl_3$, 500 MHz) of **6**

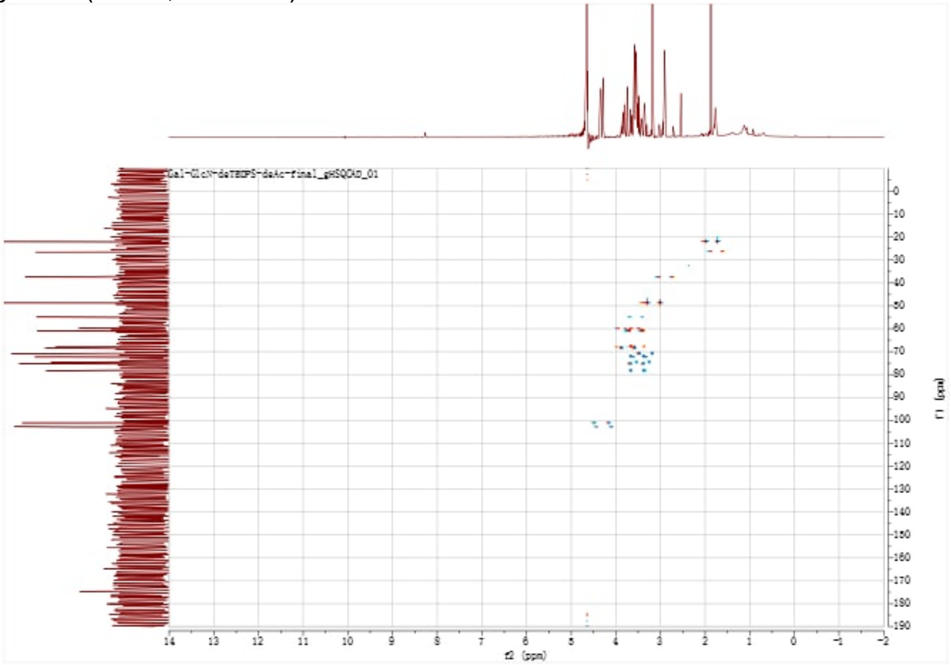


gHMBC($CDCl_3$, 500 MHz) of **6**

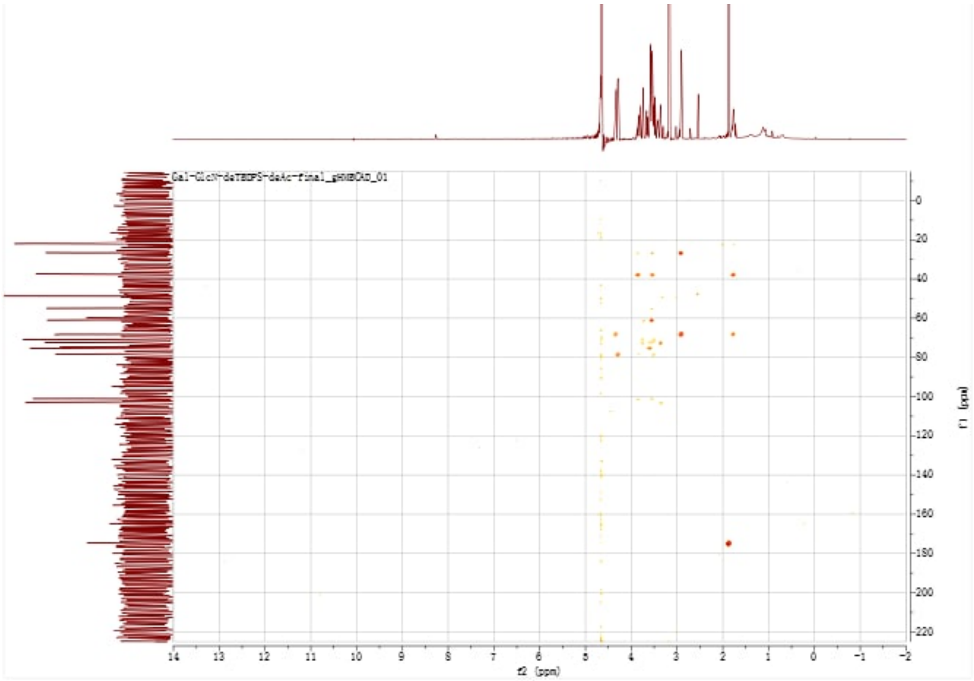

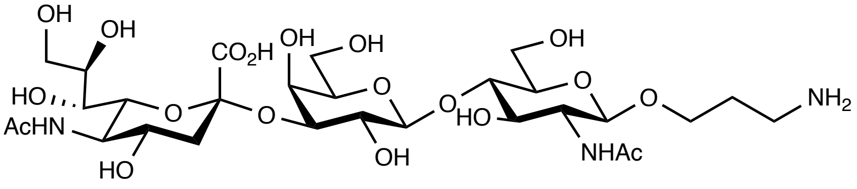

**7**

[1]H NMR ($D_2O$, v700 MHz) of **7**

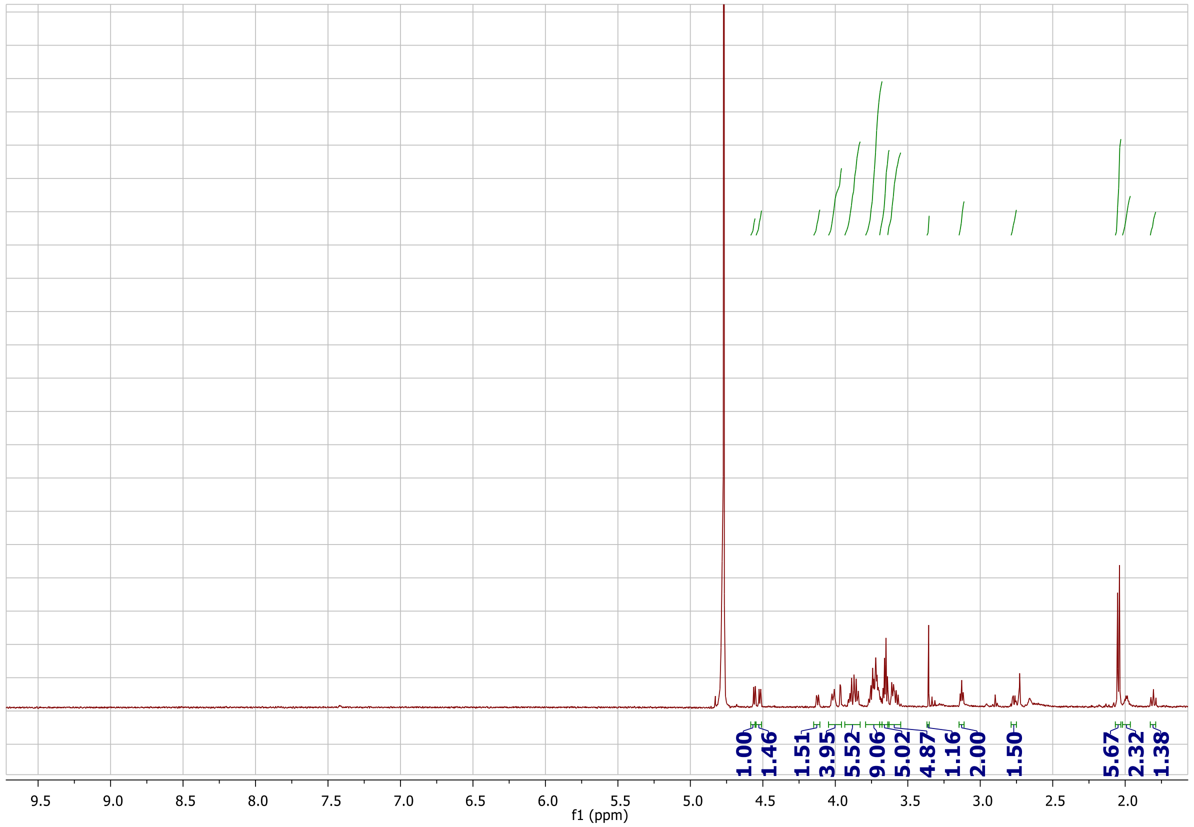

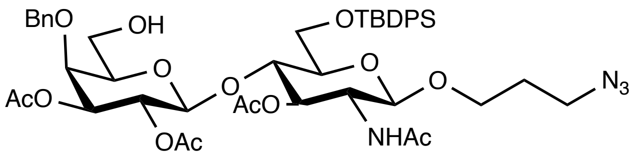


**8**

[1]H-NMR ($CDCl_3$, 500 MHz) of **8**

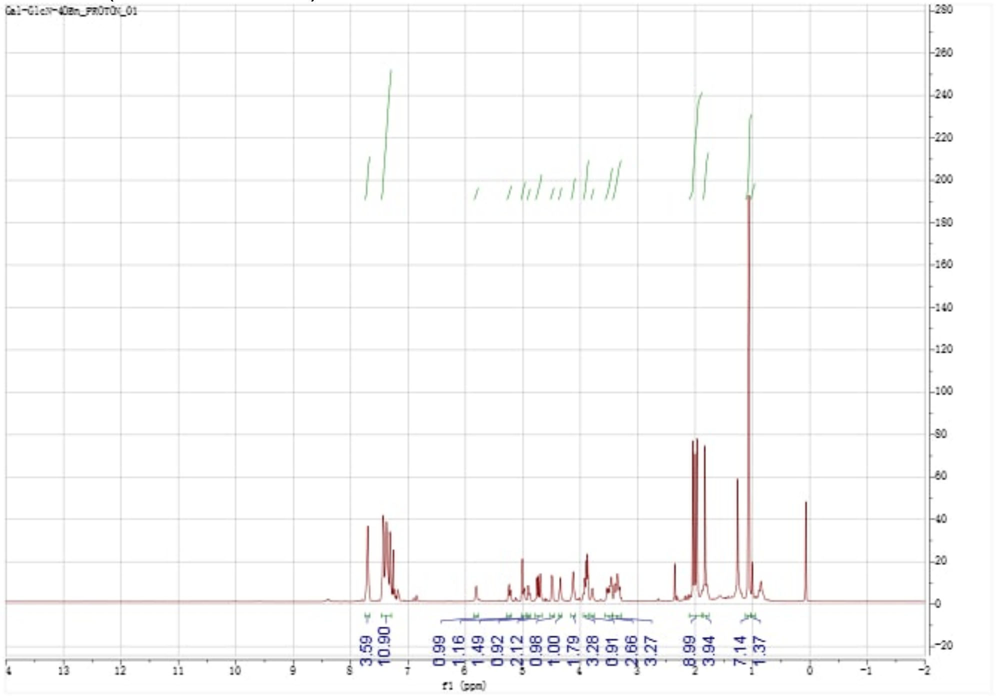

[13]C-NMR ($CDCl_3$, 126 MHz) of **8**

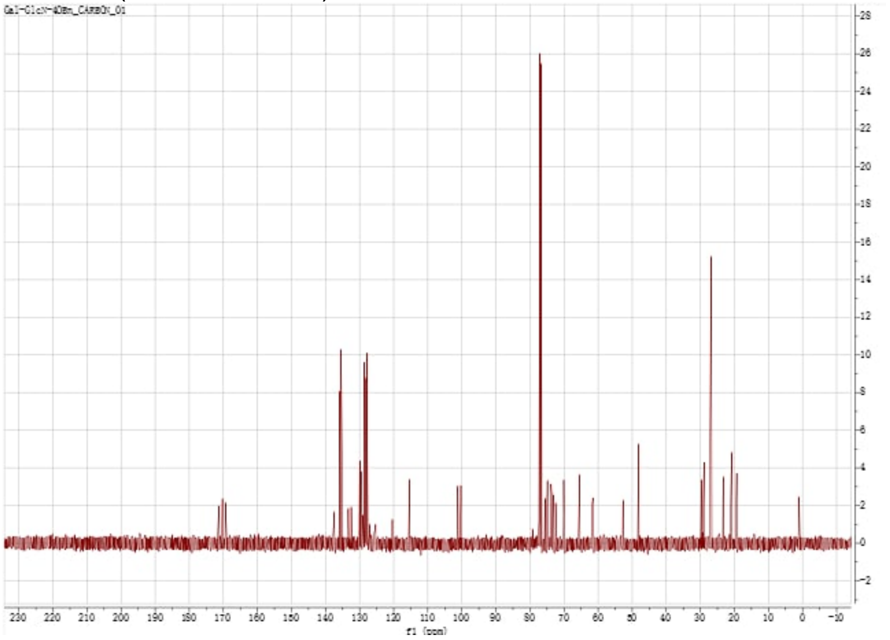

gCOSY($CDCl_3$, 500 MHz) of **8**

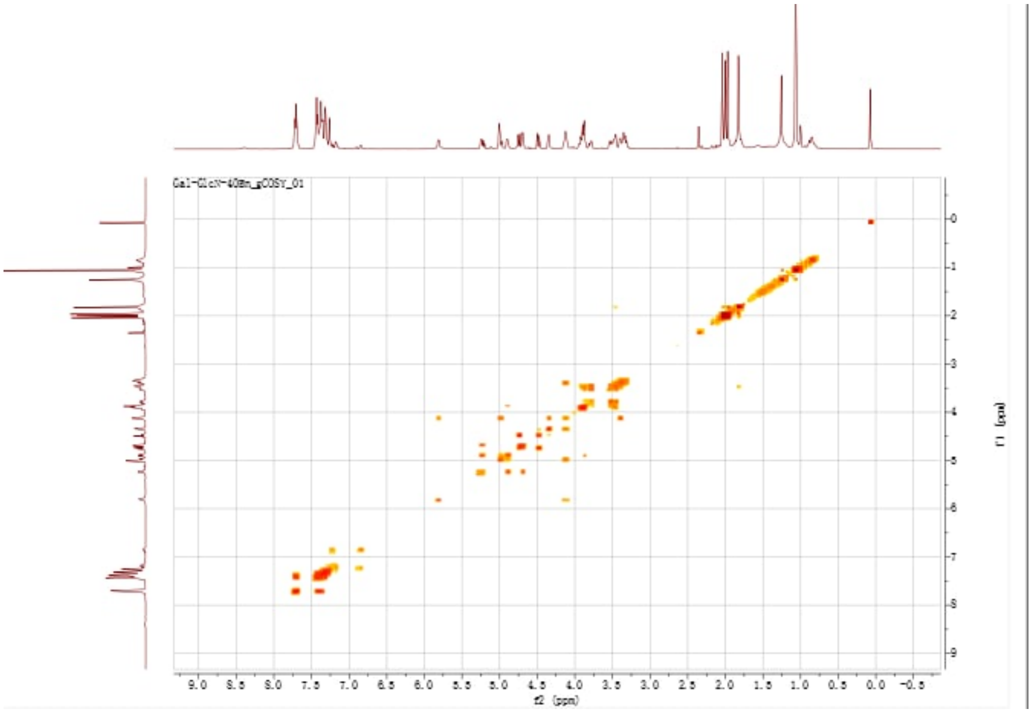


gHSQC($CDCl_3$, 500 MHz) of **8**

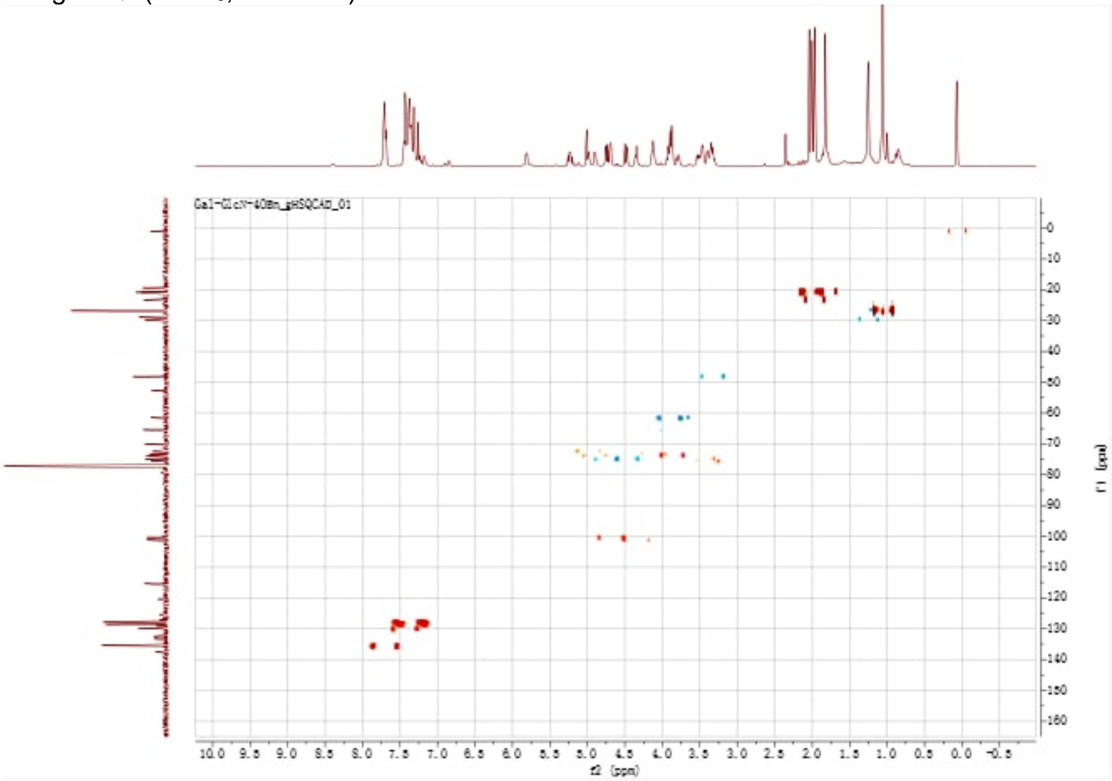


gHMBC($CDCl_3$, 500 MHz) of **8**

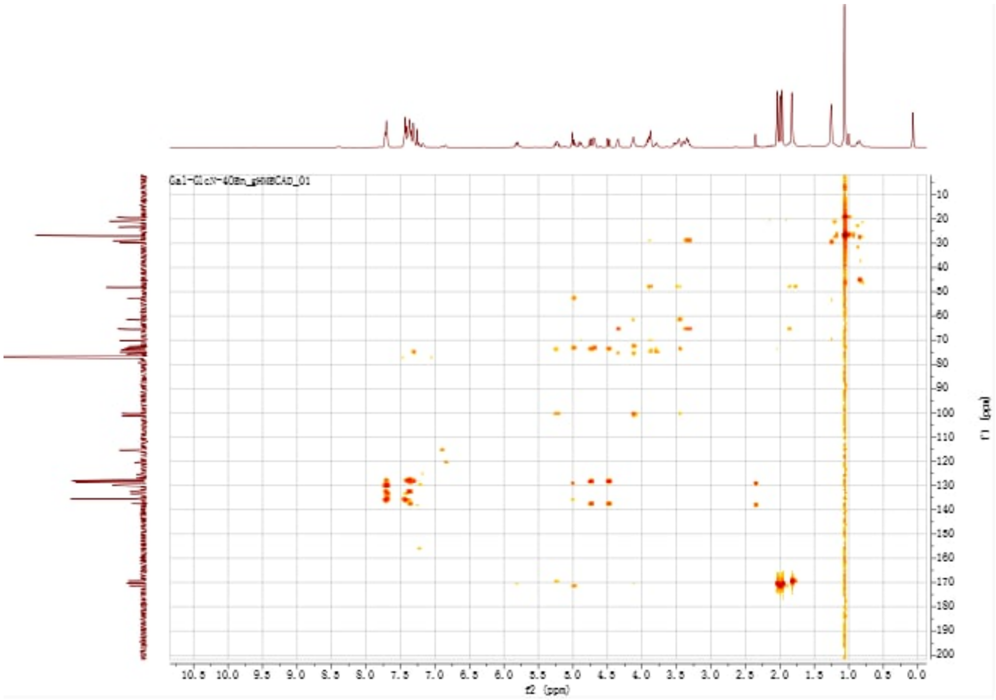

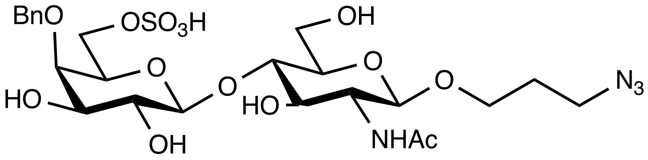

**9**

$^{1}$H-NMR (CDCl$_3$, 500 MHz) of **9**

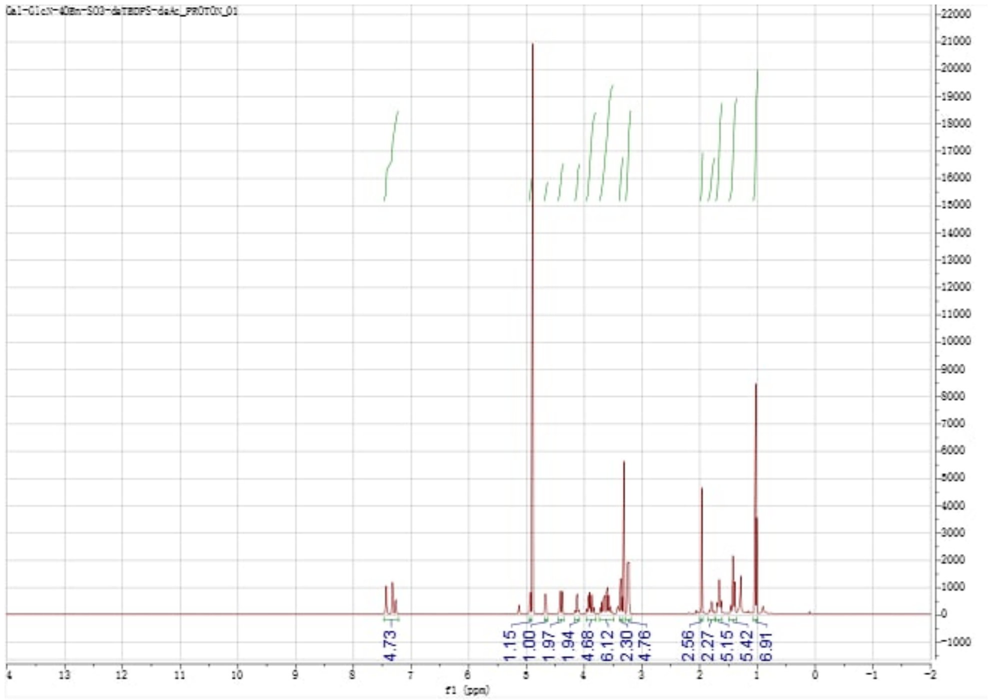


$^{13}$C-NMR ($CDCl_3$, 126 MHz) of **9**

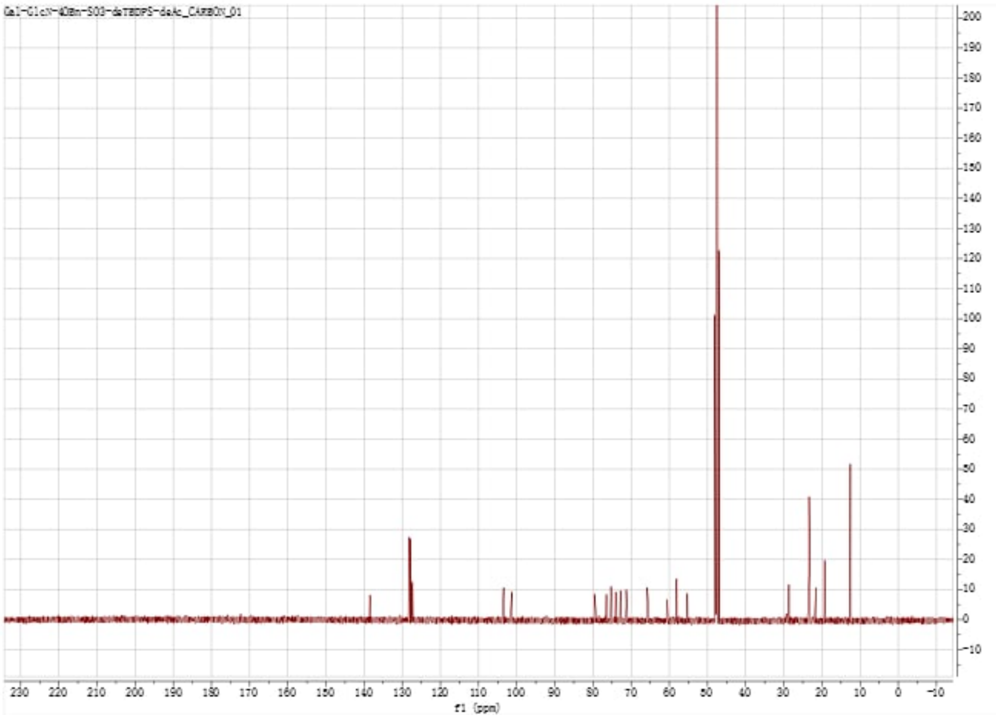


gCOSY($CDCl_3$, 500 MHz) of **9**

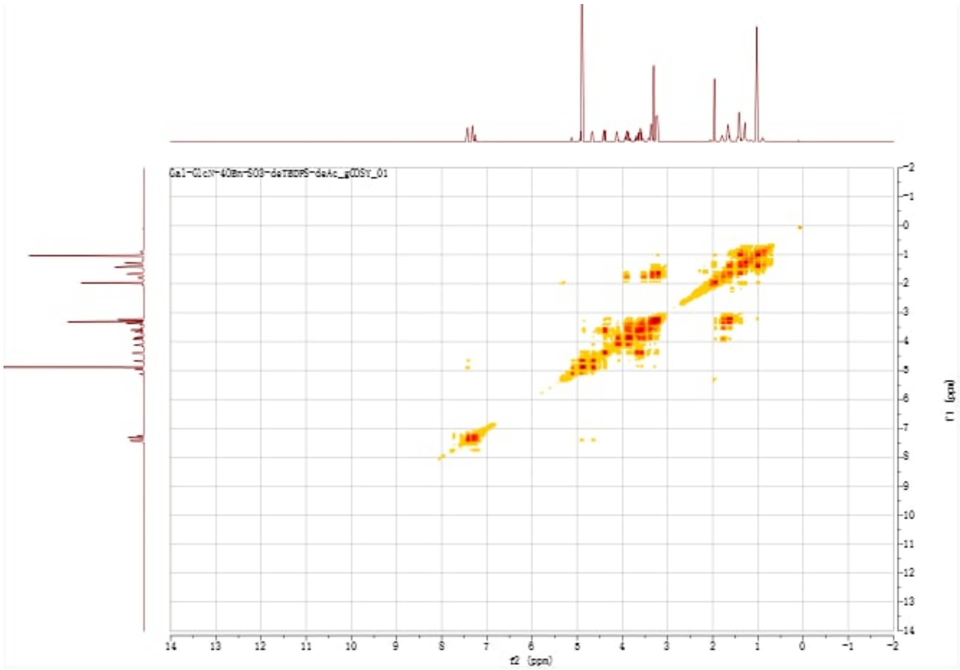


gHSQC($CDCl_3$, 500 MHz) of **9**

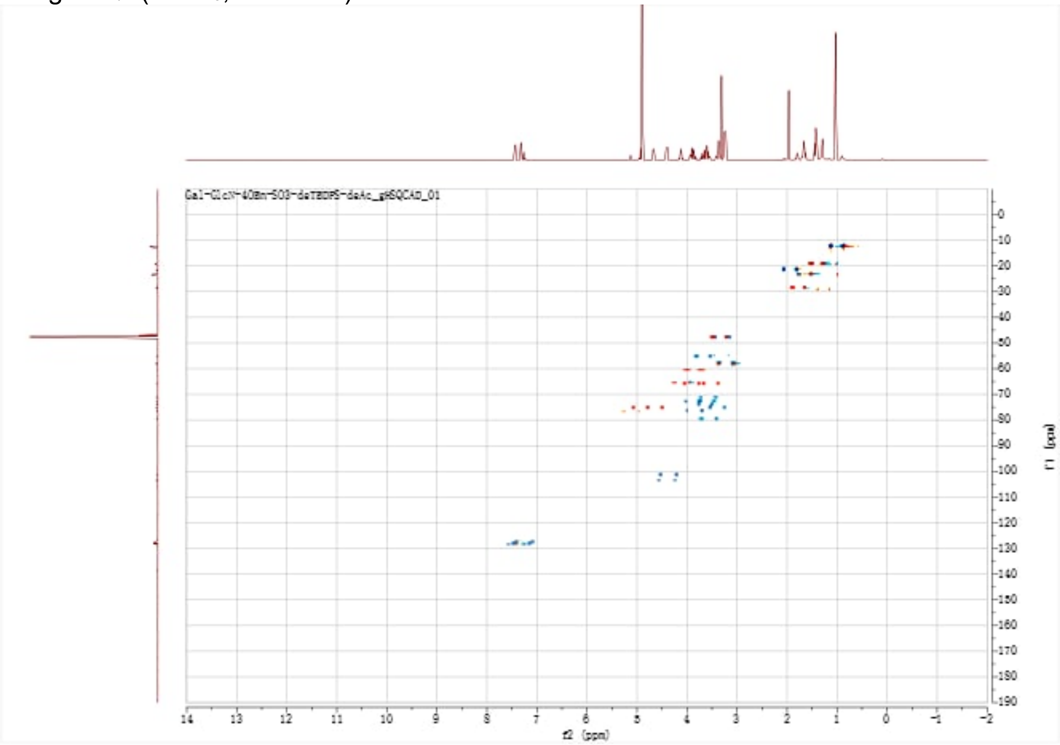


gHMBC($CDCl_3$, 500 MHz) of **9**

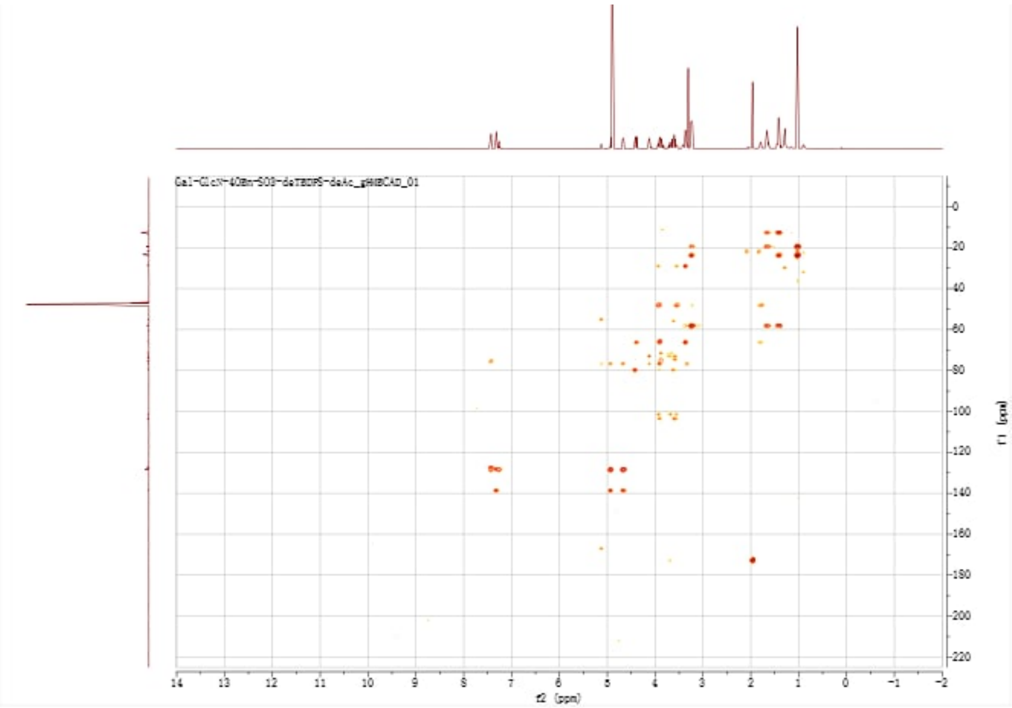


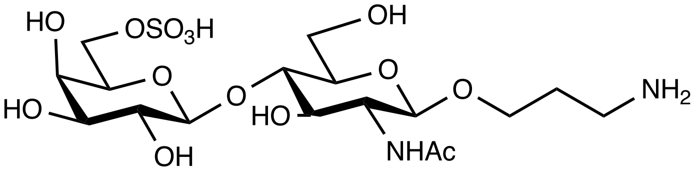


**10**

[1]H-NMR (CDCl₃, 500 MHz) of **10**

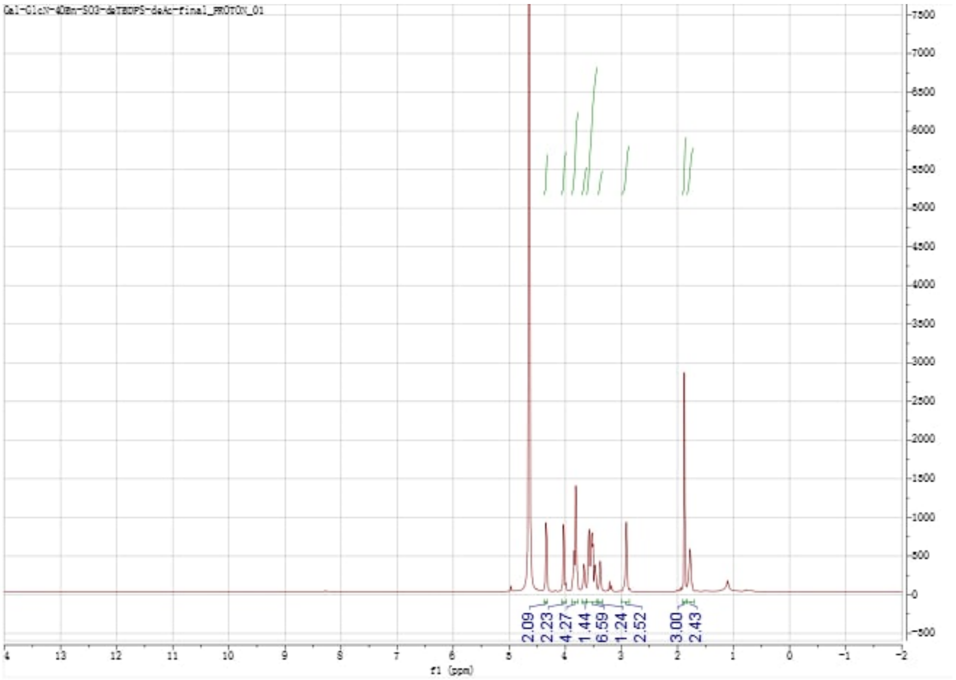

$^{13}C$-NMR ($CDCl_3$, 126 MHz) of **10**

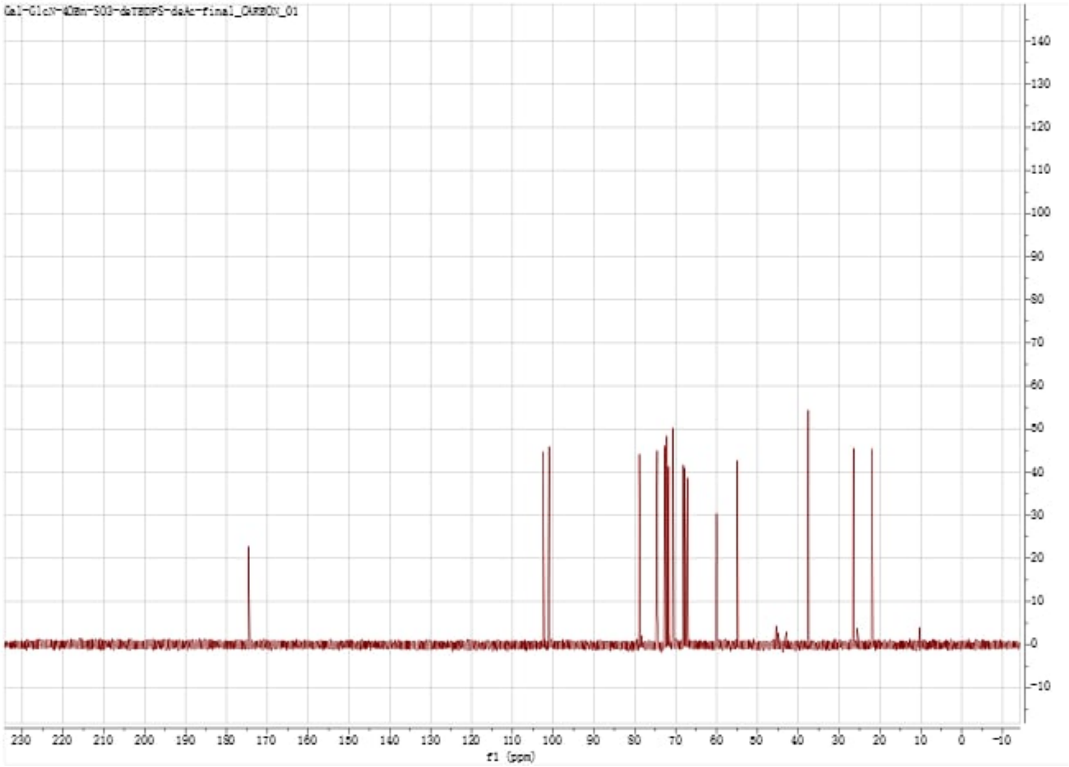


gCOSY($CDCl_3$, 500 MHz) of **10**

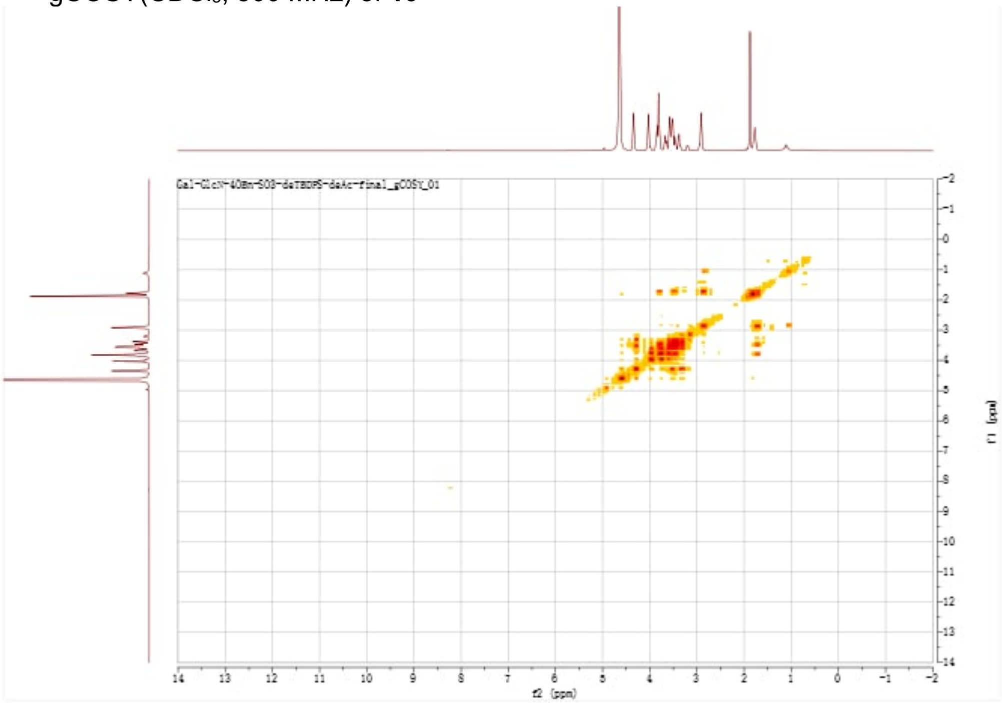


gHSQC($CDCl_3$, 500 MHz) of **10**

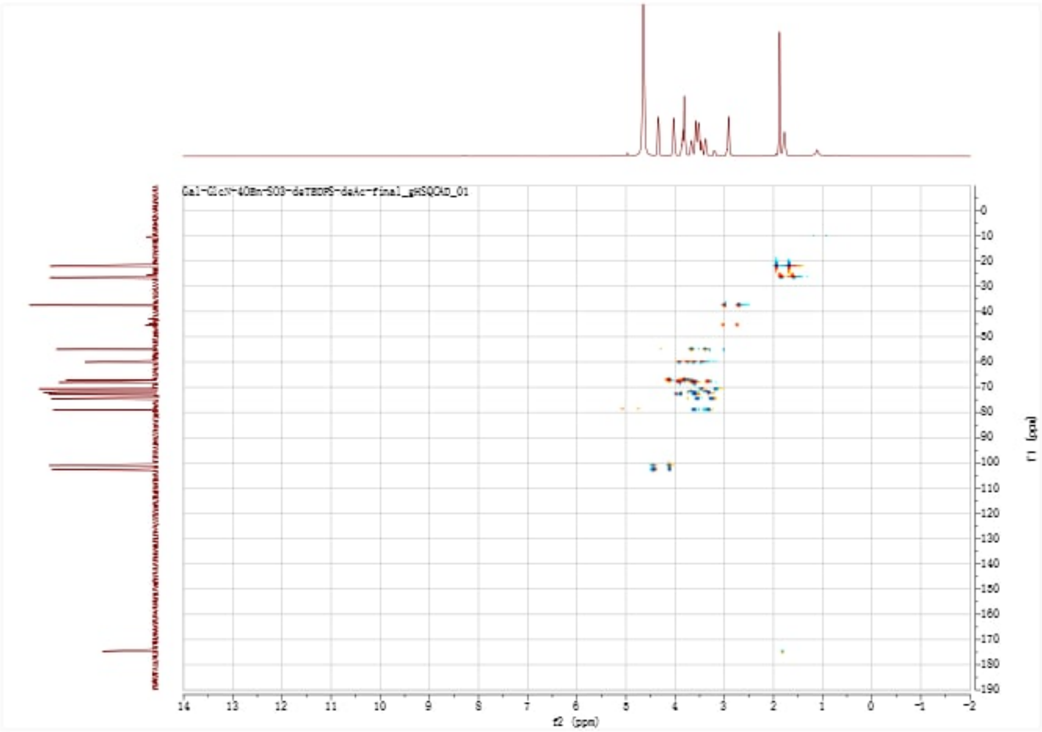

gHMBC($CDCl_3$, 500 MHz) of **10**

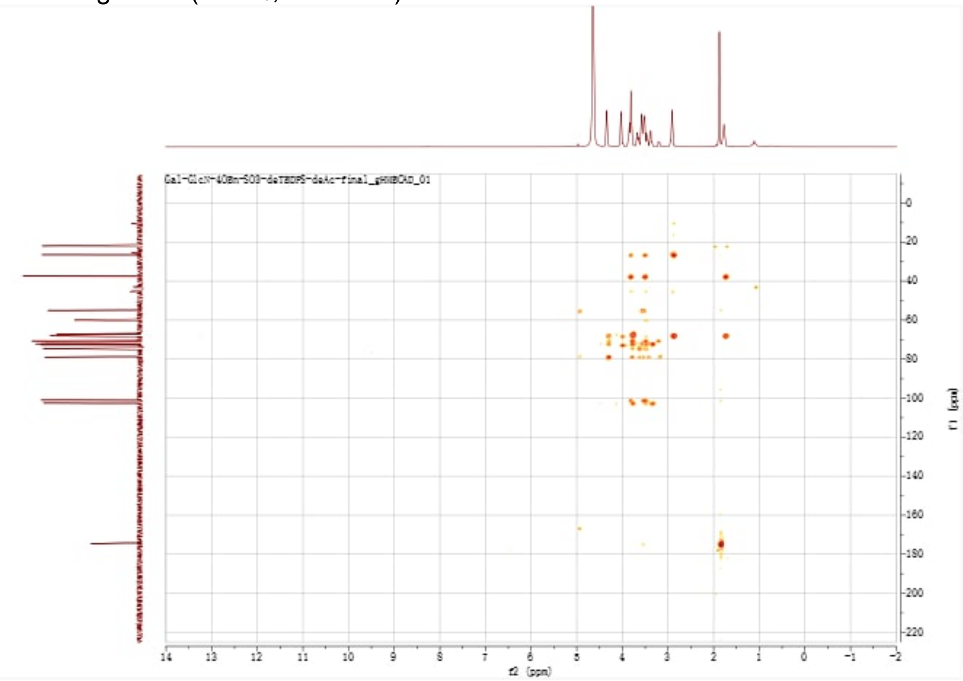

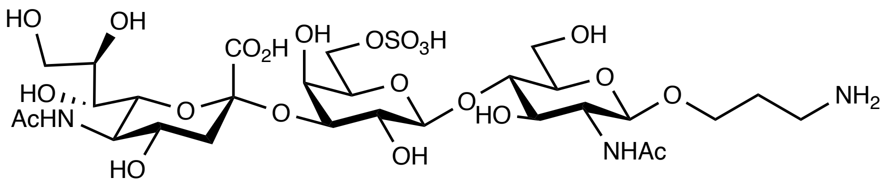


**11**

$^{1}$H NMR ($D_2O$, v700 MHz) of **11**

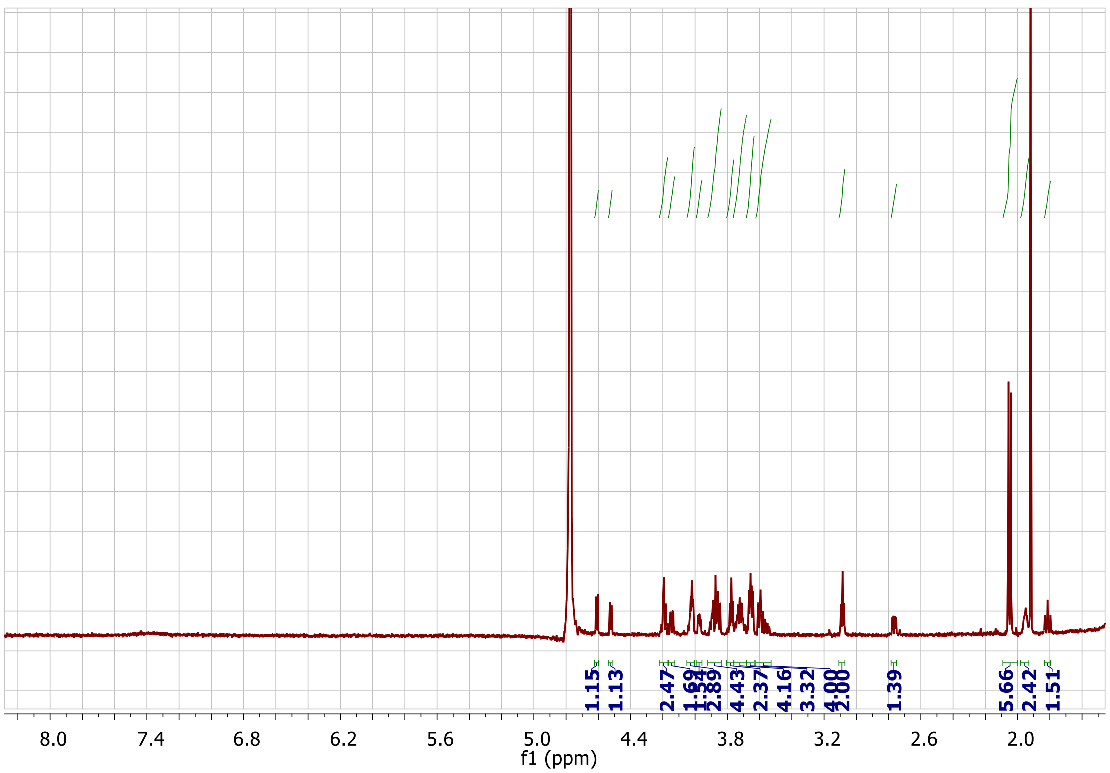


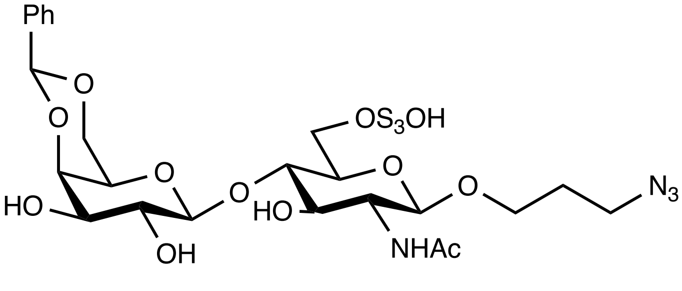


**12**

$^{1}$H-NMR ($CDCl_3$, 500 MHz) of **12**

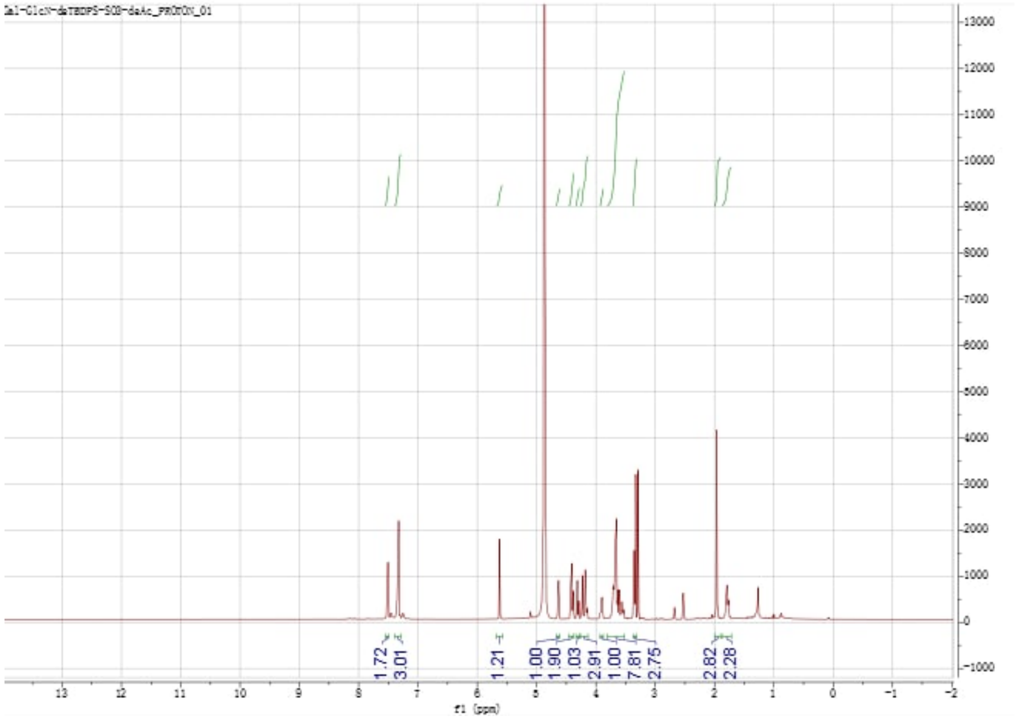


$^{13}$C-NMR ($CDCl_3$, 126 MHz) of **12**

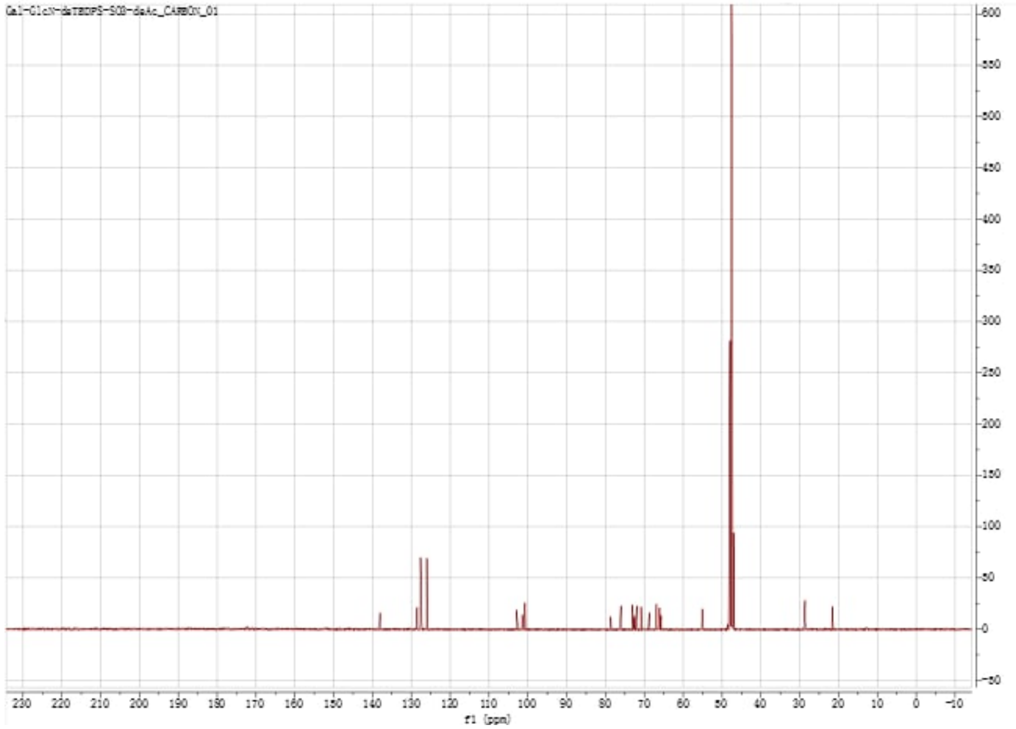

gCOSY(CDCl$_3$, 500 MHz) of **12**

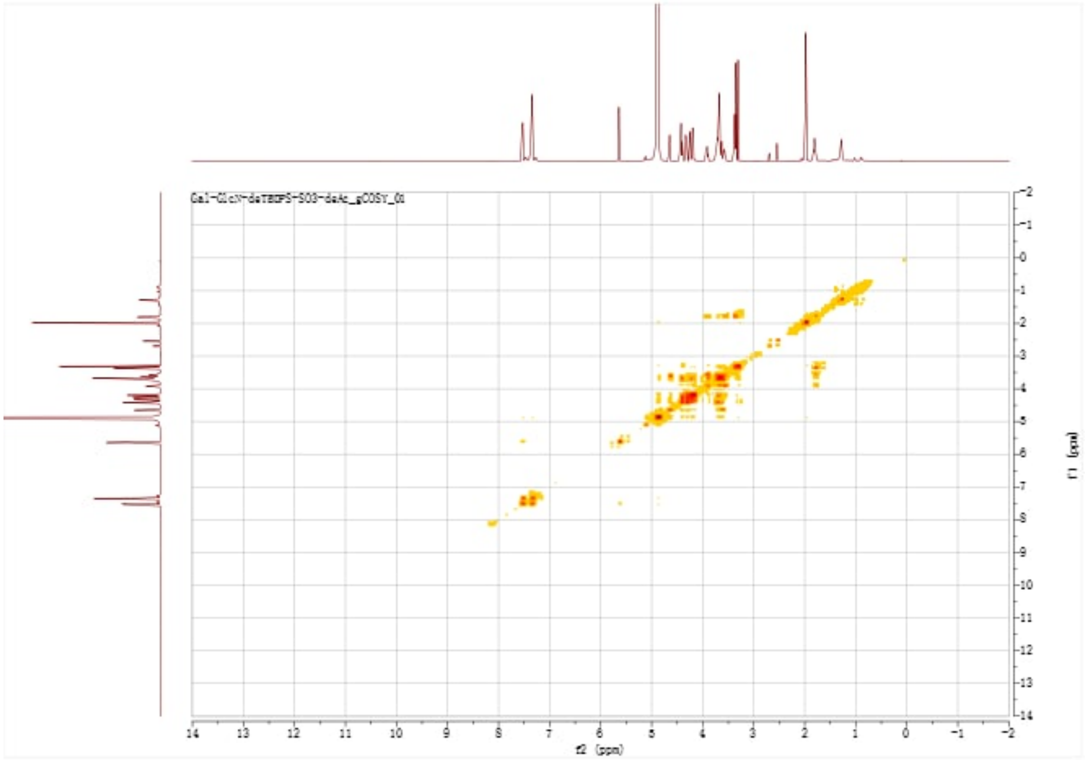

gHSQC(CDCl$_3$, 500 MHz) of **12**

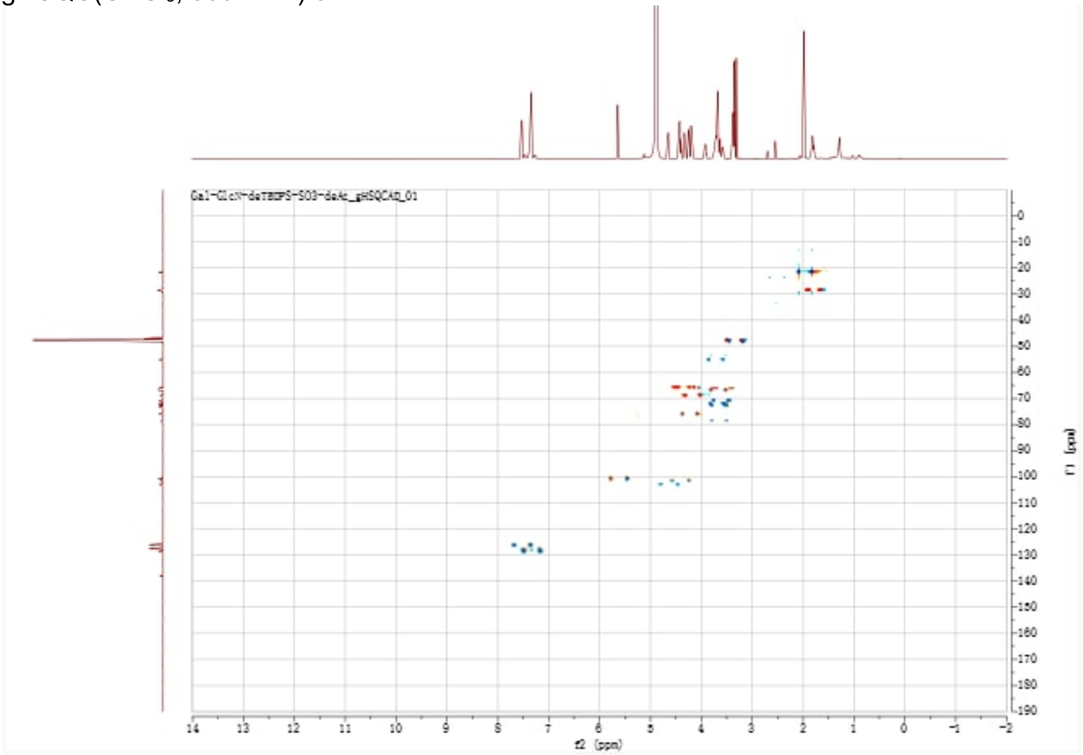

gHMBC($CDCl_3$, 500 MHz) of **12**

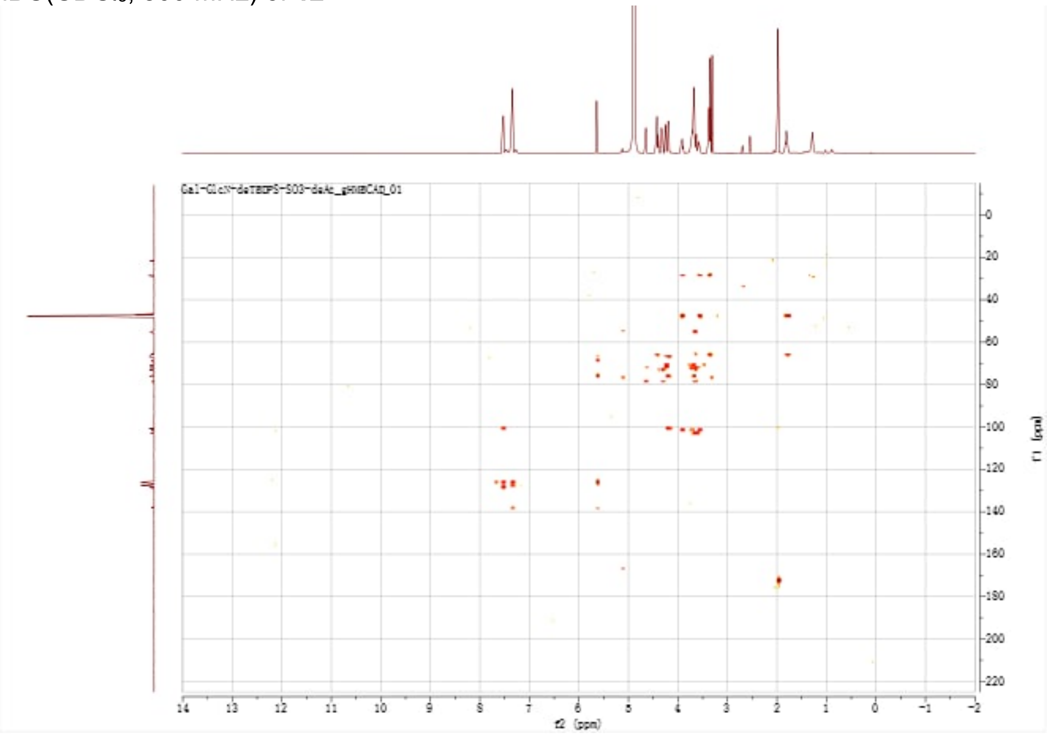

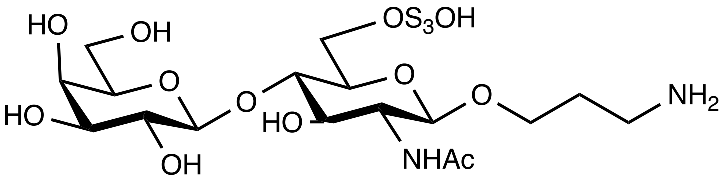


**13**

[1]H-NMR ($CDCl_3$, 500 MHz) of **13**

[13]C-NMR ($CDCl_3$, 126 MHz) of **13**

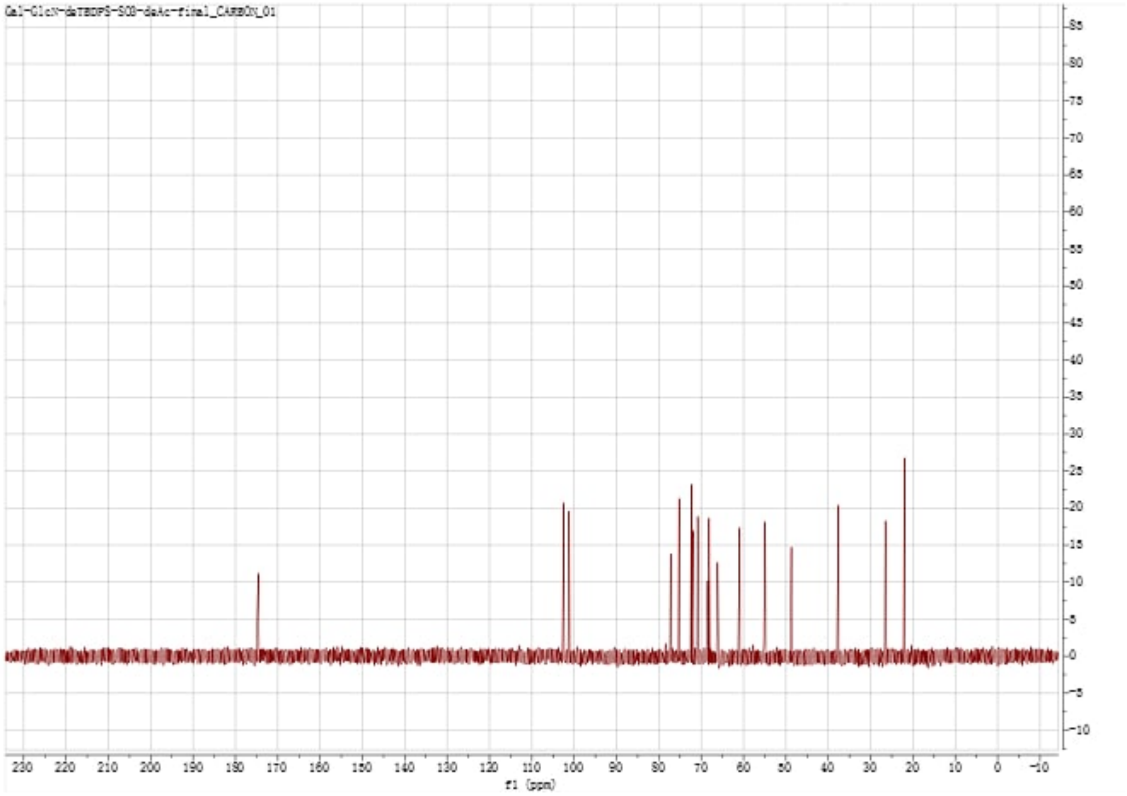

gCOSY(CDCl$_3$, 500 MHz) of **13**

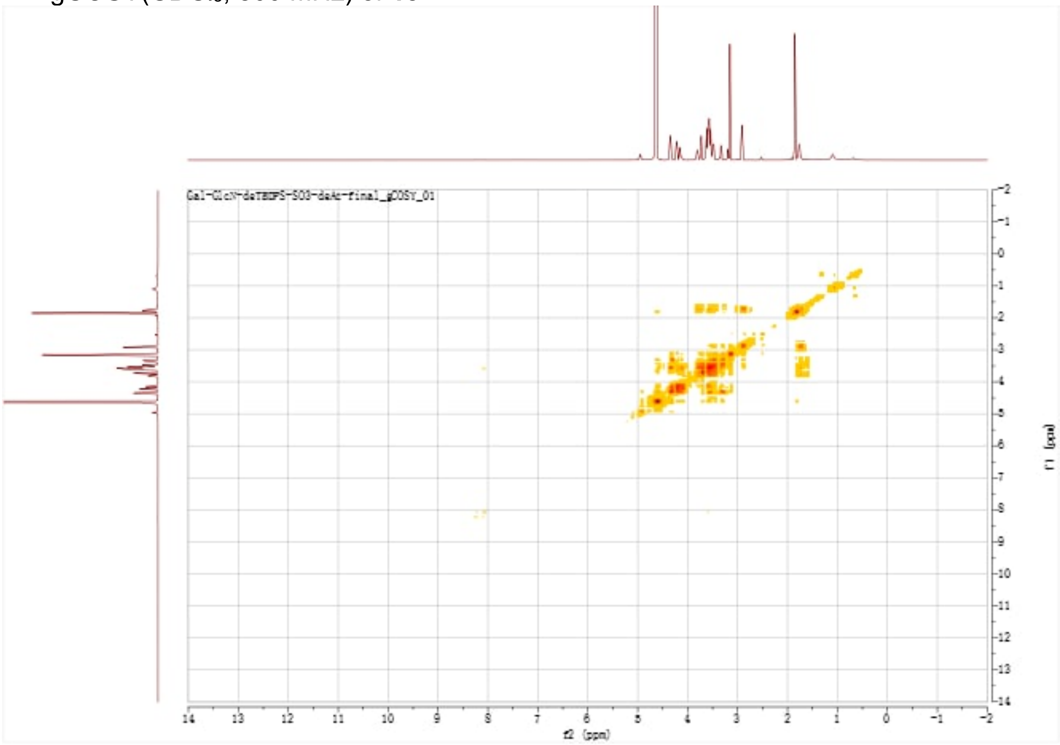


gHSQC(CDCl$_3$, 500 MHz) of **13**

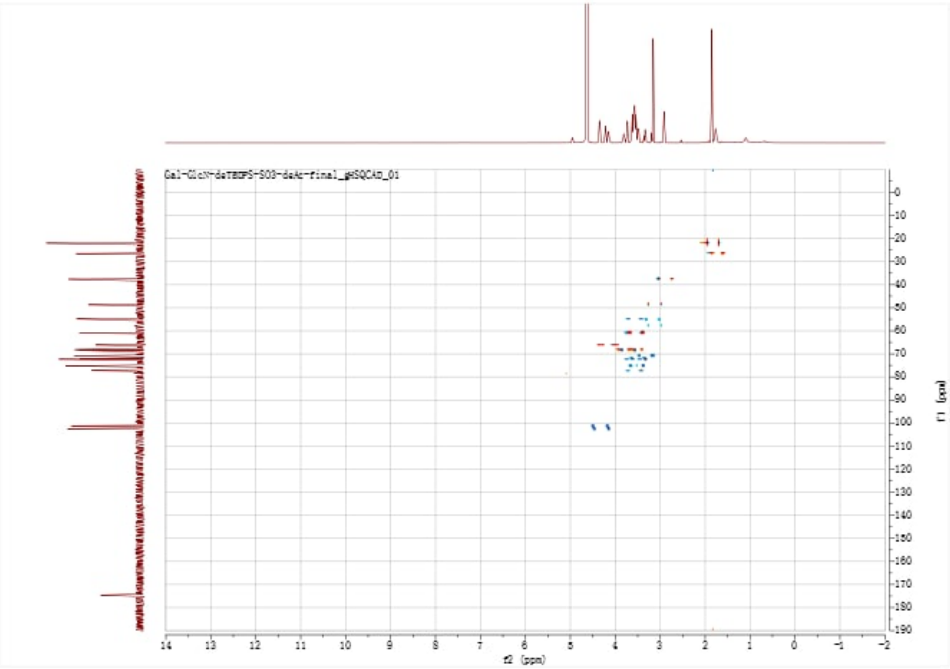


gHMBC(CDCl$_3$, 500 MHz) of **13**

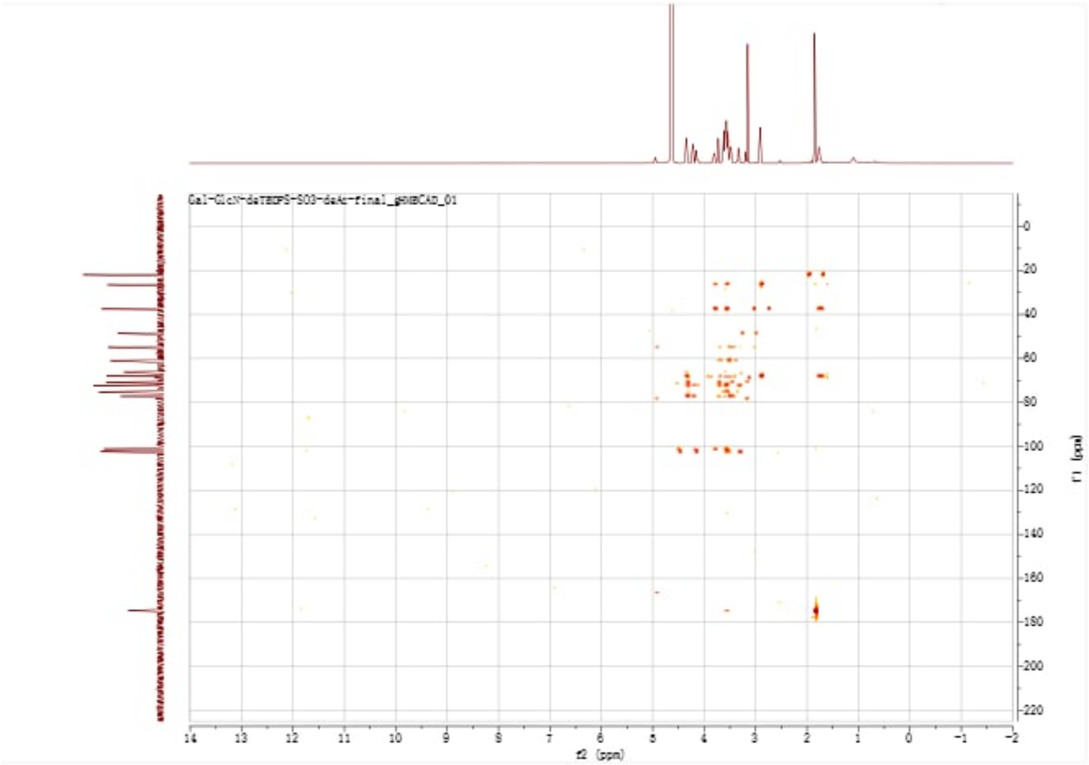


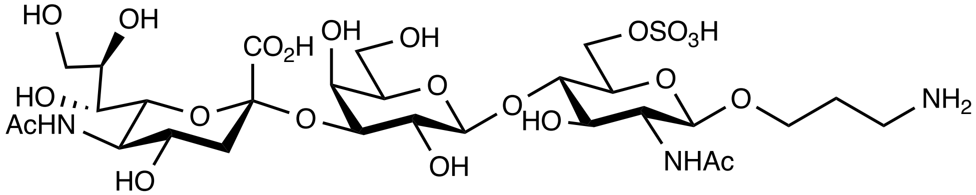


**14**

$^{1}$H NMR ($D_2O$, v700 MHz) of **14**

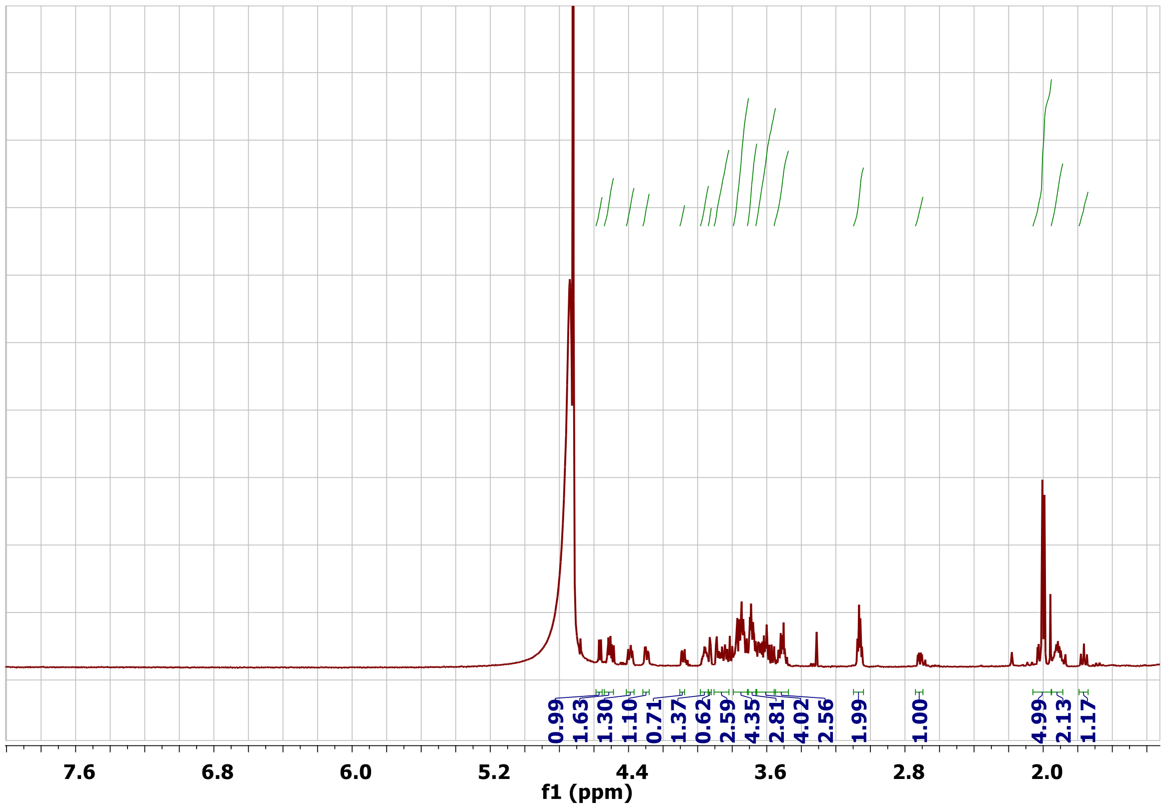


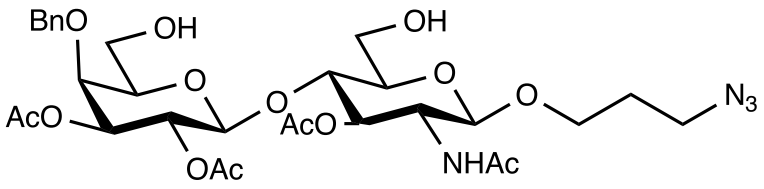


**15**

[1]H-NMR ($CDCl_3$, 500 MHz) of **15**

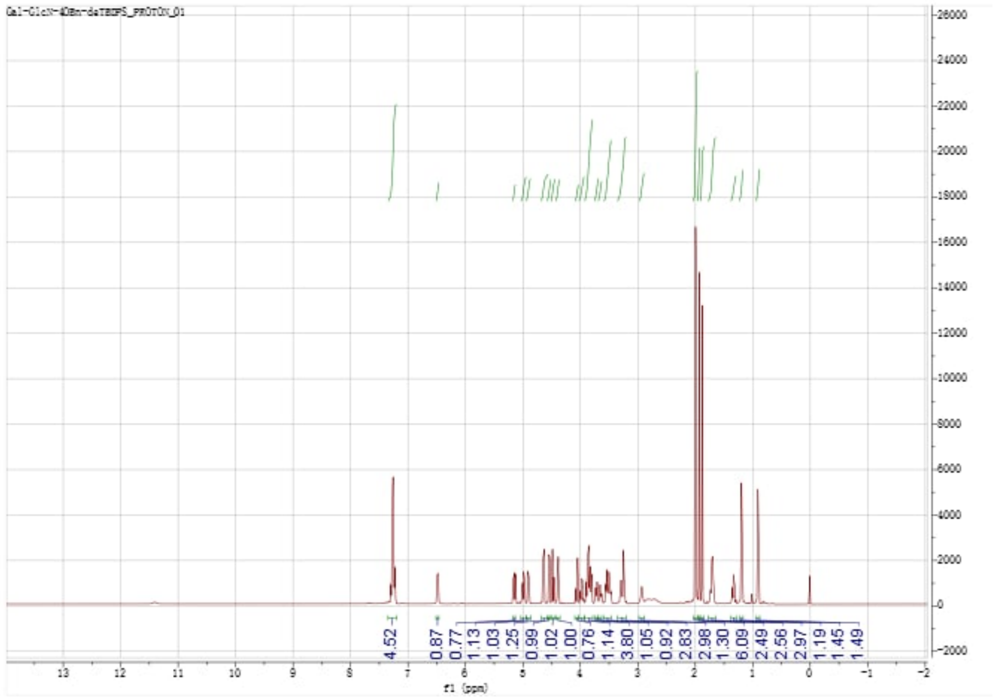


$^{13}$C-NMR ($CDCl_3$, 126 MHz) of **15**

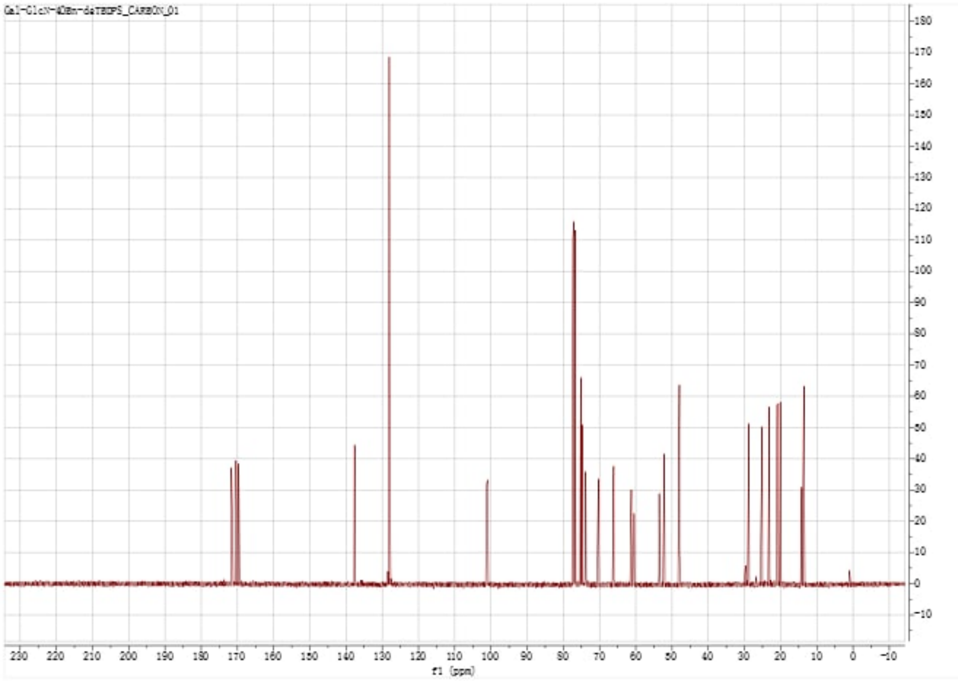

gCOSY(CDCl$_3$, 500 MHz) of **15**

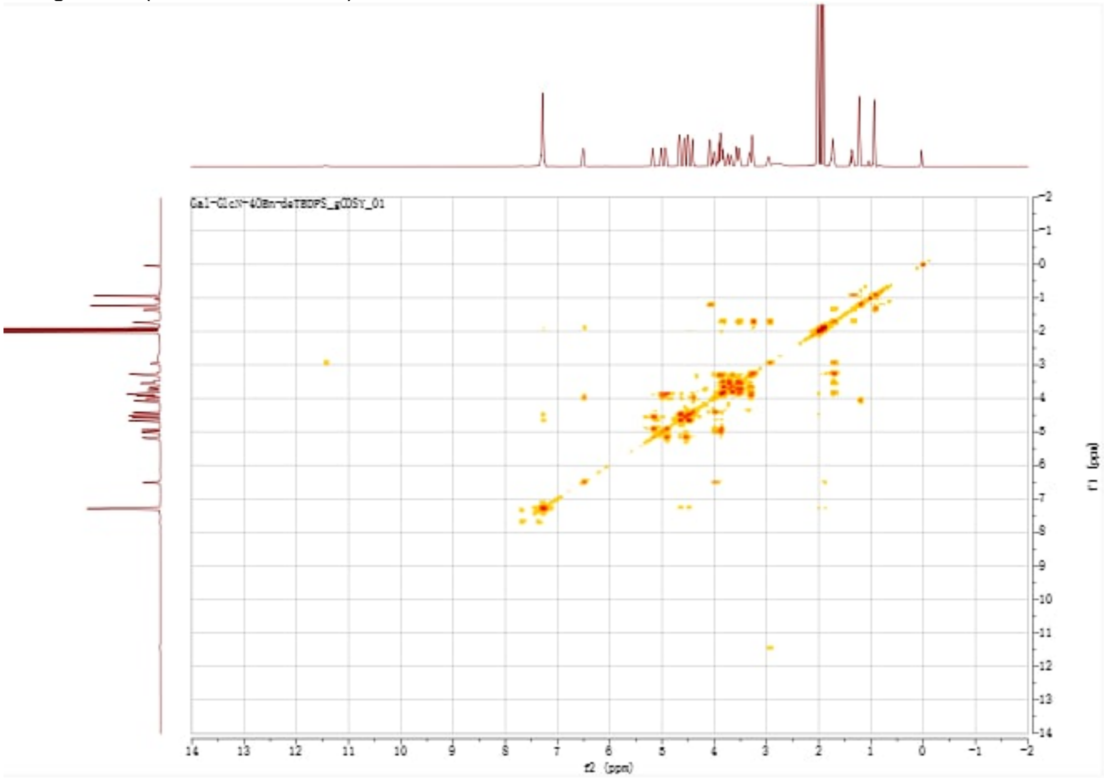


gHSQC(CDCl$_3$, 500 MHz) of **15**

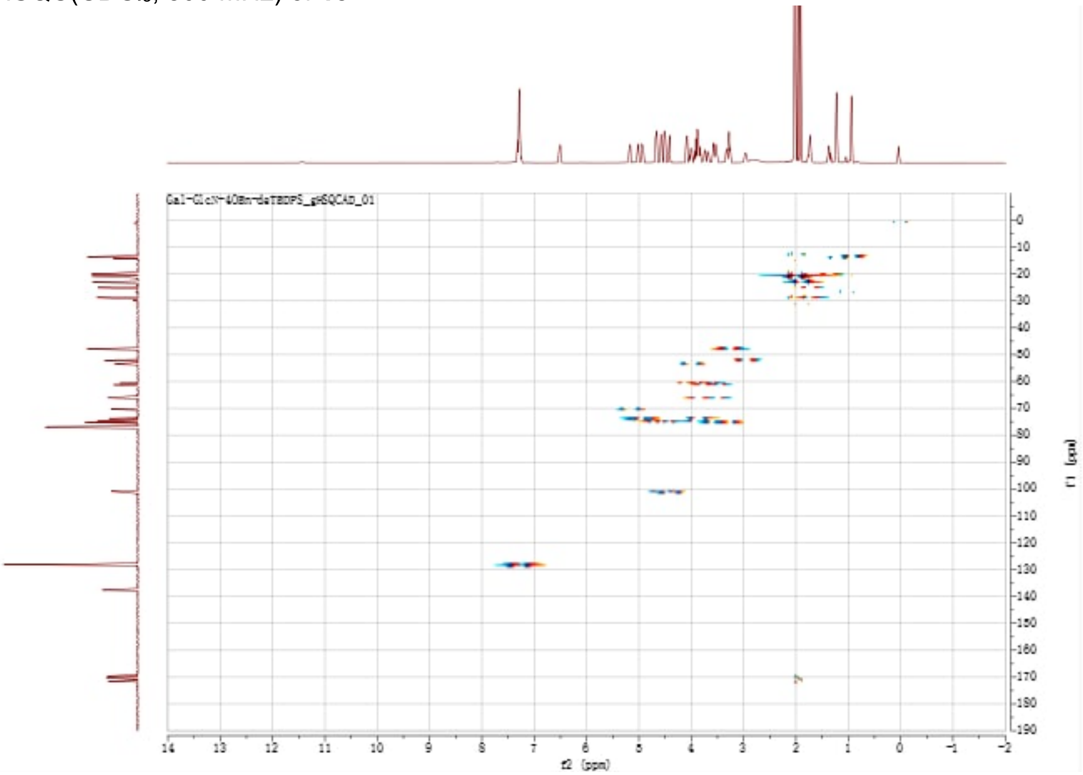

gHMBC($CDCl_3$, 500 MHz) of **15**

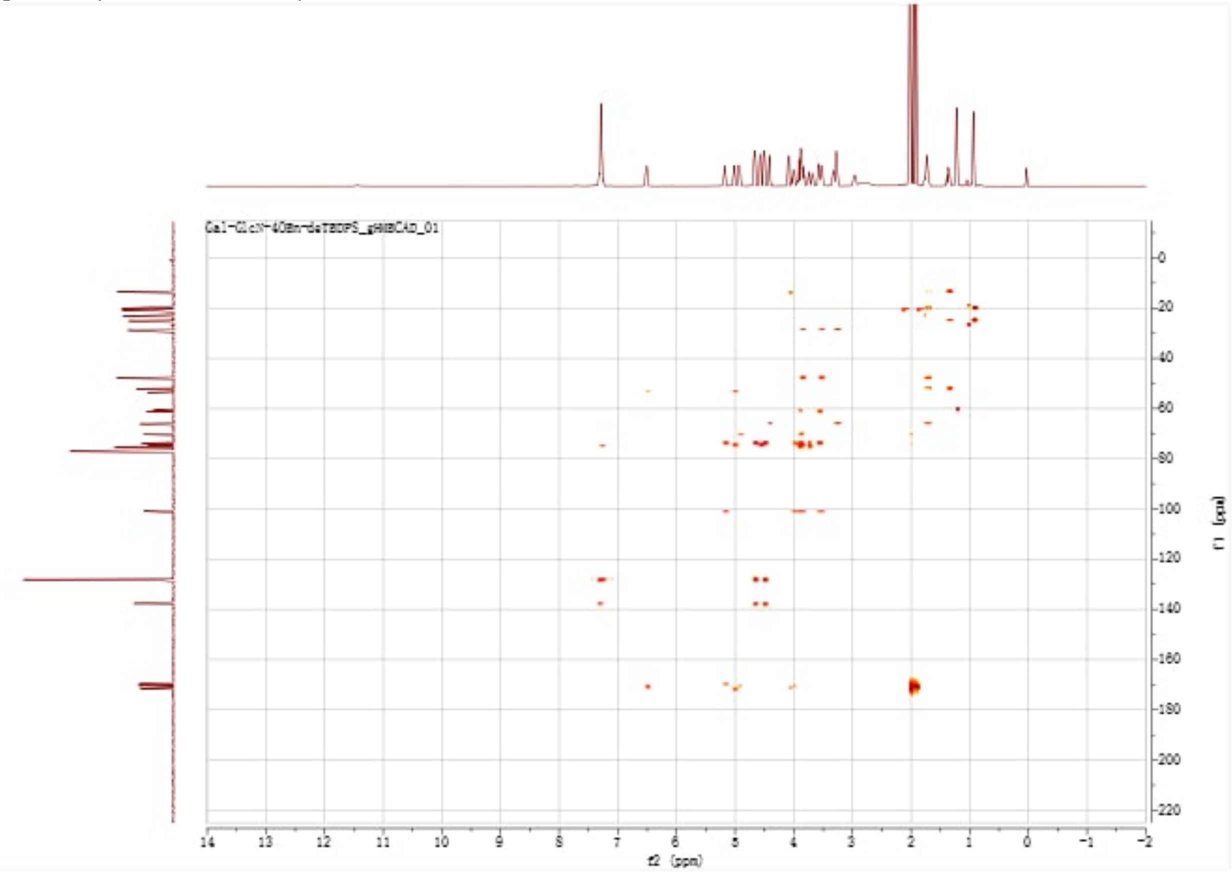

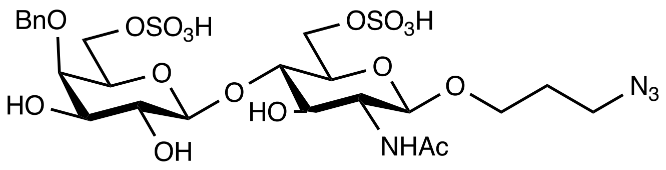


**16**

[1]H-NMR ($CDCl_3$, 500 MHz) of **16**

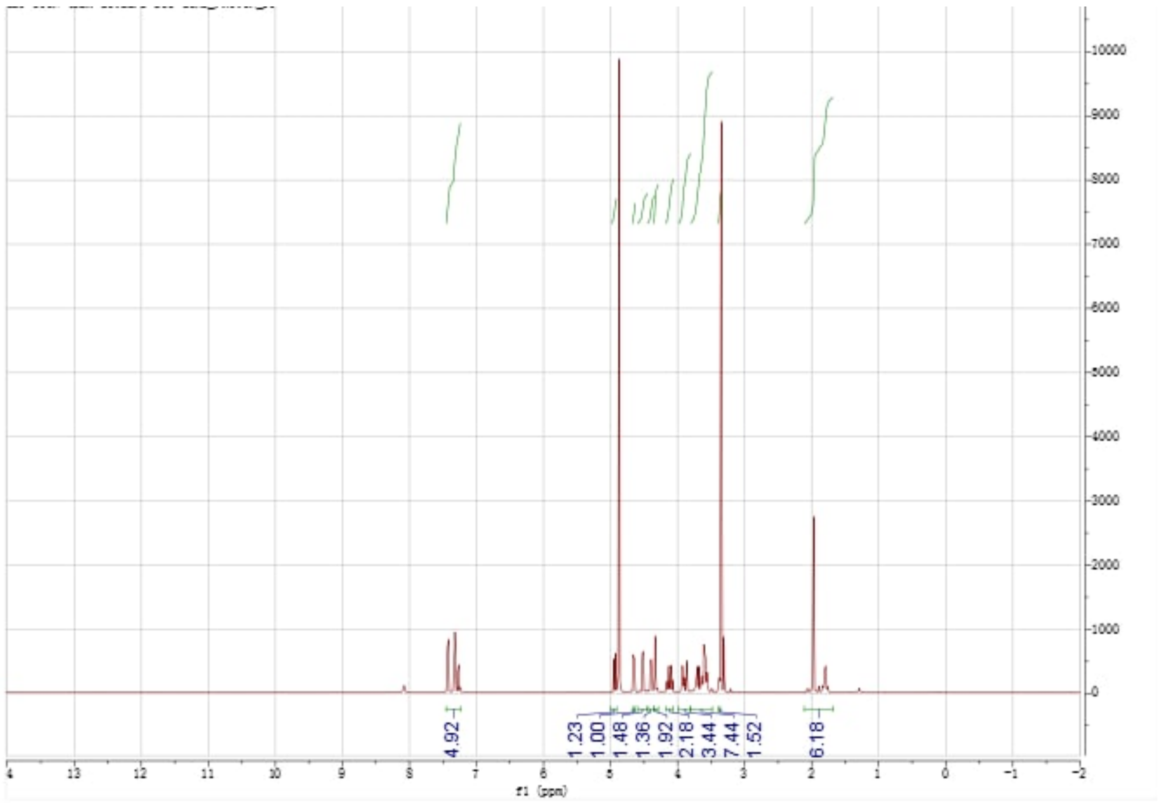

$^{13}$C-NMR ($CDCl_3$, 126 MHz) of **16**

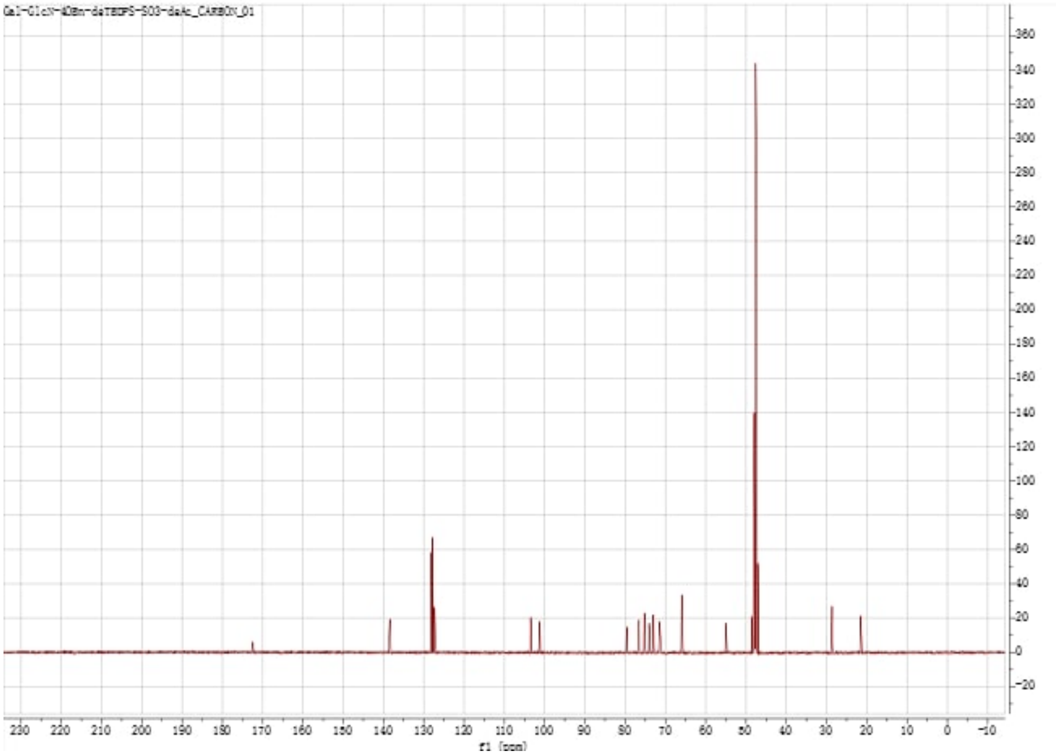


gCOSY($CDCl_3$, 500 MHz) of **16**

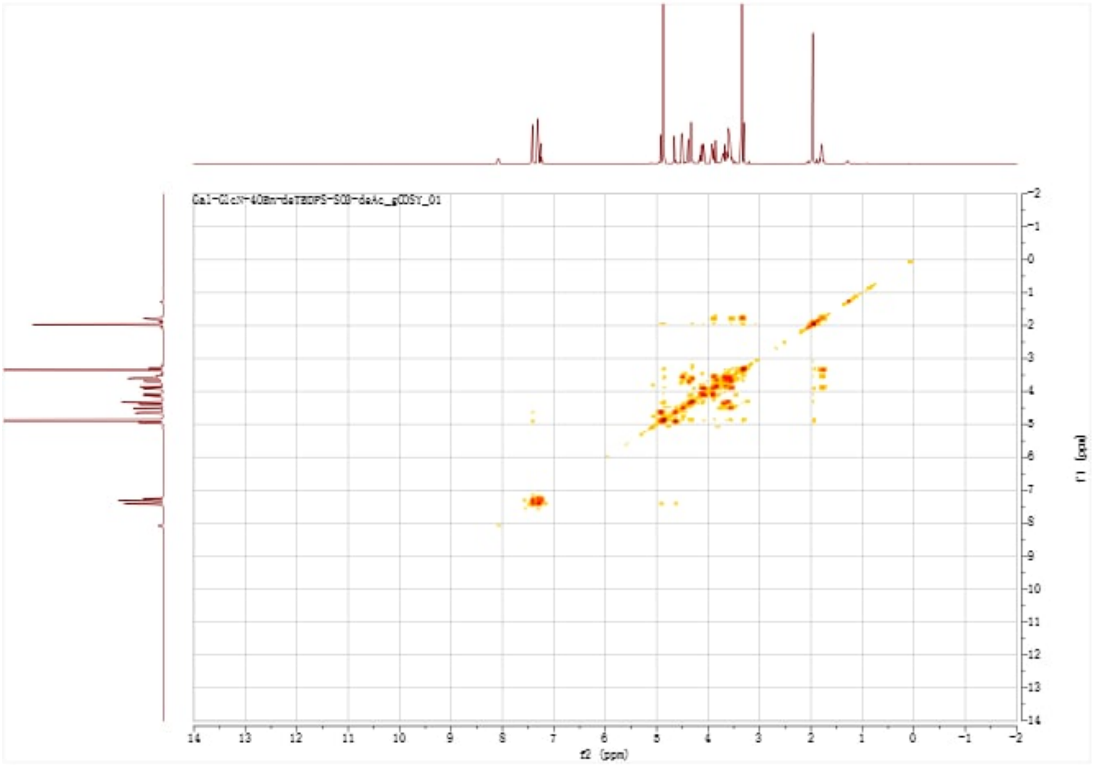

gHSQC($CDCl_3$, 500 MHz) of **16**

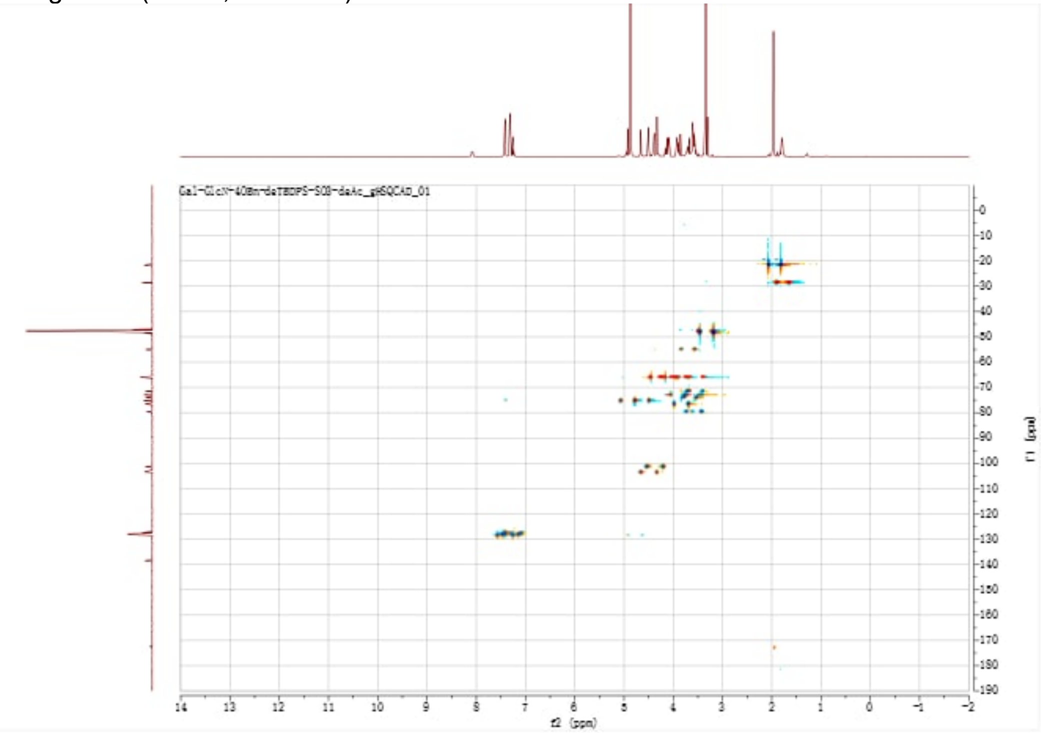


gHMBC($CDCl_3$, 500 MHz) of **16**

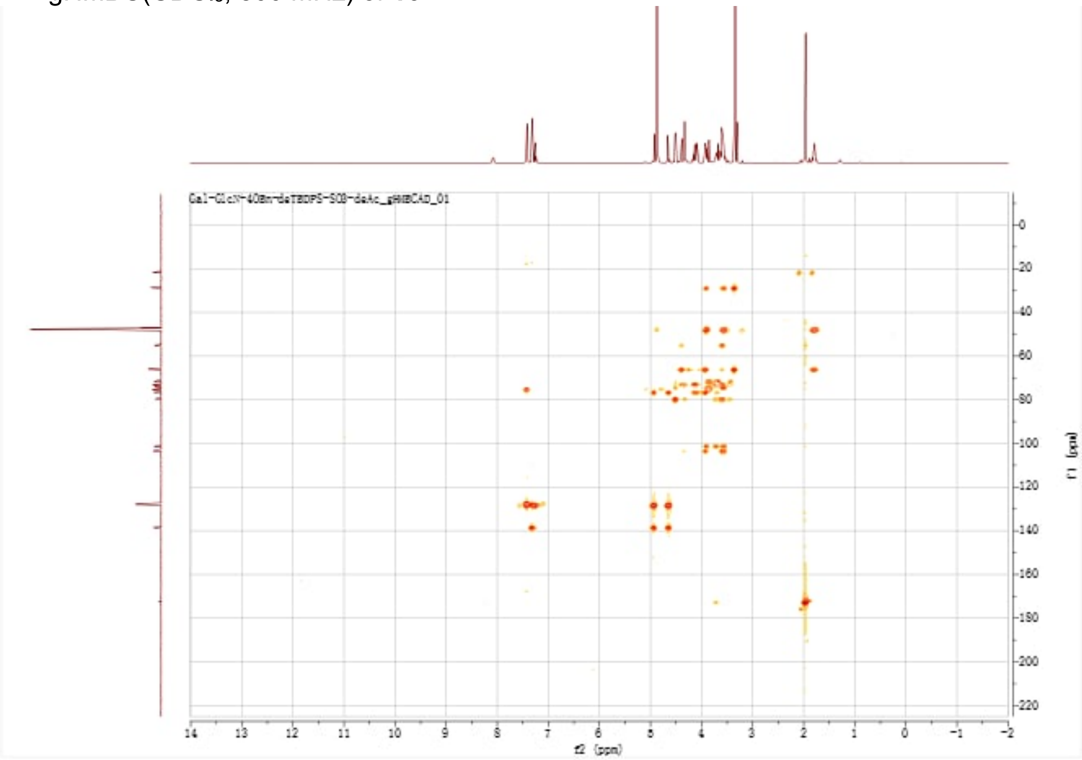

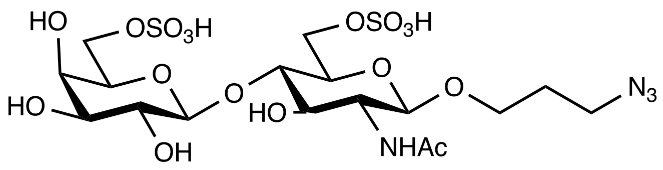

**17**

[1]H-NMR ($CDCl_3$, 500 MHz) of **17**

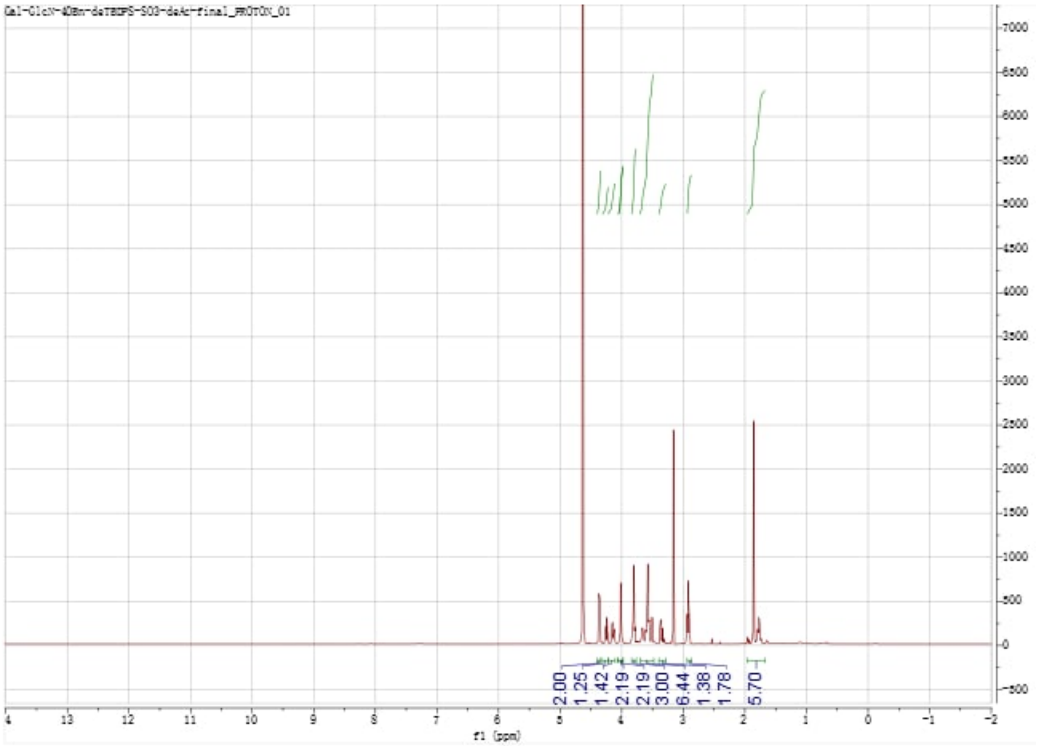

$^{13}$C-NMR ($CDCl_3$, 126 MHz) of **17**

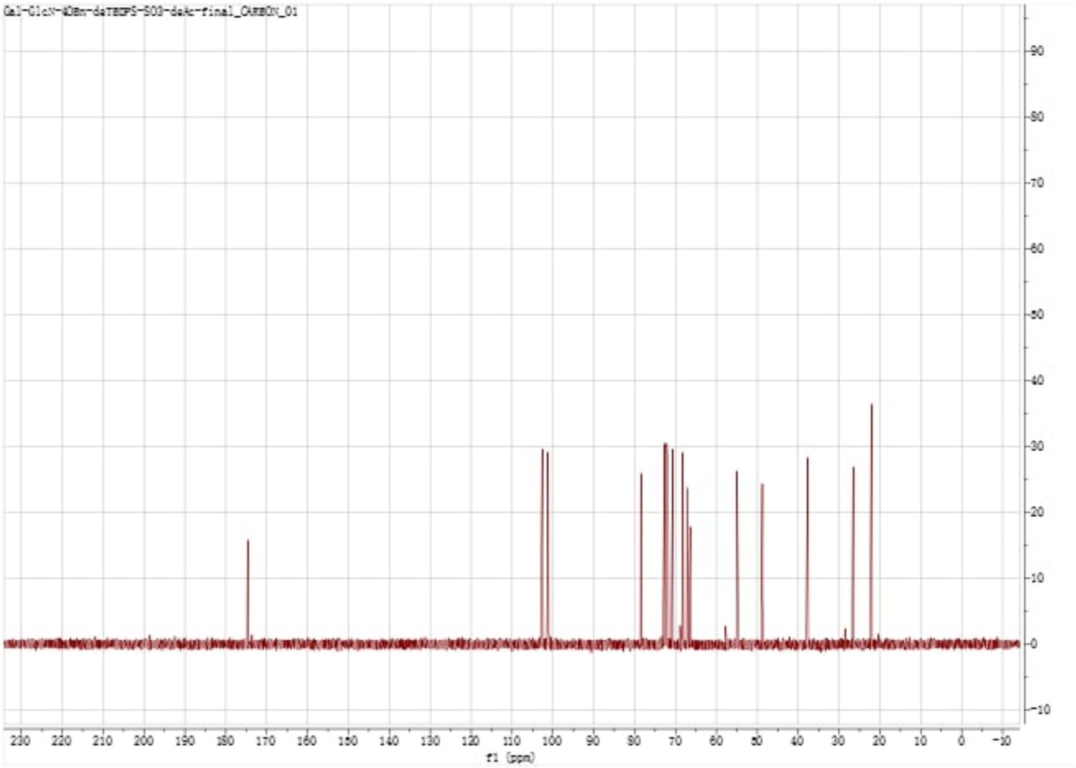


gCOSY($CDCl_3$, 500 MHz) of **17**

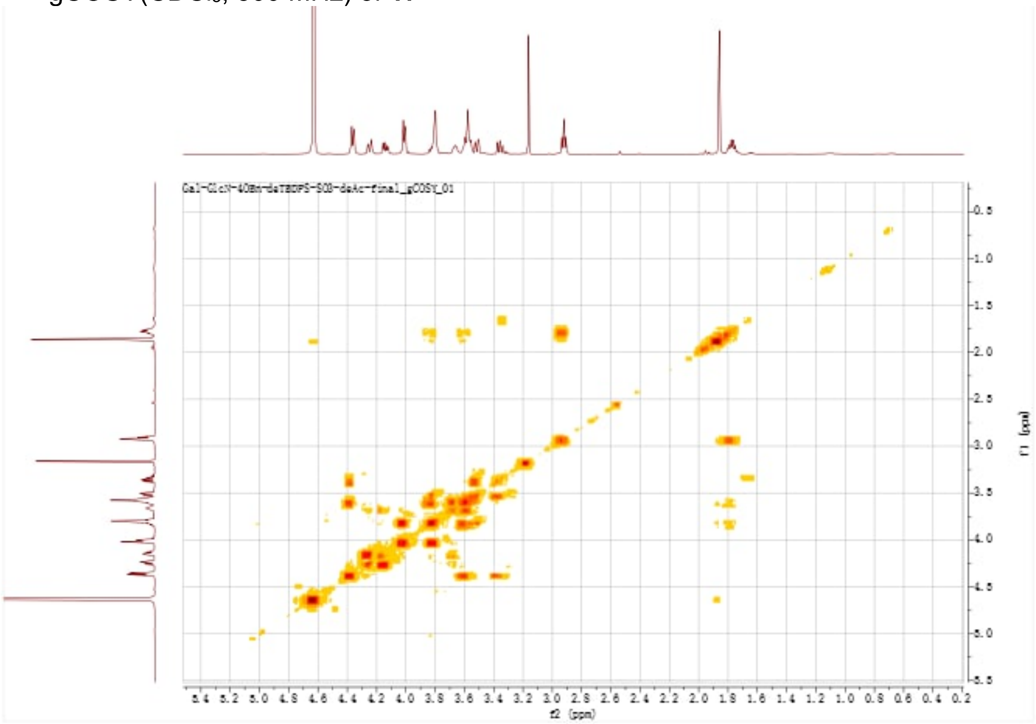

gHSQC(CDCl$_3$, 500 MHz) of **17**

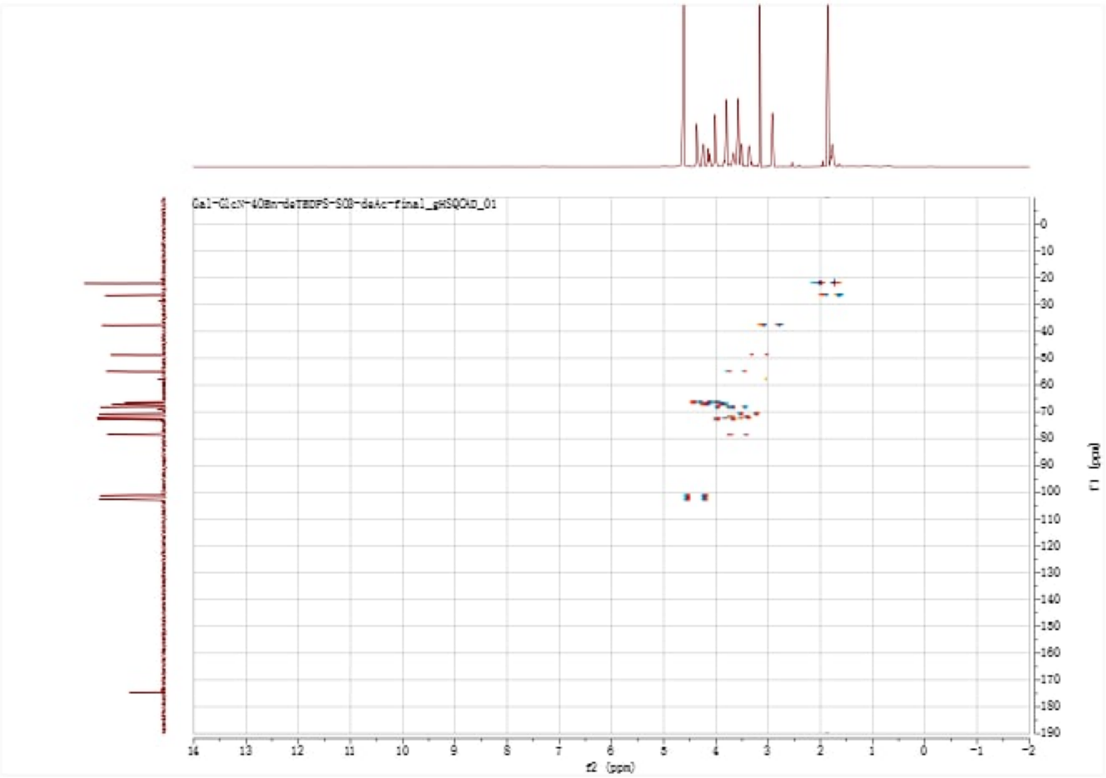

gHMBC(CDCl$_3$, 500 MHz) of **17**

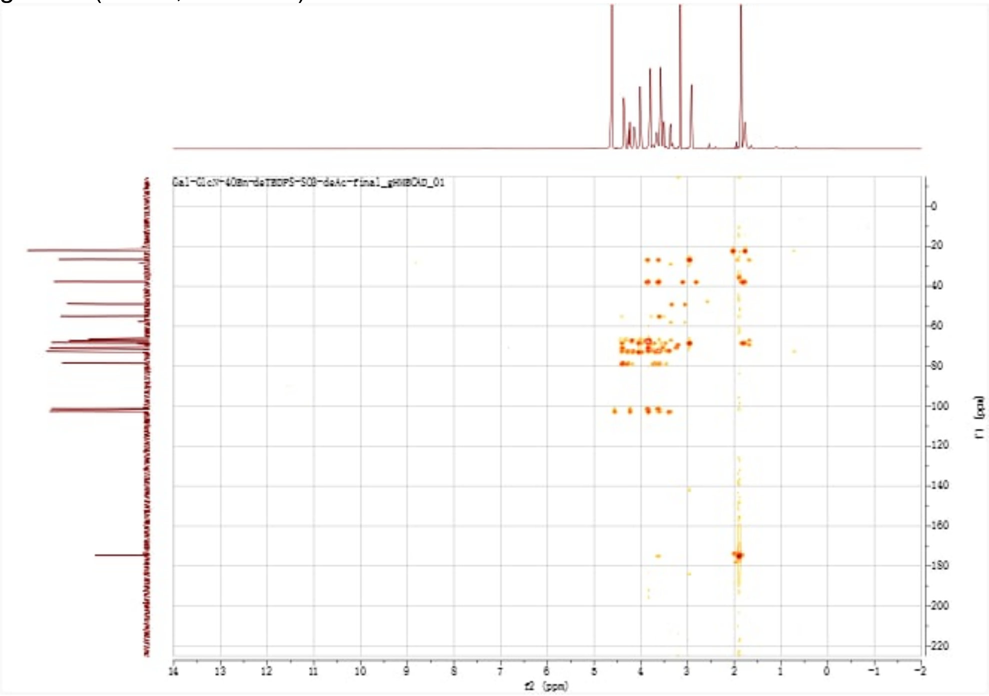

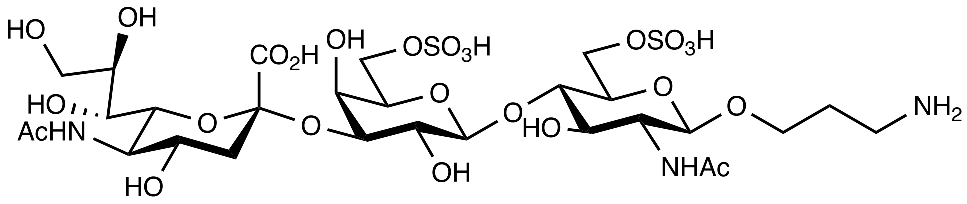


**18**

[1]H NMR ($D_2O$, v700 MHz) of **18**

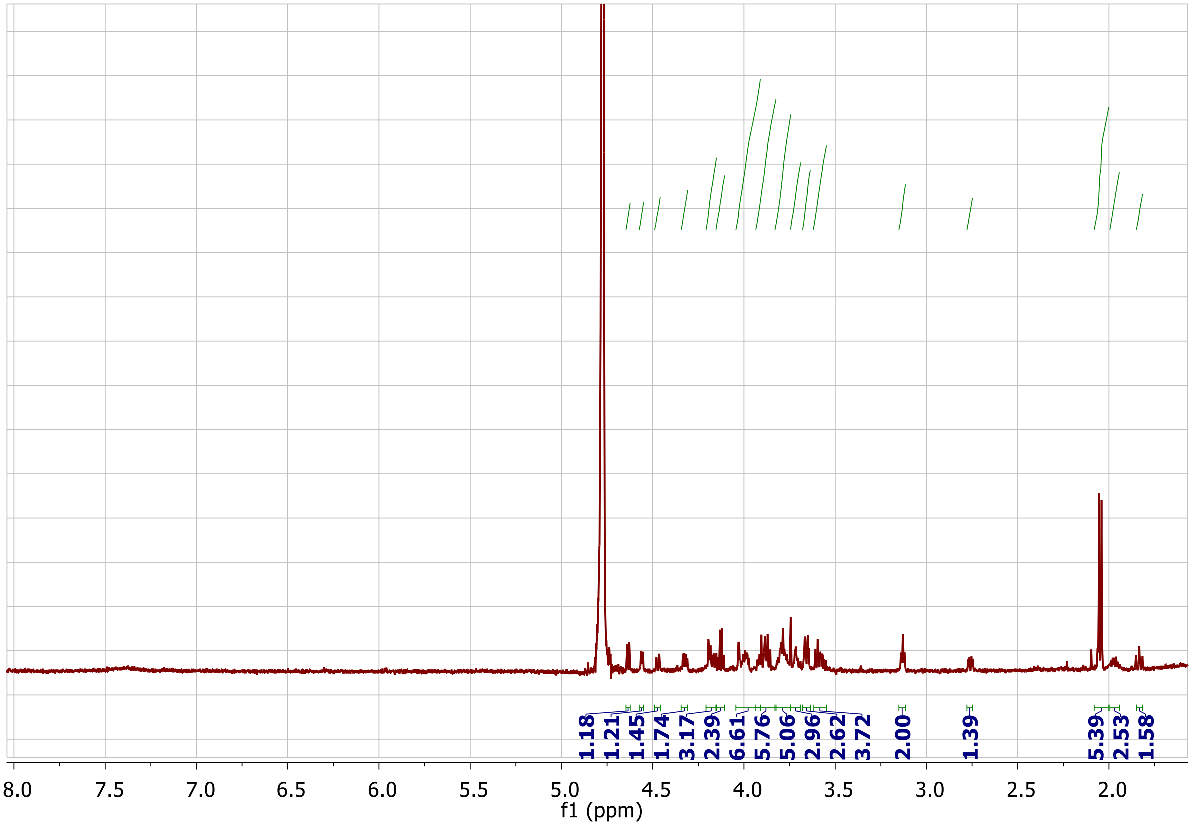
